# Direct 3D isolation in gelatin methacryloyl microbeads versus conventional 2D isolation and culture: a trade-off between mesenchymal stromal cell purity and niche-associated transcriptional profiles

**DOI:** 10.64898/2026.10.10.758039

**Authors:** Nali Hussein, Cornelius Miller, Yvonne Roger, Letizia Venturini, Michael Stadler, Marco Haertlé, Antonina Lavrentieva, Nadin Haase, Sophia Rudorf, Andrea Hoffmann, Dominik Egger

## Abstract

Bone-marrow mesenchymal stromal cell (MSC) isolation typically couples plastic-adherence selection to planar expansion. Therefore, the recovered cells may reflect culture adaptation as much as native stromal biology. Direct isolation within a three-dimensional matrix could reduce planar selection, but its effects on cellular recovery, composition and MSC transcriptional profiles remain insufficiently resolved. We therefore developed a gelatin methacryloyl (GelMA) workflow progressing from bulk hydrogels to medium-surrounded GelMA microbeads (GMBs) and compared 3D-derived cells with conventional 2D cultures. Qualitative live/dead staining with calcein acetoxymethyl ester (CAM) and propidium iodide (PI) and 5-ethynyl-2′-deoxyuridine (EdU) imaging established cellular activity in both GelMA formats. In bulk GelMA cultures from four human donors, CD73-, CD90- and CD105-associated flow-cytometric readouts did not increase from day 7 to day 21. At day 21, the combined marker readout was numerically higher in donor-matched 2D cultures than in bulk 3D cultures, which retained heterogeneous marker-defined populations. GMBs enabled cell recovery and flow-cytometric characterization and motivated an exploratory one-donor single-cell RNA sequencing (scRNA-seq) analysis. The final dataset contained 37,784 cells across four samples and 23 annotated populations. MSC-focused re-embedding resolved 13,175 cells into two 3D and four 2D clusters. A pooled comparison identified 4,840 differentially expressed genes (DEGs, false discovery rate [FDR]<0.01, |log2FC|≥1), indicating a broad method-associated expression shift. The 3D C1 cluster combined progenitor-, extracellular-matrix-, adhesion-, cytokine-, hypoxia/glycolysis- and predicted-interaction-associated programmes, whereas 2D clusters showed mitochondrial, cytoskeletal/mechanoadaptive and proliferative programmes. Thus, 2D culture yielded a more strongly represented and compositionally purer MSC compartment, while 3D isolation contained rare MSC clusters with niche-associated transcriptional profiles within a heterogeneous marrow environment. These findings support a testable native-associated tendency for 3D, but donor replication, matched recovery and direct native-reference mapping are required before inferring greater in-vivo similarity or functional superiority.

## Introduction

Mesenchymal stromal cells (MSCs) are commonly isolated from human bone marrow by adherence to tissue-culture plastic and subsequent expansion. This strategy efficiently enriches adherent stromal cells and underlies the widely used minimal phenotypic definition of culture-expanded MSCs [1], but also selects for attachment, survival and proliferation on a rigid planar substrate [2, 3, 4]. Comparisons of primary and cultured stromal cells have shown reduced heterogeneity and the emergence of fibroblast-like, proliferative and metabolically adapted states during culture [2, 3, 4, 5, 6]. Although these changes can support reliable expansion, cell numbers and marker enrichment alone do not establish how closely the recovered cells represent the stromal states present in bone marrow.

Native bone-marrow stroma comprises rare, heterogeneous non-haematopoietic populations involved in extracellular matrix (ECM) organization, vascular and perivascular structure, and haematopoietic niche support. Single-cell studies of freshly isolated adult and fetal human marrow have identified progenitor-like, perivascular, matrix-remodelling and haematopoiesis-supporting stromal states [4, 7, 8, 9]. Preferential expansion of one state may therefore obscure less abundant niche-associated populations. Canonical MSC-associated markers such as CD73, CD90 and CD105, together with CD271/NGFR, LEPR, PDGFRB and MCAM, provide complementary phenotypic information, but no single antigen or transcript establishes native identity, self-renewal or functional potency [1, 4, 7, 8, 9]. Assessing native-associated features requires coordinated evidence across stromal identity, matrix and adhesion programmes, metabolism, cytokine responsiveness and interaction potential, followed by direct reference comparison and functional validation.

Three-dimensional matrices offer an alternative to recovery through planar plastic adherence. Gelatin methacryloyl (GelMA) combines photocrosslinkable network formation with cell-interactive and protease-remodellable motifs. Encapsulation changes cell shape, adhesion, mechanical loading and matrix accessibility, creating a different selection and adaptation environment from rigid plastic [10, 11, 12]. In some stromal-cell systems, 3D culture retains stronger MSC- or osteolineage-associated signatures, whereas conventional propagation favours more homogeneous fibroblast-like states [3]. However, matrix composition, mechanics, local gradients and enzymatic digestion can themselves affect cell behaviour and recovery. A 3D-associated transcriptional profile may consequently reflect retention, selective survival, induced adaptation or preferential recovery rather than preservation of a native state.

We initially evaluated bulk GelMA with primary adipose-derived MSCs and heterogeneous bone-marrow mononuclear cells (BM-MNCs). The format allowed qualitative assessment of cellular activity, DNA synthesis and marker-defined recovery, but longer transport paths, restricted optical access and the hydrogel volume requiring digestion motivated its miniaturization. A GelMA microbead (GMB) workflow was therefore adapted from millifluidic encapsulation concepts [13, 14]. Approximately 500-µm, medium-surrounded beads were intended to shorten transport paths, increase surface area, facilitate imaging and enzymatic recovery, and permit sampling of one bone-marrow preparation across time points and analyses. These geometric advantages provided the rationale for experimental evaluation rather than evidence of improved transport.

It remains unclear whether cells with MSC-like phenotypes can be recovered directly from heterogeneous human bone marrow in a 3D GelMA environment and whether the resulting cell populations differ systematically from those selected by conventional 2D isolation and culture. In particular, reduced MSC representation after 3D isolation could reflect lower recovery alone or the presence of rare niche-associated transcriptional profiles that are lost, obscured or remodelled during plastic-adherent selection. Distinguishing these possibilities requires comparison of cellular composition and transcriptional profiles, not marker enrichment alone.

Here, we developed a workflow from bulk GelMA to medium-surrounded GMBs and compared the cellular composition, MSC heterogeneity and transcriptional programmes associated with 3D and conventional 2D isolation. We hypothesized that 2D culture would generate a more strongly enriched, culture-adapted MSC compartment, whereas 3D isolation would contain fewer MSCs but retain or induce distinct matrix-, progenitor- and niche-response-associated transcriptional profiles. The exploratory single-cell comparison addressed these method-associated transcriptional profiles while distinguishing representation in the final dataset from absolute recovery. Native similarity was not inferred without direct comparison to freshly isolated references.

## Materials and Methods

### Human material and culture conditions

Primary adipose-derived mesenchymal stromal cells (adMSCs) were isolated in-house from human adipose tissue, cryopreserved and encapsulated at passage 3 after thawing. The use of human adipose tissue was approved by the Ethics Committee of the University of Lübeck (approval numbers 20-233, November 2020, and 2024-455_1, November 2024). Written informed consent was obtained from all adipose-tissue donors. Cryopreservation medium contained 80% α-MEM, 10% human platelet lysate and 10% DMSO (v/v).

Fresh marrow obtained from surgically removed femoral heads was used for bulk cultures and the GMB flow-cytometry pilot experiment. Bulk flow cytometry comprised four donor-matched preparations. The GMB pilot involved an independent additional donor. Fresh iliac-crest aspirate from one healthy volunteer was used for single-cell RNA sequencing (scRNA-seq). The study was conducted in accordance with the Declaration of Helsinki and approved by the Institutional Ethics Committee of Hannover Medical School (565-2009 [28 August 2009], 565-2016 [22 April 2016] to A.H. for the use of cells from bone marrow from the bone (femoral head) of human patients as well as 2858-2015 [25 May 2021] to A.H. for the use of cells from bone marrow from the iliac crest of human voluntary donors). Written informed consent was obtained from all bone-marrow donors.

All cell cultures were maintained statically at 37 °C, 5% CO₂ and 5% O₂ in α-MEM (Gibco, Thermo Fisher Scientific) supplemented with 2.5% (v/v) human platelet lysate (PL BioScience) and 0.5% (v/v) gentamicin stock solution (Sigma-Aldrich). Approximately half the medium was replaced twice weekly. Trypsin was used for the first detachment of primary, conventionally isolated bone-marrow MSCs. Subsequent passages and adMSC culture used Accutase (BioLegend).

### GelMA preparation and characterization

Gelatin methacryloyl (GelMA) was prepared by reacting gelatin with methacrylic anhydride (MAA), followed by dialysis and lyophilization [11, 15]. The terms low, medium and high degree of functionalization (DoF) identify formulations rather than nominal functionalization percentages. For bulk cultures, equal volumes of 5% BATCH1 low-DoF and 5% BATCH1 high-DoF GelMA were mixed to yield 5% (w/v) medium-DoF GelMA with 0.2% (w/v) lithium phenyl-2,4,6-trimethylbenzoylphosphinate (LAP, Tokyo Chemical Industry). GMBs were prepared from 5% (w/v) BATCH2 high-DoF GelMA with 0.4% (w/v) LAP.

Functionalization was determined by ¹H-NMR in D₂O at 313 K against unmodified gelatin from the same source batch. Storage (G′) and loss (G″) moduli were measured for cell-free 5% (w/v) BATCH1 medium- and high-DoF and BATCH2 high-DoF formulations containing 0.2% (w/v) LAP (Supplementary Table S2). Synthesis, NMR and rheology are described in Supplementary Methods SM1–SM3.

### Bulk GelMA and conventional 2D culture

Bone marrow was diluted with three volumes of PBS and passed through a 100-µm cell strainer (BD, 352360). Up to 30 ml of diluted material was layered onto 15 ml Histopaque-1077 (Sigma-Aldrich, H8889) and centrifuged at 500 × g for 30 min at room temperature without braking. The MNC interface was collected, diluted in PBS, washed at 500 × g for 5 min and resuspended in culture medium. No erythrocyte lysis was performed for the bulk/2D flow-cytometry experiments.

BM-MNCs were encapsulated without prior MSC expansion at 5 × 10⁶ cells/ml GelMA. For bulk flow cytometry, separate pairs of wells were prepared for each donor and time point. Each well of a 6-well plate contained 2 ml GelMA overlaid with 4 ml culture medium, and the two corresponding wells were pooled for analysis. Photocrosslinking was performed for 3.5 min at 405 nm and 9 mW/cm² (Luna Crosslinker, Gelomics). Donor-matched 2D controls were seeded in T75 flasks at 5 × 10⁵ MNCs/cm². Bulk samples were analysed on days 7 and 21 and 2D samples on day 21. Donors 1, 2 and 4 remained at passage 0, whereas Donor 3 was passaged once and analysed at passage 1.

Primary adMSCs were encapsulated at 1 × 10⁶ cells/ml GelMA for all bulk and GMB imaging experiments. Initial validation used calcein acetoxymethyl ester (CAM)/propidium iodide (PI) and 5-ethynyl-2′-deoxyuridine (EdU)/4′,6-diamidino-2-phenylindole (DAPI) microscopy. BM-MNC cultures were assessed by brightfield and fluorescence microscopy and flow cytometry.

### GMB generation and culture

After density-gradient isolation, MNCs for the GMB flow-cytometry pilot were treated with 10 ml freshly prepared erythrocyte-lysis buffer (8.3 g NH₄Cl, 1 g KHCO₃ and 1.8 ml of 5% EDTA solution per litre) for 3 min at room temperature. Following addition of 20 ml PBS, cells were centrifuged at 500 × g for 3 min at 4 °C and washed twice with PBS. This lysis step was not used for bulk encapsulation.

A millifluidic encapsulation approach [13] was adapted for GMB generation. A Legato dual-syringe pump (Harvard Bioscience) delivered cell-containing GelMA at 100 µl/min through a Sterican needle (20 G, 0.90 × 70 mm, B. Braun) into white mineral oil (Sigma-Aldrich, 330779-1L) flowing at 4.2 ml/min. Both phases passed through transparent tubing with a 1.6-mm inner diameter (Pieper Filter) and were photocrosslinked for approximately 55 s at 405 nm and 9 mW/cm² (Luna Crosslinker, Gelomics), with the GelMA feed maintained at approximately 37 °C. After oil separation, beads were washed twice with 20 ml PBS and once with 10 ml medium. Their diameter was approximately 500 µm, estimated from calibrated images. Approximately 0.5–1 ml beads per well of a 6-well plate were cultured statically under approximately 5 ml medium.

All BM-MNC GMB experiments used 5 × 10⁶ MNCs/ml GelMA. adMSC-loaded GMBs were used for imaging validation, and BM-MNC-loaded GMBs were assessed by microscopy and day-21 flow cytometry. Separate day-7 and day-21 GMB preparations were processed for scRNA-seq.

### Staining and image analysis

CAM/PI staining used 5 µl each of calcein-AM stock (5 mg/ml) and propidium iodide stock (1 mg/ml, Sigma-Aldrich) per ml PBS. Bulk gels and GMBs were incubated with approximately 1 ml staining solution per ml GelMA for 1 h at 37 °C in the dark and washed twice with PBS. EdU was detected after 24-h pulses at 10 µM (EdU Click Kit 488, BCK-EDU488, supplied by Sigma-Aldrich). Samples were fixed with 3.7% formaldehyde for 15 min, permeabilized with 0.5% Triton X-100 for 20 min and subjected to fluorescent click labelling for 30 min. DAPI counterstaining used 3 µl of a 1 mg/ml stock per ml staining solution. The GMB EdU series comprised the displayed days 0, 3, 7, 14 and 21. For D0, EdU was added immediately after encapsulation and fixation followed 24 h later.

Bulk fluorescence images were acquired using a Leica DM IL LED Fluo IVD with LAS X. Dry 4×/0.10 and 10×/0.32 objectives were used. GMB fluorescence imaging used a Nikon Eclipse TE2000-E. Confocal imaging used a C1 scanner operated with EZ-C1 Gold 3.80, with a 4×/0.20 objective for the adMSC EdU series and a 10×/0.30 objective for the endpoint BM-MNC series. The latter was acquired at 2.5-µm z intervals. Representative optical sections were selected at approximately 0%, 25%, 50%, 75% and 100% of the stack index range. Brightfield imaging used an Axiovert.A1 (Carl Zeiss Microscopy). GMB EdU analysis used Fiji/ImageJ 1.54p. Individual beads were outlined as ROIs, channels were separated and converted to 8-bit images, and thresholding and watershed segmentation were followed by Analyze Particles with a 1–600-pixel² size filter. The EdU fraction was calculated as 100 × EdU-positive/DAPI-positive signals per bead. One bead preparation was divided among time points, and individual beads were technical subsamples. Bulk EdU and CAM/PI images were assessed qualitatively without calculation of a quantitative viability rate.

### Cell recovery

Hanks’ balanced salt solution (HBSS, Sigma-Aldrich, H1387) was supplemented with sodium bicarbonate powder (NaHCO₃, 0.35 g/L). For flow cytometry, GMBs were washed with HBSS and digested with an equal volume of 0.1% (w/v) Clostridium histolyticum collagenase (Sigma-Aldrich, C2674, ≥125 CDU/mg) in HBSS for approximately 15 min at 37 °C. Bulk gels were disrupted mechanically with a spatula and digested with twice the gel volume of 0.2% collagenase solution in HBSS for approximately 60 min in the incubator. PBS dilution and centrifugation at 300 × g for 5 min were followed by 10 min of Accutase treatment. Suspensions were diluted with medium, passed through a 40-µm cell strainer, centrifuged again at 300 × g for 5 min and resuspended in flow-cytometry buffer.

### Flow cytometry

Antibody staining used the panels in Supplementary Tables S3A–E (bulk) and S5A–D (GMB) for 10–15 min on ice in the dark. Staining reactions contained 100 µl antibody solution. Washes used MACS Running Buffer with 0.5% BSA and centrifugation at 200 × g for 5 min. Samples were acquired using a MACSQuant Analyzer 10 (Miltenyi Biotec) and analysed in FlowJo 10.10.0. Floreada was additionally used to display the GMB histograms in Supplementary Figure S4. Isotype and unstained controls were included.

Percentages refer to singlets in the respective separately acquired panel, even where the target population was defined by additional upstream gates. The separately acquired panels therefore have no common denominator. Their partly overlapping readouts must not be summed into a cellular composition. In GMB samples, the stromal enrichment fraction (SEF) was defined by combined negativity for CD3, CD14, CD235a, CD34, CD56, CD16, CD11b, CD19, CD45 and HLA-DR, measured in three exclusion channels. MSC markers were evaluated within this operational fraction. Surface markers and cocktail gates were interpreted as phenotypic readouts rather than definitive evidence of cell identity.

### Single-cell sampling and sequencing

Four samples from one bone-marrow donor were examined: uncultured D0, GMB cultures on days 7 and 21, and conventional 2D culture on day 21. Following density-gradient isolation, D0 MNCs were enriched for CD45⁻/CD235a⁻ cells by fluorescence-activated cell sorting using a BD FACSAria Fusion cell sorter (BD Biosciences) at the Central Research Facility Cell Sorting, Hannover Medical School. For the 3D sequencing samples, erythrocytes were lysed before encapsulation for 2 min using a commercial reagent (Red Blood Cell Lysis Solution, diluted from the supplied 10× stock to the manufacturer-recommended 1× working solution, Miltenyi Biotec, 130-094-183). Preparations were diluted with 20 mL PBS, centrifuged at 500 × g for 3 min at 4 °C and washed twice with PBS. The 3D samples did not undergo the additional CD45/CD235a depletion applied to D0.

For sequencing, GMBs were digested with 0.1% collagenase (C2674) in the HBSS formulation described above for 15–30 min at 37 °C, 5% CO₂ and atmospheric oxygen. PBS dilution and centrifugation at 300 × g for 5 min were followed by Accutase treatment for 10–15 min. The 2D cells were detached with trypsin-EDTA (Sigma-Aldrich, T3924-500ML) for 5 min at 37 °C and resuspended in medium. Following counting and 7-AAD staining, 7-AAD-negative cells were sorted into FCS-precoated tubes, with a sorting target of 50,000 cells per sample. After a final cell count, the numbers loaded were 27,000 for D0, 20,000 for 3D D7, 15,970 for 2D D21 and 20,000 for 3D D21. D0 viability was recorded qualitatively as good. Recorded viabilities were 91% for 3D D7, 98% for 2D D21 and 95% for 3D D21. Supplementary Table S6 distinguishes cell loading and viability from the final analysed cell counts. The primary MSC comparison used the 3D and 2D day-21 samples.

### Single-cell analysis

Single-cell libraries were prepared using droplet-based barcoding [16] on a Chromium X instrument (10x Genomics) with Chromium GEM-X Single Cell 3′ Reagent Kits v4 according to user guide CG000731 Rev A. Library fragment sizes and concentrations were assessed using the Agilent Bioanalyzer High Sensitivity DNA Assay and Qubit dsDNA HS Assay, respectively. Libraries were sequenced at Hannover Medical School on an Illumina NovaSeq X Plus with the NovaSeq X 10B Reagent Kit (100 cycles), loaded at 180 pM with 1% PhiX. Read lengths were 28 bp for Read 1, 90 bp for Read 2 and 10 bp for each index read. Mean sequencing depths ranged from 23,521 to 75,316 reads per cell (Supplementary Table S6B). Total read counts and Cell Ranger cell estimates are reported separately in Supplementary Table S6C.

Data analysis comprised two parts. The first part generated a quality-controlled, annotated dataset and consisted of (i) generation of a corrected count matrix, (ii) quality control (QC) with delayed filtering, which allowed cell clusters to be evaluated before any cells were removed, (iii) reference-based cell type prediction, and (iv) manual cell type annotation integrating these predictions with marker gene evidence. The second part used this dataset for (v) differential expression analysis between two-dimensional (2D) and three-dimensional (3D) mesenchymal stromal cell (MSC) cultures, (vi) gene set enrichment analysis, (vii) cell–cell communication analysis, and (viii) trajectory analysis. See Supplementary Methods SM4 for full details.

### Statistical analysis

The donor was the biological experimental unit. Bulk flow-cytometry results were displayed as individual donor values and mean ± SD, with lines connecting paired measurements. Two-sided paired t-tests were performed in GraphPad Prism 8.0.1. Raw p values were subsequently adjusted by the Holm procedure (Supplementary Table S4), separately for eleven D7–D21 time-course comparisons and twelve 2D–3D day-21 comparisons. Only complete pairs were included: four per comparison, except for the CD3 day-21 endpoint comparison (D1, D3 and D4, n = 3), for which the 2D value of D2 was missing. Missing values were not replaced with zero. Significance annotations use Holm-adjusted p values <0.05.

The GMB pilot experiment was described without inferential statistics. The scRNA-seq analysis was exploratory and involved one donor. P values, FDR estimates and GSEA q values were used for within-dataset ranking rather than donor-level inference. NMR processing ranges indicate technical sensitivity rather than biological SD. Exported zero permutation p values or q values were not interpreted as exact zero probabilities.

## Results

### Bulk GelMA supports cellular activity without enrichment of MSC-associated phenotypes

Primary adipose-derived MSCs were first assessed qualitatively in bulk GelMA with a medium degree of functionalization (DoF) using CAM/PI and EdU/DAPI imaging. CAM-positive cells, classified as viable, remained detectable from day 3 through day 21 (Figure 1A). Cell spreading developed gradually, with more extensive cell–cell contacts apparent from approximately day 14. In deeper hydrogel fields, CAM signal was reduced and PI-positive cells, classified as dead, became more prominent at later time points (Supplementary Figure S2). Among the displayed images, EdU-positive nuclei were clearly detectable only at day 3 and were concentrated near the hydrogel periphery, indicating early, surface-proximal DNA synthesis during the labelling pulse (Figure 1A). These observations established the feasibility of supporting cellular activity in bulk GelMA and prompted evaluation of heterogeneous bone-marrow input.

**Figure 1.**
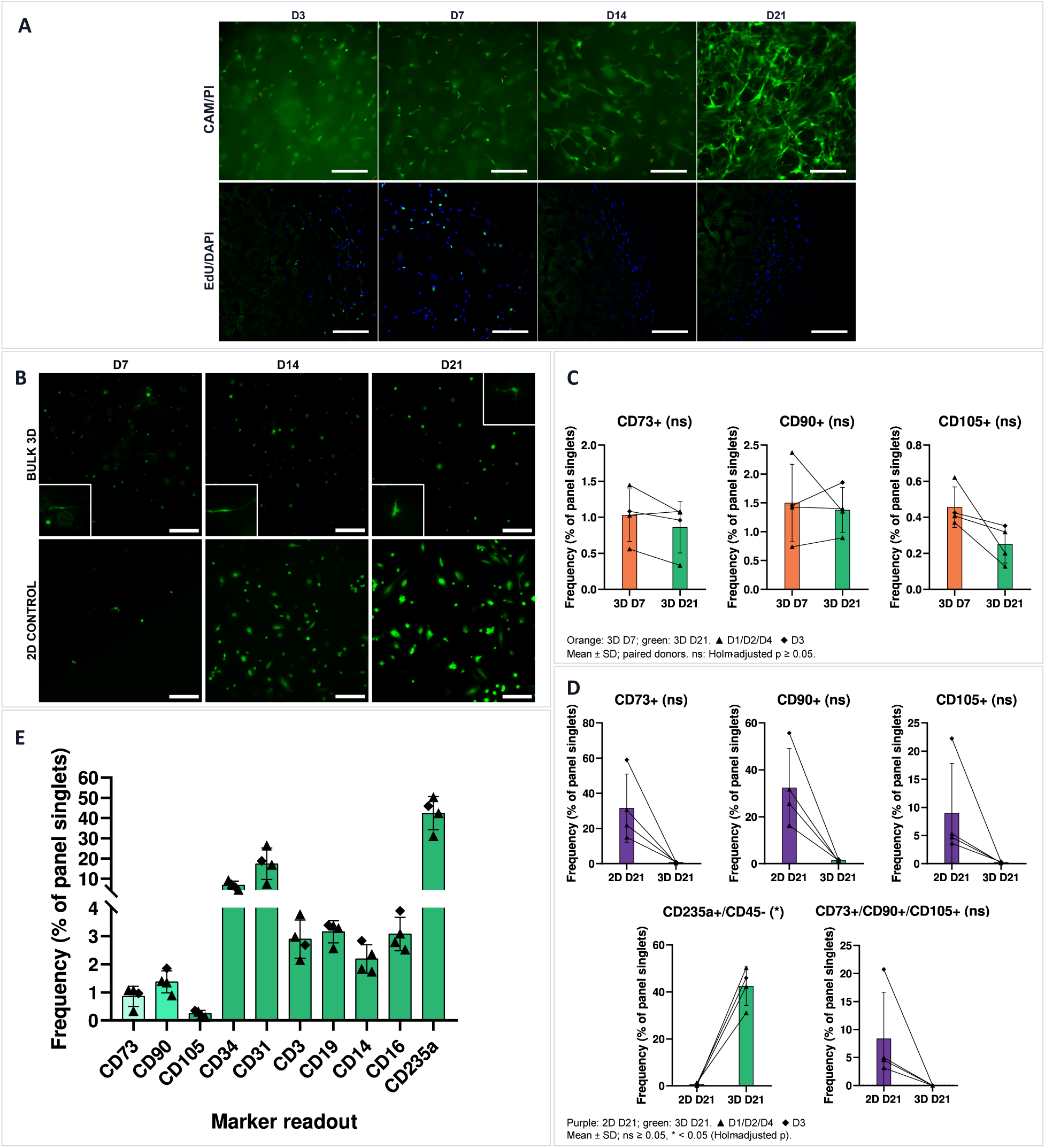
Bulk-GelMA validation with primary adMSCs and bone-marrow-derived cells. (A) CAM/PI (CAM, green and PI, red) and EdU/DAPI (EdU, green and DAPI, blue) images of primary adMSCs in medium-DoF bulk GelMA at the displayed culture days. Scale bars represent 200 µm. (B) BM-MNC CAM/PI time series in bulk GelMA and the displayed 2D control. White scale bars represent 200 µm. Insets show elongated cells at approximately twofold magnification relative to the main images. Each square inset represents an area of approximately 175 × 175 µm. (C) Paired 3D day-7 and day-21 MSC-marker readouts. (D) Paired day-21 2D and 3D readouts, including the combined CD73/CD90/CD105 readout. (E) Marker-frequency profile of bulk 3D day 21. Bars show mean ± SD and black symbols show donors. D1, D2 and D4 are triangles, and D3 is a diamond. Lines in C and D connect the same donor. The 2D cultures of D1, D2 and D4 remained at P0, whereas D3 was analysed at P1. Comparisons use two-sided paired t-tests with separate Holm correction for the eleven temporal and twelve day-21 comparisons. Significance symbols refer to Holm-adjusted p values, with ns denoting p ≥ 0.05 and * denoting p < 0.05. The displayed comparisons in C and D include four paired donors. The condition overview in E uses all available observations. Frequencies use the singlet denominator of the respective acquisition panel and are not mutually exclusive population proportions. Exact summaries and p values are provided in Supplementary Table S4.

BM-MNC cultures contained predominantly rounded cells with occasional adherent elongated or spindle-shaped cells (Figure 1B). CAM-positive structures remained detectable at days 7, 14 and 21, but the qualitative image series did not show progressive enrichment or expansion of spindle-shaped cells. Flow cytometry was therefore used to assess marker-defined populations that could not be resolved by morphology, comparing 3D days 7 and 21 and the day 21 endpoints of 3D and conventional 2D culture.

In paired 3D cultures from four donors, none of the measured MSC-associated readouts changed significantly between days 7 and 21 after Holm correction (Figure 1C). Mean individual CD73-, CD90- and CD105-positive frequencies remained below 2%, and CD73/CD90/CD105 triple-positive events were practically undetectable. Extending bulk culture to day 21 therefore did not increase the measured MSC-associated frequencies. Complete donor-paired summaries, raw p values and Holm-adjusted p values are provided in Supplementary Table S4. Acquisition panels, gating sequences and denominators are documented in Supplementary Table S3, with expanded flow-cytometric views in Supplementary Figure S3.

At day 21, the individual CD73⁺, CD90⁺ and CD105⁺ fractions and the combined CD73⁺/CD90⁺/CD105⁺ readout after exclusion gating were numerically more frequent in 2D than in bulk 3D culture (approximately 8–32% versus 0.007–1.38%, Figure 1D, Supplementary Table S4), without statistical significance after Holm correction. Donors 1, 2 and 4 remained at passage 0, whereas Donor 3 was analysed at passage 1. Passaging may have contributed to its deviation from the other 2D samples, although its 2D-versus-3D contrast followed the same direction. The higher MSC-associated frequencies in 2D were compatible with selection during plastic-adherent culture.

CD14⁺/CD45⁺ events (monocyte/macrophage-associated fraction), CD16⁺/CD45⁺ events (CD16-expressing myeloid-associated fraction) and total CD31⁺ events (CD31-associated fraction) were likewise numerically more frequent in 2D at day 21 but did not meet the Holm-adjusted significance threshold. Conversely, CD235a⁺/CD45⁻ events (erythroid-associated fraction) were more frequent in bulk 3D than in 2D at day 21 (42.43 ± 8.21% versus 0.68 ± 0.57%, Holm-adjusted p = 0.0286) and did not change significantly between 3D days 7 and 21 (Figure 1D–E, Supplementary Figure S3 and Supplementary Table S4). Frequencies from separately acquired antibody panels must not be summed into an overall cellular composition. Thus, bulk GelMA supported viability-associated signals and early DNA synthesis without enriching MSC-associated phenotypes under the conditions examined. Persistent erythroid-associated events and stronger PI-positive signals in deeper fields motivated evaluation of smaller, medium-surrounded GMBs.

### GelMA microbeads support three-dimensional bone-marrow culture and cell recovery

An established millifluidic encapsulation concept [13, 14] was adapted for 3D isolation, culture and recovery of bone-marrow-derived cells in GMBs (Figure 2A). The medium-surrounded format enabled modular sampling and facilitated optical and enzymatic access relative to the bulk hydrogel. Qualitative CAM/PI imaging showed viable adMSCs throughout the displayed culture period, without an obvious time-dependent accumulation of PI-positive cells. Cells rapidly adopted elongated or stellate morphologies and formed extensive cell–cell contacts (Figure 2B). EdU/DAPI imaging and image-derived EdU-positive/DAPI-positive fractions indicated early DNA synthesis, with only sparse EdU-positive nuclei from day 7 onwards (Figure 2B–C), consistent with proliferative activity concentrated in the initial culture phase.

**Figure 2.**
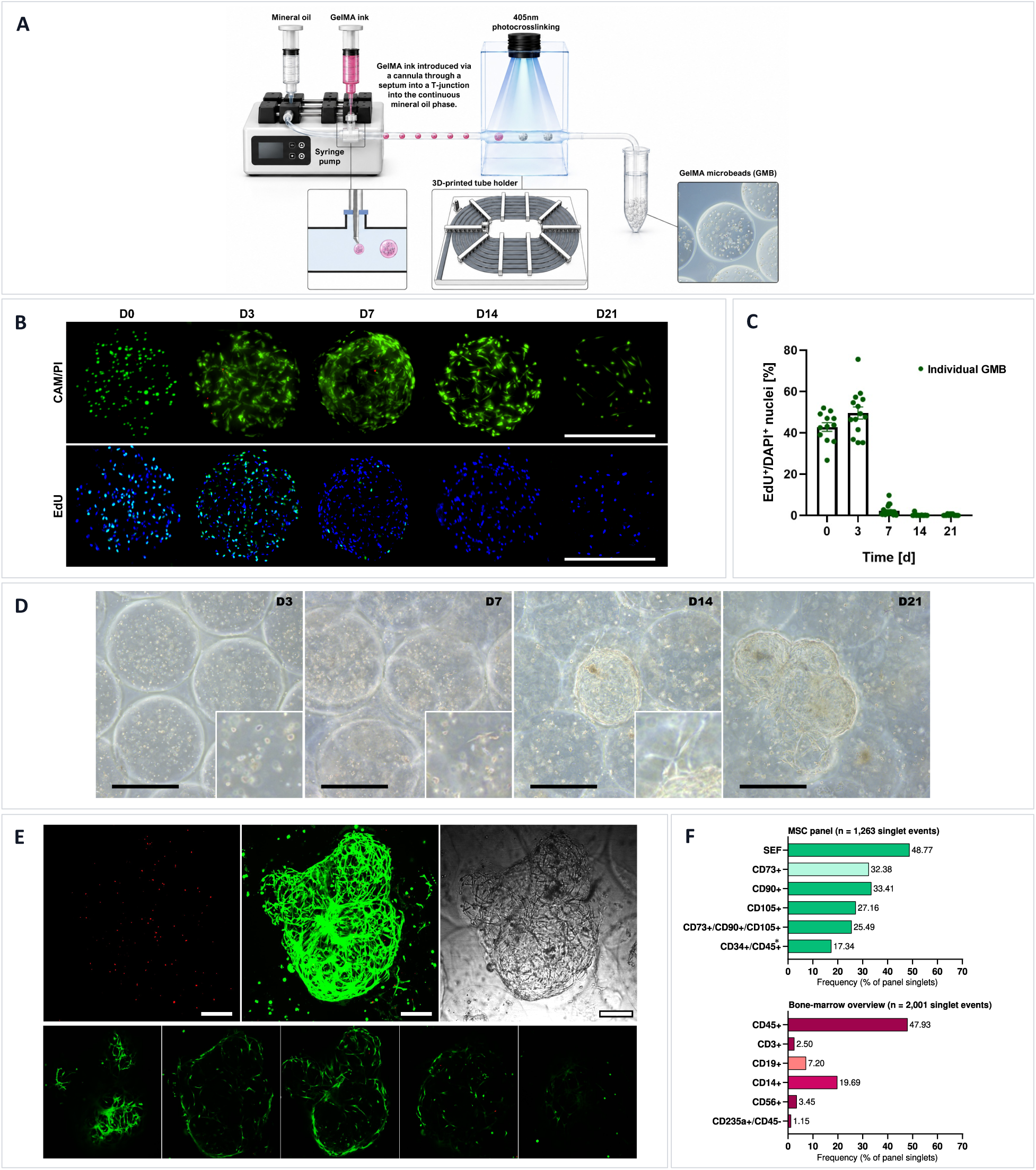
Generation and exploratory biological validation of GelMA microbeads. (A) Adapted millifluidic GMB generation. (B) Qualitative adMSC CAM/PI and EdU/DAPI time series. White scale bars represent 500 µm. (C) EdU-positive/DAPI-positive fractions in individual GMB images after a 24-h EdU pulse. Data are presented as mean ± SEM. Beads were technical subsamples of one preparation, not independent biological replicates. (D) Brightfield time series of BM-MNC-loaded GMBs, with insets showing spindle-shaped or stellate cells and bead-associated networks. Black scale bars represent 500 µm. Insets show approximately threefold magnification relative to the main images, with each square inset representing an area of approximately 215 × 215 µm. (E) Endpoint CAM/PI maximum projection, brightfield image and optical sections selected at approximately 0%, 25%, 50%, 75% and 100% of the stack index range. Scale bars represent 200 µm. (F) Day-21 single-donor flow-cytometry readouts. The MSC panel contained 1,263 singlets and the separate bone-marrow overview panel contained 2,001 singlets. SEF comprised 616 events and was defined by sequential exclusion of the FITC cocktail targeting CD3, CD14 and CD235a, the PE cocktail targeting CD11b, CD16, CD19, CD45, CD56 and HLA-DR, and PE-Vio770-conjugated CD34. The MSC-marker gates were evaluated within SEF but the displayed percentages use initial MSC-panel singlets as denominator. Triple-positive events numbered 322, equivalent to 25.49% of singlets and 52.27% of SEF. Overview-panel percentages use its separate singlet denominator. Bars are descriptive frequencies without error bars or inferential statistics. Representative fluorescence histograms and isotype controls are provided in Supplementary Figure S4. The panel definitions are detailed in Supplementary Table S5. *CD45 was not labelled individually but was included in a PE-labelled antibody cocktail.

BM-MNC-loaded GMBs initially contained predominantly rounded cells. Rare adherent spindle-shaped cells appeared by day 7, followed by numerous spindle- and stellate-like cells with cell–cell contacts at day 14 and larger, bead-spanning multicellular complexes at day 21 (Figure 2D). Endpoint confocal projections and optical sections detected CAM-positive cells within the beads, with higher apparent cell frequency and density near their boundaries (Figure 2E).

Exploratory single-donor flow cytometry at day 21 detected MSC-associated marker frequencies of approximately 25.5–33.4% among MSC-panel singlets: 32.38% for CD73, 33.41% for CD90, 27.16% for CD105 and 25.49% for CD73/CD90/CD105 triple positivity (Figure 2F). Representative fluorescence distributions and isotype controls are shown in Supplementary Figure S4. These readouts were descriptively higher than the corresponding bulk 3D values of 0.007–1.38%. The stromal enrichment fraction (SEF) was operationally defined by negativity for CD3, CD14, CD235a, CD11b, CD16, CD19, CD45, CD56, HLA-DR and CD34. Several immune-marker-defined fractions remained detectable in the separate overview panel, whereas CD235a⁺/CD45⁻ events accounted for only 1.15%, consistent with reduced erythroid carryover relative to bulk 3D culture. The triple-positive readout comprised 322 of the 616 SEF events, corresponding to 52.27% within SEF. The recovered population nevertheless remained heterogeneous. Because a limited surface-marker panel could not resolve the identity and diversity of its stromal-associated cells, scRNA-seq was used to compare 3D-derived and conventional 2D MSC transcriptional profiles.

### Single-cell profiling reveals distinct cellular compositions and MSC clusters

The exploratory scRNA-seq experiment comprised four samples from one healthy human bone-marrow donor: uncultured day 0 (D0), 3D day 7, 3D day 21 and conventional 2D day 21. D0 underwent density-gradient MNC isolation followed by CD45/CD235a depletion to enrich rare native stromal cells. However, no MSC-annotated population was identified in this sample, precluding its use as a direct native-MSC comparator. The 3D samples underwent isolation by density-gradient centrifugation and erythrocyte lysis without CD45/CD235a depletion. Cultured 2D and 3D samples were maintained at a nominal 5% O₂. Cell loading differed between samples (Supplementary Table S6). In addition, D0-specific selection precluded an unbiased longitudinal comparison of cellular composition. We therefore interpreted the 3D day-7 to day-21 comparison descriptively and focused the MSC analysis on the pooled day-21 3D-versus-2D contrast (Supplementary Table S6).

The final dataset contained 37,784 cells: 14,387 from D0, 7,389 from 3D day 7, 3,138 from 3D day 21 and 12,870 from 2D day 21. In the unintegrated joint UMAP, 2D day 21 occupied a comparatively discrete region, whereas D0 and the 3D samples extended across a broader, partly overlapping landscape (Figure 3A). Annotation resolved 23 populations spanning haematopoietic progenitors, myeloid, lymphoid and erythroid lineages, and six MSC-associated clusters (Figure 3B). All cells in the final 2D sample were MSC annotated, whereas the 3D day-21 sample contained rare MSC clusters within a heterogeneous bone-marrow-derived environment. These patterns reflect combined effects of isolation, culture, composition and potentially technical variation. UMAP separation is not a quantitative effect size. The design cannot distinguish retention of starting populations from differential survival, proliferation, induction of transcriptional profiles or recovery.

**Figure 3.**
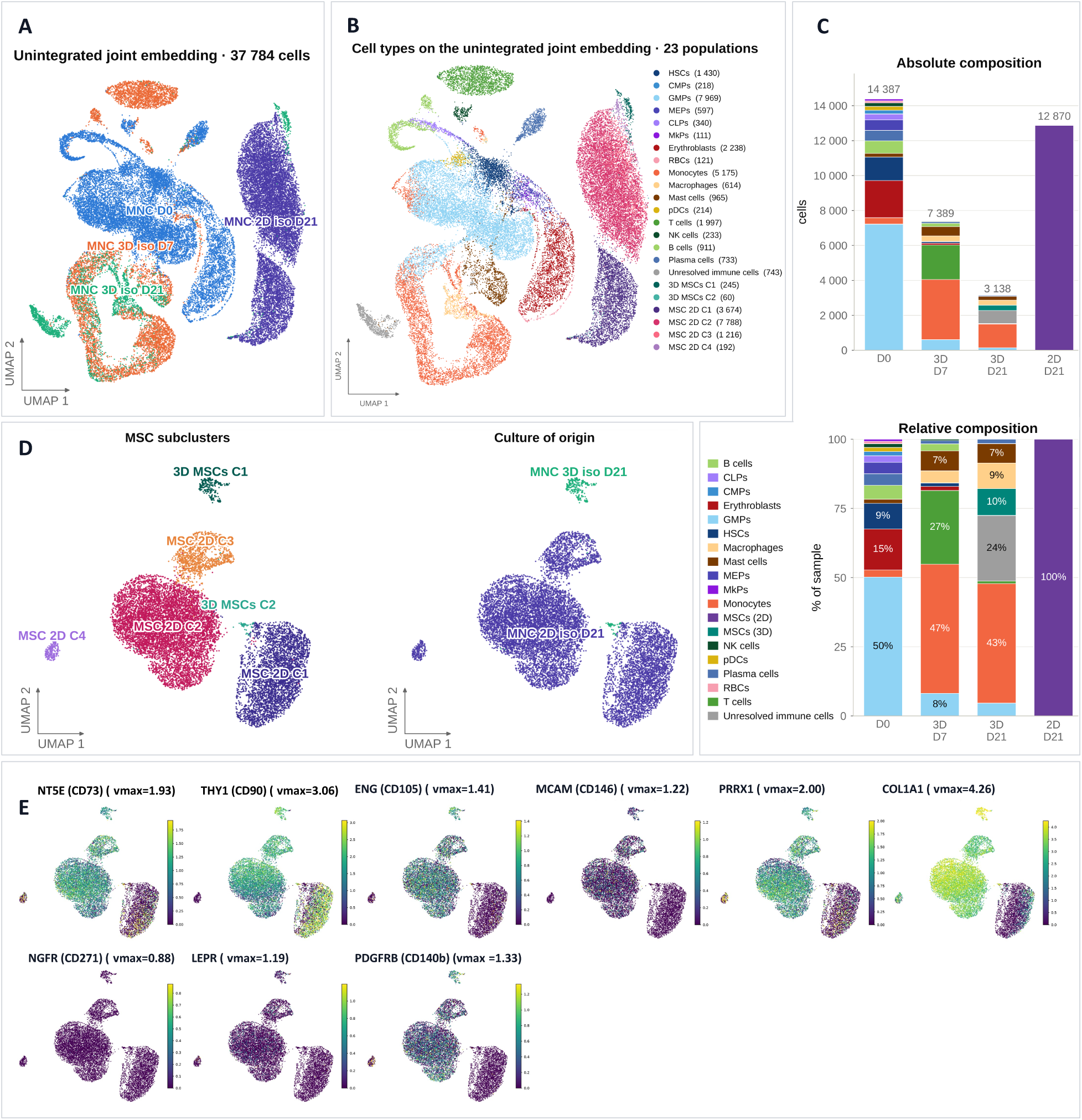
Single-cell landscape and MSC-associated heterogeneity. (A) Unintegrated joint UMAP coloured by condition (37,784 cells). (B) Annotation of 23 populations. (C) Absolute displayed cell counts and within-sample relative composition. D0 underwent CD45/CD235a depletion and is not an unbiased compositional baseline. (D) MSC-only re-embedding of 13,175 cells, resolving 3D C1/C2 and 2D C1–C4. UMAP proximity does not establish quantitative transcriptomic or functional similarity. (E) Feature UMAPs for NT5E, THY1, ENG, NGFR, LEPR, PDGFRB, MCAM, PRRX1 and COL1A1. For each gene, vmax was the 97.5th percentile of positive MSC expression and was identical across all six clusters. Scales are not directly comparable between genes. Sample-specific cell loading is reported in Supplementary Table S6. Final cell counts measure representation in this one-donor dataset rather than absolute isolation yield.

Within-sample frequencies showed a compositional shift in 3D cultures (Figure 3C). Monocytes represented 46.66% of cells at day 7 and 43.21% at day 21, whereas macrophages increased in relative frequency from 4.38% to 9.24%. haematopoietic stem cell (HSC)-annotated cells decreased from 1.25% to 0%, and granulocyte–monocyte progenitors (GMPs) from 8.17% to 4.62%. In the selectively processed D0 sample, the corresponding HSC and GMP fractions were 9.30% and 50.18%. The macrophage share of the monocyte/macrophage compartment increased from 8.59% to 17.62% between the two 3D time points. These relative changes were compatible with myeloid maturation while progenitor-associated cells remained detectable at day 21. They did not establish a specific differentiation route or absolute expansion of macrophages.

Re-embedding 13,175 MSC-annotated cells resolved two 3D and four 2D clusters: 3D C1 (245 cells), 3D C2 (60), 2D C1 (3,674), 2D C2 (7,788), 2D C3 (1,216) and 2D C4 (192) (Figure 3D). 3D C1 occupied a visually distinct region, while 3D C2 appeared adjacent to 2D C1. This proximity did not establish quantitative transcriptomic or functional similarity. The 12,870-versus-305 difference in 2D- and 3D-derived MSC counts demonstrated stronger 2D representation in the final dataset. Cell loading was 15,970 for 2D day 21 and 20,000 for 3D day 21. Composition, cell recovery, viability assessment, capture, transcript quality and QC attrition could all contribute, so these counts cannot be converted into absolute isolation yield without matched recovery measurements per defined marrow input.

Canonical and bone-marrow-associated marker transcripts further characterized the six clusters (Figure 3E). NT5E/CD73, THY1/CD90 and ENG/CD105 were heterogeneous, while PDGFRB, MCAM/CD146, PRRX1 and COL1A1 occupied substantial parts of the embedding. NGFR/CD271 and LEPR were sparse at the displayed thresholds. In the log-normalized matrix, 3D C1 expressed NT5E and THY1 in 97.1% of cells each, ENG in 95.5%, PDGFRB in 94.3%, MCAM in 44.9%, LEPR in 25.7%, PRRX1 in 97.1% and COL1A1 in 100%. These descriptive transcript-detection fractions are not protein measurements or donor-level estimates. NGFR/CD271 transcripts were detected in fewer than 0.1% of all MSC-annotated cells, limiting the usefulness of this marker for transcriptomic validation in the cultured dataset. Thus, coordinated stromal marker expression, rather than any single present or absent transcript, motivated the interpretive focus on 3D C1. The smaller 3D C2 cluster remained included in pooled analyses and cluster-resolved displays. Neither RNA detection nor its absence establishes functional MSC identity or protein negativity. Extended marker maps and detection profiles are provided in Supplementary Figure S6 and Supplementary Table S10.

### Three-dimensional and planar culture are associated with distinct MSC transcriptional programmes

To assess whether the isolation routes differed beyond selected markers, 3D C1/C2 were pooled and compared with 2D C1–C4. Of 15,444 tested genes, 4,840 met FDR <0.01 and |log2FC| ≥1: 2,735 were higher in 3D and 2,105 in 2D (Supplementary Table S7). The stricter FDR <0.001 threshold displayed in Figure 4A retained 4,285 genes, comprising 2,400 higher in 3D and 1,885 in 2D. This bilateral shift indicated broad transcriptional differences within the dataset. All cells originated from one donor and were randomly divided into three non-independent count-summed blocks per condition for exploratory dispersion estimation. Consequently, rep1– rep3 are not biological replicates, and model-derived p and FDR values do not estimate between-donor variation or support population-level inference. Effect directions and rankings were interpreted descriptively.

**Figure 4.**
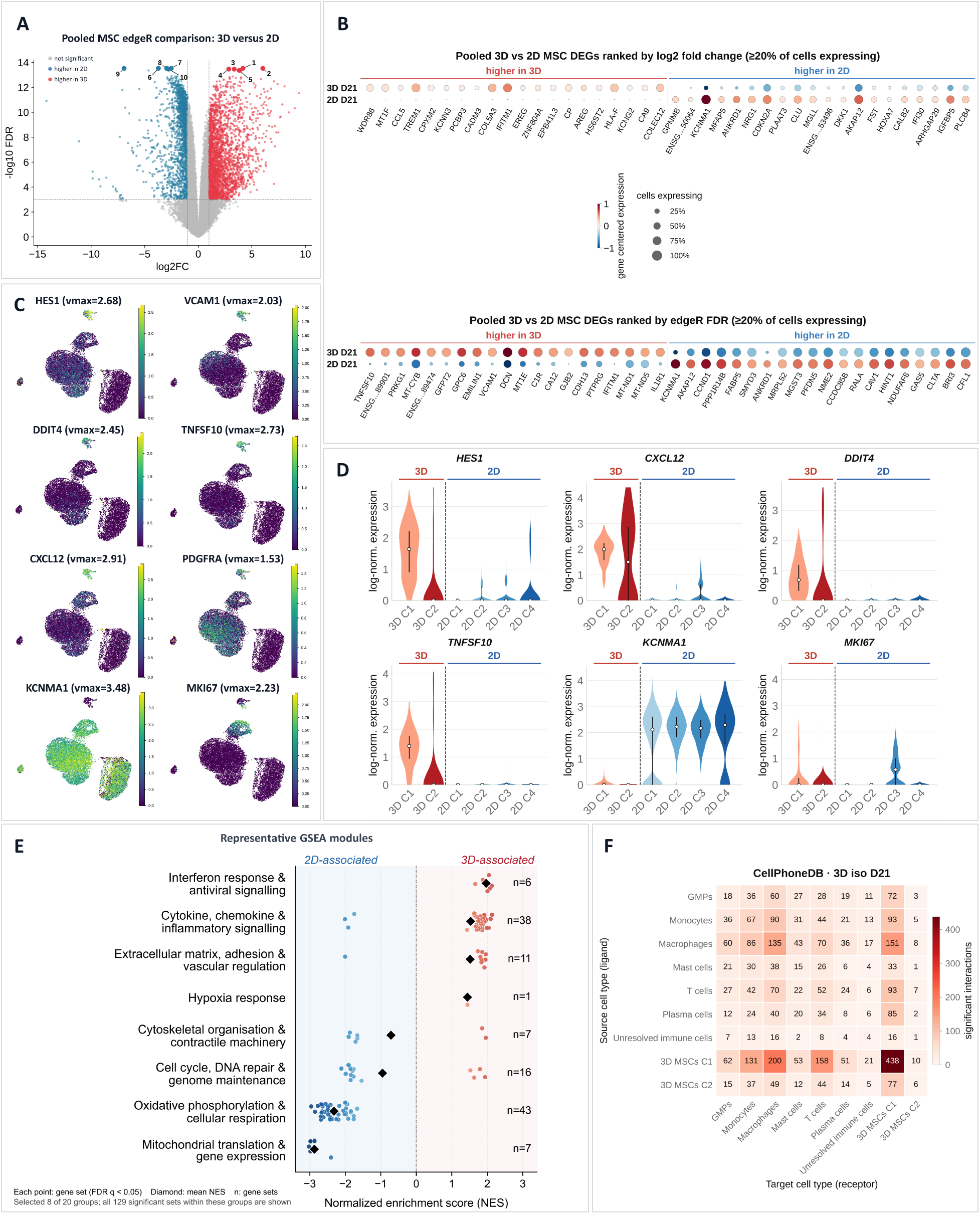
Transcriptional programmes associated with 3D and 2D isolation. (A) Pooled 3D-versus-2D edgeR volcano plot. Positive log2FC denotes higher expression in pooled 3D MSCs. The displayed threshold is FDR < 0.001 and |log2FC| ≥ 1 (4,285 genes), whereas the broader DEG analysis uses FDR < 0.01 (4,840 genes). Numbered labels identify ENSG00000289901 (1), TNFSF10 (2), PRKG1 (3), MT-CYB (4), ENSG00000289474 (5), AKAP12 (6), PPP1R14B (7), CCND1 (8), KCNMA1 (9) and FABP5 (10). P/FDR values derive from non-independent random cell splits of one donor and do not provide population-level inference. (B) Pooled condition-level dotplots ranked by effect magnitude (upper) or FDR (lower). The displayed plots are labelled with an expression-prevalence filter of at least 20%. Colour indicates gene-centred pooled mean expression and dot size indicates the expressing-cell fraction. (C) Feature UMAPs for HES1, VCAM1, DDIT4, TNFSF10, CXCL12, PDGFRA, KCNMA1 and MKI67. (D) Violin plots across 3D C1/C2 and 2D C1–C4. (E) Eight selected functional groups from the pooled GSEA, containing all 129 significant sets assigned to these groups. Each point denotes a gene set at its original NES, and black diamonds denote the unweighted arithmetic mean of signed NES values within each group. Red indicates 3D association and blue indicates 2D association. The complete 20-group landscape includes all 381 significant sets (Supplementary Figure S7). Group means are descriptive, and overlapping sets are not independent confirmations. (F) CellPhoneDB reported-pair matrix for 3D day 21. Pair counts are hypothesis-generating expression-compatible candidates rather than measurements of communication strength. Cell counts describe representation in one donor and do not quantify absolute isolation yield.

Complementary pooled dotplots presented effect-size- and FDR-prioritized views of 3D-versus-2D expression (Figure 4B). Both plots display 20 genes per direction and are labelled with a minimum expression prevalence of 20%. Prominent displayed 3D effect-size candidates included WDR86, MT1F, CCL5, TREM1, COL5A3, IFITM1, EREG, AREG, HLA-F and CA9, spanning low-oxygen response, matrix, immune and growth-response biology. Displayed 2D candidates included GPNMB, KCNMA1, MFAP5, ANKRD1 and AKAP12. The FDR-prioritized plot included TNFSF10, GPC6, VCAM1, DCN, IFITM1 and C1R among 3D-associated genes, and KCNMA1, AKAP12, CCND1, ANKRD1, FABP5, CAV1 and CFL1 among 2D-associated genes. Neither ranking alone defines biological importance. Effect size, prevalence, expression level and cluster localization were considered together. Extended top-50 rankings and cluster-versus-rest profiles are provided in Supplementary Figure S5 and Supplementary Tables S7–S8.

Feature UMAPs and violin plots localized these pooled differences to individual MSC clusters (Figure 4C–D). HES1, VCAM1, DDIT4 and TNFSF10 were preferentially associated with 3D C1, with CXCL12 and PDGFRA providing complementary niche/progenitor context. CXCL12 was assessed descriptively because it was not retained in the pooled edgeR output. Each feature-UMAP scale was shared across all six clusters for the corresponding gene, and violin plots distinguished broad expression from small highly expressing fractions. HES1 showed pooled log2FC +3.72 (FDR 7.24 × 10⁻¹³) and a stronger 3D-C1-versus-rest signal (log2FC +4.75, FDR 3.01 × 10⁻³⁷). Its role in Notch-linked maintenance of bone-marrow mesenchymal progenitors [17] supported interpretation alongside GAS1, GREM1, KLF4, ID2, PDGFRA and PDGFRB, and the matrix-, adhesion- and cytokine-associated genes GPC6, VCAM1, DCN, TNFSF10, IFITM1 and HLA-B. Together, these features characterized a progenitor- and niche-response-associated 3D C1 cluster without establishing functional self-renewal or multipotency.

A targeted literature-guided comparison of the DEG results identified additional stromal-associated candidates (Supplementary Table S13). SERPING1 and EBF3 met the pooled DEG criteria in the 3D direction and were also higher in the 3D-C1-versus-rest analysis. EGR1 and FOSB were higher in the pooled 3D comparison. These observations did not form a uniformly native-associated signature. PTGDS was higher in 2D, while LEPR, KITLG, OAS2, PPARG, PDGFRA and PDGFRB did not meet the combined pooled FDR and effect-size criteria. TNFAIP6 was higher in the C1-versus-rest comparison but not in the pooled contrast. The table reports both concordant and non-concordant candidates without introducing a new DEG model or claiming donor-level inference.

The 2D compartment was transcriptionally heterogeneous despite its broad MSC annotation. KCNMA1 was expressed across several 2D clusters, while AKAP12, ANKRD1, CAV1/CAV2, PFN1 and CFL1 delineated membrane, caveolar, cytoskeletal and contractile programmes compatible with attachment, spreading and force transmission on planar plastic. A distinct 2D C3 cluster expressed MKI67 and TOP2A, separating a proliferative population from the larger metabolically and structurally adapted 2D C1/C2 compartments. The smaller 2D C4 cluster had a different membrane/endosomal profile. In contrast, 3D C2 comprised only 60 cells and was dominated by ribosomal, translation and selected complex-I signals, making its profile and proximity to 2D C1 particularly sensitive to sampling and QC. Pooled comparisons therefore required cluster-resolved interpretation rather than treating either condition as a uniform MSC population.

Gene-set enrichment was assessed separately to determine whether pathway-level patterns converged with the DEG results. At q <0.05, 381 of 8,506 tested gene sets were enriched, with 117 in the 3D direction and 264 in the 2D direction. The 381 significant sets were assigned to 20 descriptive functional groups. Eight selected groups containing 129 sets are shown in Figure 4E, and the complete pooled landscape is provided in Supplementary Figure S7. Group means summarize signed NES values rather than constituting separately tested pathway effects. Representative pooled and cluster-resolved terms are provided in Supplementary Table S9. The 3D-associated modules included TNF signalling (NES +2.13), interferon-γ response (+2.13), cytokine–receptor interaction (+2.04), collagen/ECM-related terms (+1.96) and cell-adhesion molecules (+1.94). Strong 2D-associated modules included mitochondrial translation elongation (NES −3.02), oxidative phosphorylation (−2.96) and respiratory electron transport and ATP synthesis (Reactome R-HSA-163200, NES −2.78), consistent with the gene-level contrast between 3D ECM/immune responsiveness and 2D bioenergetic/structural adaptation. Pooled GSEA nevertheless placed a G2/M module in the 3D direction, whereas the clearest cell-cycle-defined cluster was 2D C3. Pooled rank enrichment, cluster abundance and cluster-versus-rest expression measure different aspects of heterogeneity. This discrepancy requires cluster-resolved and donor-replicated follow-up.

Hallmark Hypoxia was enriched in the 3D direction (NES +1.44, q =0.048), alongside a coordinated CA9, EGLN3, NDRG1, DDIT4, PFKFB3, HK2, SLC2A3, EPAS1 and PDK1 profile concentrated in 3D C1. A large HIF1A mRNA change was not required, given the substantial post-translational regulation of HIF-1α activity. Both conditions were cultured at nominal external 5% O₂, excluding different incubator settings as the explanation. Although the approximately 500-µm medium-surrounded beads were designed to limit transport distances, neither this geometry nor the transcriptional profile directly measured intrabead oxygen. The findings were compatible with a niche-responsive metabolic profile maintained, induced or selected in 3D.

The enrichment pattern did not indicate a uniform increase in stress-related programmes in 3D. Hallmark Apoptosis (NES +1.05, q =0.483), Reactive Oxygen Species Pathway (−1.21, q =0.358) and Unfolded Protein Response (−0.86, q =0.978) did not reach q <0.05. Hallmark p53 Pathway (−1.54, q =0.0346), Reactome Stabilization of p53 (−1.91, q =0.0117) and Regulation of Apoptosis (−1.91, q =0.0115) instead favoured 2D. These results identified condition-associated stress-response components but did not establish a general difference in cellular injury, oxygen concentrations, viability or apoptosis rates.

The presence of macrophages alongside the cytokine- and ECM-associated 3D C1 cluster prompted condition-specific CellPhoneDB analysis. At 3D day 21, 438 ligand–receptor pairs were reported for 3D C1→3D C1, 200 for 3D C1→macrophages and 151 for macrophages→3D C1. Corresponding 3D C2 counts were 6, 49 and 8 (Figure 4F). The broader C1 repertoire was concordant with its cytokine and ECM programme, but pair numbers represent expression-compatible candidates, not communication strength or demonstrated physical and functional interactions. Directed candidates could be tested by stromal–macrophage co-culture, protein localization and ligand/receptor blockade (Supplementary Figure S9, Supplementary Table S11).

Palantir analysis provided additional context for the myeloid-rich 3D environment. Among 21,901 haematopoietic cells, an HSC-proximal computational root preceded erythroblast, megakaryocyte-progenitor, mast-cell, pDC, T-cell and macrophage terminal states. Along the inferred monocyte/macrophage lineage, CD34 declined, S100A8/S100A9 increased in early-to-intermediate regions, and LST1, MAFB, CD68 and CXCL8 marked overlapping later populations (Supplementary Figure S8, Supplementary Table S12). Late pseudotime bins were enriched for 3D cells but predominantly reflected day 7, not a monotonic day-7 to day-21 progression. This transcriptional ordering was compatible with a myeloid maturation continuum. It was neither experimental lineage tracing nor a direct test of the MSC isolation comparison.

Overall, 2D culture was associated with greater MSC representation and purity, together with mitochondrial, oxidative-phosphorylation, cytoskeletal, mechanoadaptive and proliferative programmes. The rarer 3D C1 cluster combined stromal and progenitor-associated identity with ECM/adhesion, cytokine/interferon, hypoxia/glycolysis and predicted macrophage-interaction features. This coordinated profile was compatible with aspects of native bone-marrow niche biology and a profile less completely adapted to planar culture. However, heterogeneous canonical markers, sparse NGFR/CD271 expression and potential effects of GelMA digestion limit this interpretation. Direct mapping to freshly isolated adult human bone-marrow stromal references remains a proposed follow-up. The present design cannot distinguish preservation, induction, selective survival or preferential recovery of 3D C1, and does not establish greater in-vivo similarity.

## Discussion

Direct isolation in GelMA microbeads revealed MSC-associated clusters within a heterogeneous bone-marrow culture that differed from those obtained by conventional two-dimensional isolation and culture. Whereas the final 2D single-cell dataset contained a strongly represented, exclusively MSC-annotated population, 3D C1 combined stromal identity with progenitor-associated, matrix-related, immune-responsive and metabolically adaptive programmes. The biological value of an isolation method therefore depends not only on the number of MSCs obtained, but also on the cell populations recovered and their cellular context. Our findings suggest a trade-off between efficient MSC selection and the recovery of niche-associated transcriptional profiles, supporting further development of direct 3D isolation as a distinct approach alongside established 2D expansion.

### Bulk GelMA as a starting platform for bone marrow culture

The bulk experiments first established that GelMA supported cellular activity over the investigated culture period. CAM-positive adMSCs up to day 21 and early EdU incorporation indicated viability-associated activity and DNA synthesis, respectively, without quantifying survival or net expansion. BM-MNC cultures contained rounded and elongated cells, but no clear increase in spindle-shaped cells. These observations were consistent with the low MSC-associated frequencies and the near-absence of the combined CD73/CD90/CD105 readout. None of the temporally compared individual MSC-marker readouts changed detectably after Holm correction (Figure 1 and Supplementary Table S4). Bulk GelMA therefore supported heterogeneous culture but did not generate an MSC-enriched population under the conditions examined.

The prominent CD235a⁺/CD45⁻ fraction and stronger PI signals in deeper or later images also identified constraints relevant to further development. Retaining multiple cell fractions may enable cellular interactions but creates additional requirements for transport and recovery. Spatial oxygen differences have been demonstrated experimentally in GelMA cultures, with material properties and cell behaviour influencing local oxygen availability [12]. Transport limitations are therefore plausible but were not directly measured in our bulk samples. Functionalization and rheological properties remain independently measured material parameters and do not permit a simple causal attribution of the cellular findings (Supplementary Figure S1 and Supplementary Tables S1–S2).

The higher MSC-associated frequencies in 2D are consistent with selection through plastic adherence and subsequent culture. Early culture stages are already biologically relevant. In a comparison of primary human bone-marrow stromal cells and their cultured progeny, Ghazanfari et al. identified substantial transcriptional changes at passage 0 [2]. The additional passage of Donor 3 may have contributed to variation among our 2D samples without reversing the shared direction of the contrast with bulk 3D. Subsequent evaluation therefore required marker enrichment and biological state to be considered separately.

### Microbeads as an advanced format for isolation and culture

The transition to medium-surrounded microbeads addressed culture handling and spatial organization. adMSC validation, the emergence of elongated and stellate cells, and confocal detection of cells within beads demonstrated that this format supported three-dimensional cellular organization (Figure 2). Related millifluidic GelMA systems support MSC networks and can be scaled for dynamic culture [13]. Our adaptation for direct bone-marrow isolation extends this technical basis to a different biological objective. Stromal cells are studied within a three-dimensional, multicellular environment from the isolation stage rather than only after prior 2D expansion.

In the GMB flow-cytometry pilot experiment, individual MSC marker frequencies were 27.16–33.41% and the triple-positive readout accounted for 25.49% of MSC-panel singlets. These frequencies were markedly higher than the corresponding bulk readouts (Figure 2F and Supplementary Tables S4–S5). With SEF comprising 48.77% of singlets, the triple-positive readout defined within this gate corresponded to approximately 52.3% of SEF. Thus, a substantial proportion of the operationally enriched fraction expressed the investigated MSC-marker combination. This is a promising descriptive observation, although the single-donor pilot and differences in sample preparation do not constitute a controlled comparison of effectiveness against bulk culture. SEF remains an exclusion gate rather than a pure or functionally validated MSC population.

The low CD235a⁺/CD45⁻ frequency of 1.15% may be compatible with reduced retention of non-adherent cells in the microbead system in addition to the preceding erythrocyte-lysis step. The smaller geometry and exposure to surrounding medium could facilitate cellular escape and subsequent removal during medium changes. This mechanism was not quantified and remains a hypothesis. Overall, the method is still under development, particularly with respect to culture conditions and cell recovery for reproducible isolation of MSC populations suitable for downstream applications.

### Cellular heterogeneity and temporal changes in bone marrow populations

The single-cell data allow a distinction between MSC enrichment and the presence of a multicellular bone-marrow context. The final dataset contained 12,870 MSC-annotated cells from 2D and 305 from 3D, including 245 in C1 and 60 in C2 (Figure 3). Cell loading was 15,970 for 2D day 21 and 20,000 for 3D day 21. However, final counts describe representation after capture and quality filtering rather than absolute MSC isolation yield per amount of marrow (Supplementary Table S6). The MSC-annotated 2D population was itself transcriptionally heterogeneous, encompassing four clusters. By contrast, 3D provided a context of stromal, myeloid and other haematopoietic populations in which additional interactions were possible. Single-cell and tissue analyses of human marrow support the biological importance of such spatially and functionally organized environments [8, 18].

The relative decline of early haematopoietic populations is particularly informative in this context. HSC- and GMP-annotated fractions accounted for 9.30% and 50.18% of D0 cells, respectively. Within 3D cultures, these fractions decreased from 1.25% to 0% and from 8.17% to 4.62% between days 7 and 21 (Figure 3C). D0 preselection limits the baseline comparison but cannot by itself explain the additional change between the two 3D time points. Meanwhile, the macrophage fraction increased from 4.38% to 9.24%, whereas monocytes remained abundant at 46.66% and 43.21%. These relative changes are compatible with a shift towards more differentiated myeloid populations, without establishing a specific differentiation route.

The remaining GMPs also demonstrate that the late 3D culture was not composed exclusively of terminally differentiated cells. GMPs are lineage-restricted progenitors rather than a functional substitute for long-term self-renewing HSCs. Moreover, the absence of an HSC annotation at day 21 does not establish the disappearance of every such cell from the entire culture. The appropriate interpretation is therefore a heterogeneous culture containing residual progenitor-associated populations but a reduced representation of early haematopoietic populations. Complete preservation of the original composition or a stable equilibrium cannot be inferred.

Trajectory analysis adds transcriptional context to the compositional changes. Along the inferred monocyte/macrophage lineage, CD34 decreased, S100A8/S100A9 increased in early-to-intermediate regions, and LST1, MAFB, CD68 and CXCL8 marked overlapping later populations (Supplementary Figure S8 and Supplementary Table S12). This pattern is compatible with myeloid maturation. However, Palantir orders transcriptional states rather than experimentally tracing cellular ancestry [19]. Because late pseudotime regions predominantly contained day-7 cells, they do not simply represent the later culture time point. Composition and trajectories therefore provide complementary evidence concerning culture development.

These findings suggest that GMBs support some stromal–haematopoietic relationships and progenitor populations but do not yet reproduce all conditions required to maintain the most primitive populations. In mouse models, HSC maintenance depends in part on locally supplied stem cell factor from endothelial and perivascular stromal cells [20]. Human spatial atlases likewise identify distinct neighbourhoods of early haematopoietic and myeloid states [18]. Incompletely organized vascular contacts or niche signals could therefore contribute to the observed shift. This identifies a possible biological limitation of the current model without asserting a measured deficiency of any specific factor.

### Niche responsive MSC programmes and prospects for defined reconstruction

The strongest evidence for a distinct 3D MSC cluster lies in the coordinated C1 profile. HES1, GAS1 and GREM1 occurred alongside stromal identity markers, VCAM1 and matrix-associated genes such as DCN. GSEA complemented this pattern with ECM, adhesion and cytokine programmes (Figures 3–4 and Supplementary Figures S5–S7). This profile is compatible with enhanced responsiveness to the cellular and extracellular environment. Its relevance derives from the combination of programmes and literature-based comparison with defined stromal reference populations, rather than from any single presumed marker of a native state.

Primary human data provide specific points of comparison. Li et al. identified a stromal population expressing CXCL12, LEPR and DCN, together with predicted VCAM1-mediated contacts with haematopoietic cells [8]. Fiévet et al. described an MSC programme in uncultured adult marrow that included CXCL12, PDGFRA and LEPR and conserved adhesion and matrix genes such as VCAM1 and DCN [21]. Higher pooled VCAM1 and DCN expression and complementary CXCL12 and PDGFRA signals in our 3D clusters are therefore biologically consistent with these observations. PDGFRB supports stromal or perivascular identity but is not MSC-specific and also identifies mural cells in human references [21].

An additional comparison of DEG results strengthens this connection for selected genes (Supplementary Table S13). SERPING1 was higher both in the pooled 3D comparison and in C1 relative to the other MSC clusters. This agrees with Ghazanfari et al., who assigned SERPING1 to a human programme with higher expression in primary and sphere-cultured cells than in adherent cultures [2]. EBF3 was also higher in both comparisons. Genetic studies of murine CAR/LepR-positive populations support its relevance to niche-forming stromal cells, but do not directly establish a corresponding function in our human cells [22]. EGR1 and FOSB provide further overlap with primary human data [2, 21]. As immediate-response genes, however, they may also reflect preparation or activation and cannot independently establish greater similarity to in-vivo cells [23].

The correspondence remains partial. CXCL12 was examined descriptively and was not retained in the pooled DEG table. LEPR was detected in approximately one quarter of C1 cells but, like KITLG, did not meet the pooled DEG criteria. PDGFRA and PDGFRB did not reach the combined effect-size threshold. Conversely, PTGDS, which is reported in native human stromal populations, was higher in 2D, whereas OAS2 and PPARG did not meet the combined DEG threshold (Supplementary Table S13). NGFR/CD271 transcripts were also very rare. These counterexamples favour comparison of related programmes and appropriate stromal subtypes rather than equating 3D C1 with freshly isolated MSCs [2, 8, 21]. A future comparison with published data from freshly isolated human bone-marrow stromal cells would therefore be particularly informative.

HES1 adds a possible regulatory link to progenitor state. Its pronounced C1 signal is consistent with mouse studies in which Notch signalling preserved mesenchymal progenitors by restraining osteoblast differentiation [17]. In murine lineage-tracing experiments, Grem1 identified osteochondroreticular stem cells with bone, cartilage and stromal potential [24]. Gas1, by contrast, was associated with transiently amplifying descendants of an endosteal stem-cell population, representing a downstream progenitor stage rather than exclusively the most primitive stem cells [25]. Together, these genes provide plausible links to distinct progenitor programmes. They do not yet establish greater long-term self-renewal or functional multipotency of the human 3D C1 cells.

Comparison with cultured cells is equally important. Ghazanfari et al. described both shared culture-induced changes and programmes that primary cells shared more strongly with mesenspheres than with adherent cultures [2]. In human and murine data, Stalmann et al. observed a shift of cultured stromal cells towards fibroblast-like profiles despite retained haematopoietic support [3]. Our 2D-associated mitochondrial, oxidative and cytoskeletal programmes can therefore be interpreted as culture adaptation. AKAP12, ANKRD1, CAV1/CAV2, PFN1 and CFL1 are consistent with altered adhesion and force transmission, while MKI67/TOP2A-positive 2D C3 shows additional proliferative specialization. Such adaptation may support expansion and does not necessarily imply functional loss. The broad cell-cycle/DNA-repair group is predominantly 2D-associated but includes 3D-associated subprogrammes such as G2/M. Neither the group summary nor an individual cluster quantifies the proliferation rate of the entire culture.

The metabolic contrast also has a plausible niche-related interpretation. CA9, EGLN3, NDRG1, DDIT4 and glycolysis-associated genes, together with hypoxia enrichment, indicate coordinated adaptation in 3D C1 (Figure 4 and Supplementary Table S9). In human MSCs, HIF-associated glycolysis has been linked to an undifferentiated state, whereas osteogenic differentiation was accompanied by an oxidative shift [26]. Direct measurements in living mouse marrow demonstrate low and spatially heterogeneous oxygen tensions [27]. These studies provide biological context for the observed metabolic configuration. However, both cultures were maintained at nominal 5% O₂. The signature may reflect differences in local supply or cellular state and does not substitute for oxygen measurements. Non-significant enrichment of apoptosis, ROS or UPR programmes likewise does not establish an absence of cellular injury.

Cytokine and interferon programmes and predicted contacts between 3D C1 and macrophages connect these clusters to their culture environment (Figure 4F, Supplementary Figure S9 and Supplementary Table S11). Chow et al. demonstrated in mice that CD169-positive macrophages contribute to haematopoietic stem- and progenitor-cell retention through stromal niche cells [28]. Our macrophages were not functionally identified as this CD169-positive subtype. Nevertheless, the study provides a rationale for considering accompanying myeloid cells as more than contaminants. CellPhoneDB supplies testable interaction candidates rather than measurements of communication strength or demonstrated signalling [29]. Immune-responsive MSC programmes likewise do not by themselves establish immunosuppressive function.

An alternative to direct 3D isolation would be to isolate and expand MSCs in 2D before transferring them into GelMA with defined macrophages. The remaining plasticity of cultured MSCs supports this possibility. Bartosh et al. enhanced anti-inflammatory properties of human MSCs through spheroid formation [30], while Rao et al. partially restored selected surface markers and secretory functions after transfer to soft PEG hydrogels [31]. Direct co-culture of primary human macrophages and MSCs in methacrylated gelatin has also been established experimentally [32]. These findings support the feasibility of a more defined and potentially more scalable culture, but do not demonstrate reproduction of our C1 profile. Our TNFAIP6 result is also cluster-dependent, with higher expression in the C1-versus-rest comparison but no pooled 3D-versus-2D difference (Supplementary Table S13).

Cellular history argues against assuming complete equivalence. Subpopulations lost during adherence selection cannot be recovered simply by changing the matrix. Yang et al. also showed that the duration of prior culture on stiff substrates influenced subsequent YAP/TAZ and RUNX2 activity in human MSCs on softer hydrogels [33]. Studies in bovine MSCs provide a possible chromatin-related basis for mechanical memory, although under dynamic tensile loading rather than our culture conditions [34]. These findings support possible persistent priming, not a universally irreversible effect of 2D culture. Partial restoration on hydrogels and the responsiveness of cultured states argue against such a generalization [3, 31].

The macrophage component also requires functional consideration. In the gelatin co-culture studied by Romero-López et al., macrophages promoted osteogenic differentiation, particularly under pro-inflammatory polarization [32]. Guihard et al. identified oncostatin M from activated monocytes or macrophages as one mechanism promoting osteogenesis [35]. Co-culture could therefore induce selected 3D-associated programmes while favouring differentiation rather than progenitor maintenance. Macrophage origin, activation state, cell ratio and contact duration are consequently relevant variables. Direct 3D isolation remains particularly interesting if the C1 profile emerges from an early complex cellular community that a subsequently assembled two-cell system only partially reproduces.

This interpretation is based on a single-cell dataset from one donor. Individual cells and random cell splits do not replace independent biological replicates. The contribution of cell recovery to the observed clusters also remains to be clarified. These limitations constrain generalization without negating the cluster differences observed within the dataset.

For cell therapy and human laboratory models, the value of this approach lies in recovering and studying context-dependent stromal phenotypes. Therapeutic applications require reproducible identity, safety and indication-relevant function rather than an unspecified claim of greater stemness. In drug-testing models, joint assessment of stromal and haematopoietic responses could provide additional information. Vascularized human bone-marrow models with measurable drug toxicities and recovery responses demonstrate the practical potential of such interactions [36]. Our GMB culture has not reached that level of validation but provides a justified starting point. Further development should improve recovery and scalability while preserving the niche-responsive programmes that distinguish direct 3D isolation from conventional expansion.

## Conclusion

Direct GelMA microbead isolation recovered a rare MSC population with niche-associated transcriptional profiles within a heterogeneous bone-marrow culture, whereas conventional 2D isolation produced a more strongly represented and compositionally purer MSC compartment. The coordinated progenitor-associated, matrix, adhesion, cytokine and metabolic programmes of 3D C1 overlapped with selected features reported for native bone-marrow stroma and differed from the culture-adapted 2D clusters. These findings support further development of direct 3D isolation for studying context-dependent stromal biology. Direct reference comparisons, independent donors and functional validation are needed to determine whether the observed clusters offer advantages for cell therapy or human experimental models.

## Declarations

### Ethics approval and consent to participate

The study was conducted in accordance with the Declaration of Helsinki. The use of human adipose tissue was approved by the Ethics Committee of the University of Lübeck (20-233 and 2024-455_1). The use of femoral-head bone marrow and iliac-crest bone marrow was approved by the Ethics Committee of Hannover Medical School (565-2009 and 565-2016 for femoral-head material, and 2858-2015 for iliac-crest material). Written informed consent was obtained from all donors.

### Author contributions

NHussein: Conceptualization, Methodology, Investigation, Formal analysis, Data curation, Validation, Visualization, Writing – original draft. CM: Software (bioinformatics), Formal analysis, Methodology, Visualization, Data curation, Writing – original draft, Writing – review & editing. YR: Methodology, Resources. LV: Methodology, Resources. MS: Methodology, Resources. MH: Methodology, Resources. AL: Methodology, Resources, Writing – review & editing. NHaase: Writing – Review & Editing, Supervision (bioinformatics). SR: Writing – Review & Editing, Supervision (bioinformatics), Methodology, Resources. AH: Methodology, Resources, Project administration, Funding acquisition, Writing – review & editing. DE: Conceptualization, Resources, Supervision, Project administration, Funding acquisition, Writing – review & editing.

## Acknowledgements

The authors gratefully acknowledge the expert technical assistance by Anika Hamm and Kirsten Elger as well as the technical support by the Research Core Units Genomics and Cell Sorting of MHH. In addition, all authors are deeply indebted to the patients and voluntary donors for their bone marrow donations and thank Maike Keck for support in acquisition of adipose tissue donations.

## Generative AI statement

The authors declare that generative artificial intelligence (ChatGPT 5.5/6, OpenAI) was used during manuscript preparation for translation, spelling and grammar correction, and revision of selected passages to improve linguistic clarity and conciseness. Figure 2A was designed by Nali Hussein and illustrated with assistance from ChatGPT (OpenAI). All scientific content, data interpretation and phrasing were reviewed and approved by the authors.

## Data availability

The data supporting the findings of this study are available from the corresponding author upon reasonable request. Bioinformatic analyses pipeline: All necessary details are provided in the Methods and Supplementary Information.

