## Supplementary Information for "Direct 3D isolation in gelatin methacryloyl microbeads versus conventional 2D isolation and culture: a trade-off between mesenchymal stromal cell purity and niche-associated transcriptional profiles"

### Supplementary Methods

#### SM1 GelMA synthesis

Porcine-skin gelatin, type A, 140 Bloom (Carl Roth, 20 g), was dissolved in 200 ml of 0.25 M carbonate buffer for 30 min at 45 °C and adjusted to pH 9 with NaOH [11, 15]. MAA (Sigma-Aldrich) was added dropwise: 1 ml for low DoF and 2 ml for high DoF. After 3 h, the pH was adjusted to 7.4 with HCl. Products were dialysed at approximately 45 °C using a 14-kDa-MWCO membrane against 4 l double-distilled water, changed daily over four days, then frozen at −20 °C and lyophilized. Material was UV-treated for 30 min, turned over and irradiated for a further 30 min. It was stored in 50-ml tubes in the dark at room temperature. Medium DoF was prepared by mixing equal volumes of the two 5% BATCH1 solutions. BATCH2 high DoF was used separately for GMBs.

#### SM2 $^1$H NMR acquisition and independent raw-FID processing

For NMR, 20 mg material was dissolved in 700 µl $D_2O$ (99.9%, Deutero). Spectra used for DoF estimation were acquired at 313 K on an Avance Neo (Bruker) at a $^1$H frequency of 600.33 MHz. Raw metadata document the zg pulse programme, eight scans, eight dummy scans, a 2-s relaxation delay, a spectral width of 10,869.57 Hz and 131,072 time-domain points (corresponding to a 6.03-s acquisition, calculated as TD/[2 × spectral width]). Unmodified gelatin from the same starting lot served as reference. Measurements at 298 K provided a supplementary temperature comparison. Raw FIDs underwent Bruker digital-filter removal, 0.3-Hz exponential line broadening, zero filling to 131,072 points, Fourier transformation and automatic phase correction. Phase correction used ACME/peak-minima candidates followed by visual quality control. Spectra were aligned by cross-correlation in the phenylalanine region and normalized to the baseline-corrected phenylalanine integral at 7.00–7.50 ppm.

The reference lysine ε-$CH_2$ signal shape was fitted over 2.95–3.15 ppm together with a cubic background, using 0.04-ppm flanking anchors and a non-negative lysine amplitude. DoF was calculated as $100 \times (1 - A_{GelMA}/A_{gelatin})$, where A denotes the normalized fitted lysine amplitude. Vinyl signals provided supporting evidence of methacryloyl groups rather than an alternative DoF definition. Processing sensitivity was assessed using 0- and 0.3-Hz variants, with 0.5 Hz evaluated separately as a technical stress test. Independent raw-FID processing used Python 3.12.13, NumPy 1.26.4, SciPy 1.16.1 and nmrglue 0.11.

#### SM3 Rheology

Time-dependent storage (G′) and loss (G″) moduli of cell-free 5% (w/v) BATCH1 medium- and high-DoF and BATCH2 high-DoF formulations were measured using an MCR 302 (Anton Paar) at 37 °C. A 20-mm parallel-plate geometry, a 0.5-mm gap and a frequency of 1 Hz were used. Measurement formulations contained 0.2% (w/v) LAP. After an 18-s baseline, samples were exposed to 365-nm LED light for 180 s, followed by approximately 40 s of post-irradiation monitoring. The reported G′ and G″ values were obtained at the end of this observation period. Each pair represents one observation from the reported measurement run for each formulation, rather than a mean across independent replicates (Supplementary Table S2). Rheological photocrosslinking at 365 nm differed from the 405-nm crosslinking used for cell-culture gels. GMB culture formulations contained 0.4% LAP, whereas the rheological test formulations contained 0.2%.

#### SM4 Single-cell analysis pipeline

##### Generation of the scRNA-seq count matrix

The dataset comprised four samples of human bone marrow mononuclear cells (MNCs) from a single donor: day 0, day 7 (3D), day 21 (2D), and day 21 (3D). Raw FASTQ files generated on the 10x Genomics platform were processed with Cell Ranger (v10.0.0) [16] using the multi pipeline, which performs demultiplexing, read alignment, and gene quantification (cellranger multi). Reads were aligned to the human reference genome GRCh38 (10x Genomics reference version 2024-A). Because the resulting count matrices contain ambient RNA and empty droplets, both were removed with CellBender (v0.3.2) [37] using the remove-background command with default parameters (cellbender remove-background).

##### Iterative preprocessing and quality control

**Sample preparation and QC metrics.** Each of the four samples was assigned a file name encoding its metadata (e.g., MNC_2D_D21.h5). Files were read with Scanpy (v1.11.5) [38] (sc.read_h5ad), and the metadata fields were parsed from the file names and stored as columns in .obs. Mitochondrial, ribosomal, and hemoglobin genes were identified by gene name. QC metrics were computed for each cell with scanpy.pp.calculate_qc_metrics, including total counts, number of detected genes, the percentage of counts attributable to each gene group, and the percentage of counts in the top 20 genes. In addition, doublets were detected separately in each sample using scDblFinder (v1.24.0) [39] with default parameters and a random seed of 123, which yielded a continuous doublet score and a singlet/doublet classification for each cell.

**Flagging of outliers.** The prepared samples were concatenated into a single AnnData object (anndata.concat), and low-quality cells were flagged using per-sample thresholds. Flagged cells were identified separately for each sample based on the unscaled median absolute deviation, $MAD = median(|x_i - median(x)|)$. A cell was flagged as an outlier if its log1p-transformed total counts, its log1p-transformed number of detected genes, or its percentage of counts in the top 20 genes deviated from the sample median by more than 5 MADs, or if its percentage of mitochondrial counts deviated by more than 3 MADs. Furthermore, cells with fewer than 200 detected genes, fewer than 200 total counts, or a mitochondrial percentage above 8% were flagged.

**Delayed filtering.** Because the dataset was highly heterogeneous across samples, applying these thresholds directly would have removed many biologically relevant cells. Flagged cells (outliers and doublets) were therefore retained until clustering was complete, so that each cluster could be evaluated and, if appropriate, excluded from filtering (whitelisted).

**Clustering for cluster evaluation.** To obtain these clusters, the count matrix was normalized to 10 000 counts per cell (scanpy.pp.normalize_total) and log1p-transformed (scanpy.pp.log1p) to attenuate the influence of extremely highly expressed genes. The 2 000 most highly variable genes were selected with scanpy.pp.highly_variable_genes. A copy of the normalized matrix restricted to these genes was scaled with a maximum value of 10 (scanpy.pp.scale), and principal component analysis (PCA) was performed with 30 components (scanpy.pp.pca). A neighborhood graph was constructed from the PCA embedding (k = 15 neighbors) (scanpy.pp.neighbors), followed by Uniform Manifold Approximation and Projection (UMAP) [40] (scanpy.tl.umap) and Leiden clustering [41] at a resolution of 3 (scanpy.tl.leiden). A random seed of 0 was used for PCA, neighborhood graph construction, UMAP, and Leiden clustering. The PCA and UMAP embeddings, the neighborhood graph (distances and connectivities), and the Leiden cluster labels were transferred to the original AnnData object, such that all cells share the same embedding and cluster assignment.

**Cluster whitelisting.** For each cluster, the top 10 marker genes were ranked using the Wilcoxon rank-sum test. Clusters with a large proportion of flagged cells were examined based on their QC metrics, doublet fraction, and marker genes, and were whitelisted if they showed no evidence of low quality or doublet contamination. To ensure reproducibility, a cluster was considered for whitelisting only if it met four criteria, applied in the following order:

1. **Flagged cells:** At least 50% of the cells were flagged as outliers or doublets by the per-sample filter.
2. **Per-cluster MAD filtering:** Outlier thresholds were recalculated within the cluster. If fewer than 20% of the cells were still flagged, the cluster was automatically filtered using these cluster-specific thresholds and was not considered further for whitelisting.
3. **Mitochondrial content:** The 90th percentile of the mitochondrial count percentage was below the hard threshold (8%); clusters exceeding this threshold were excluded as likely dying cells.
4. **Doublet fraction:** Fewer than 15% of the cells were classified as doublets; clusters exceeding this threshold were excluded as predominantly doublets.

Clusters meeting these criteria were inspected manually based on their marker genes and QC metrics and were whitelisted where appropriate. The criteria informed the manual curation rather than replacing it, allowing clusters to be assessed in their biological context. In total, nine clusters were whitelisted (Supplementary Table S6E).

**Final filtering.** The filtering of each cluster depended on its classification during whitelisting. Clusters not meeting the first criterion were filtered using the per-sample thresholds, and clusters filtered automatically in the second criterion were filtered using per-cluster MAD thresholds. Clusters excluded by the third or fourth criterion were removed entirely. Of the nine whitelisted clusters, the two containing fewer than 150 cells (clusters 57 and 58; n = 180 cells in total) were retained in full, because the per-cluster MAD is unreliable for small, homogeneous clusters with little internal variance, whereas outliers and doublets in the remaining seven were removed using per-cluster MAD thresholds. To preserve the raw count matrix, the unprocessed files were reloaded and a copy was generated from which the filtered cells were removed. This filtered dataset served as the basis for all subsequent analyses.

##### Reference-based cell type annotation

To obtain an initial annotation, the filtered cells were classified with CellTypist (v1.7.1) [42] using three reference models. The first was the published Immune_All_Low model, which was used without modification. The other two models were trained on the annotated bone marrow reference atlases of Bandyopadhyay et al. [18] (GEO accession GSE253355) and Zeng et al. [43] (CELLxGENE collection ID f6c50495-3361-40ed-a819-fb9644396ed9). For training, raw counts were normalized to 10 000 counts per cell and log-transformed, and the 3 000 most highly variable genes were selected. Each model was trained independently on its respective reference using a logistic regression classifier optimized by mini-batch stochastic gradient descent (batch size of 2 048, 100 batches per epoch, 10 epochs) (celltypist.train). The random seed was fixed to 42.

For cell type prediction, each of the four samples was processed separately. Genes detected in fewer than 10 cells were removed, and raw counts were stored in a layer named counts for downstream analyses. Counts were then normalized and log-transformed as described above. All three models were applied independently to every cell without majority voting, yielding one predicted label and one confidence score per model for each cell (celltypist.annotate).

Because the accuracy of reference-based predictions depends on how well the reference matches the data, these predictions were complemented by another metric, the marker gene score. This score was based on the lineage-associated gene groups defined in Supplementary Table S6F. A separate score was calculated for each gene group and cell using the Scanpy function scanpy.tl.score_genes, and the median score of each lineage was then determined for each cluster.

##### Manual cell type annotation

Final cell type labels were assigned manually based on the CellTypist predictions, the lineage gene group scores described above, and canonical markers from single-cell expression databases such as PanglaoDB [44]. Each sample was annotated separately. To enhance annotation resolution, the samples were deliberately clustered at a fine resolution, and clusters assigned the same cell type label were subsequently merged. The resulting annotation defined the MSC populations used in the following analysis.

##### Differential expression analysis of 2D and 3D mesenchymal stromal cells

Differential expression between the MSC clusters of the 2D and 3D cultures was assessed using edgeR (v4.8.2) [45]. The raw count layer of the combined 2D day 21 and 3D day 21 samples served as input. Because independent biological replicates were not available, the cells of each condition were randomly assigned to three pseudoreplicates (random seed 42). Pseudobulk profiles were generated by summing the counts of each gene across the cells of each pseudoreplicate. The pseudobulk counts were imported into edgeR to create a DGEList object. Lowly expressed genes were removed with filterByExpr, and library sizes were normalized using the trimmed mean of M-values (TMM) method [46]. A linear model with condition (2D vs. 3D) as the sole covariate was specified, dispersions were estimated with estimateDisp, and a generalized linear model (GLM) was fitted to the data with glmQLFit. Differential expression was tested using the quasi-likelihood F-test (glmQLFTest), and p values were adjusted for multiple testing by the Benjamini–Hochberg procedure [47]. The pooled day-21 comparison contrasted 3D C1/C2 with 2D C1–C4, with positive log2 fold changes indicating higher expression in 3D. DEG summaries used FDR $< 0.01$ and $|\log2FC| \geq 1$. The volcano plot used an FDR threshold of 0.001. Because the pseudoreplicates originate from a single sample per condition, they do not capture biological variability between donors, and the results should therefore be considered exploratory.

##### Gene set enrichment analysis

To identify biological processes that differ between the two culture conditions, gene set enrichment analysis (GSEA) [48] was performed on the differential expression results using GSEApy (v1.3.1) [49]. All genes were ranked by $\log_2$ fold change without prefiltering for significance (gseapy.prerank). Gene set libraries were retrieved via Enrichr [50]: GO Biological Process, Molecular Function, and Cellular Component (2023) [51, 52]; KEGG Human (2021) [53]; MSigDB Hallmark (2020) [54]; Reactome (2022) [55]; and WikiPathways Human (2023) [56]. The analysis used 1 000 permutations (random seed 42) and gene set sizes of 5–500 genes after matching to the ranked list. Gene sets with a false discovery rate (FDR) $< 0.05$ and an absolute normalized enrichment score (NES) $> 1$ were considered significantly enriched, with positive NES values indicating enrichment in 3D and negative values indicating enrichment in 2D.

##### Cell–cell communication analysis

Ligand–receptor interactions between annotated cell types were inferred separately for each sample using CellPhoneDB (v5.0.0) [57]. Raw counts from the counts layer of each AnnData object were normalized to 10 000 counts per cell and log-transformed using Scanpy. Interactions were evaluated only if both the ligand and the receptor were expressed in more than 10% of the cells within the respective sender and receiver populations (cpdb_statistical_analysis_method.call). Statistical significance was assessed by a permutation test with 1 000 iterations, in which cell type labels were randomly shuffled (random seed 42), and interactions with $p < 0.05$ were considered significant. Significant interactions were then counted for each ordered cell-type pair, with the resulting counts interpreted as the number of predicted interactions rather than a measure of communication strength.

##### Trajectory analysis

To infer developmental relationships between cell lineages, a trajectory analysis was performed.

**Data integration.** Prior to trajectory inference, the day 0, day 7 (3D), and day 21 (3D) samples were integrated using scVI (v1.4.2) [58, 59] (scvi.model.SCVI.setup_anndata). The model was trained on raw counts (counts layer) restricted to the 2 000 most highly variable genes to reduce noise and improve training efficiency, with the sample specified as the batch covariate to mitigate technical effects introduced by cell culture (scvi.model.SCVI). Hyperparameters were optimized by grid search over 50 parameter combinations (Supplementary Table S6G) and 50 epochs, and the configuration yielding the highest evidence lower bound (ELBO) was selected. This configuration consisted of a latent dimensionality of 10 ($n_{latent} = 10$), two hidden layers, a zero-inflated negative binomial (ZINB) gene likelihood, a learning rate of $10^{-3}$, and a dropout rate of 0.1 (model.train).

**Pseudotime and fate probabilities.** A neighborhood graph (k = 30 neighbors) was constructed from the scVI latent representation, and a diffusion map with 10 components was computed from this graph using Palantir (v1.4.5) [19] (palantir.utils.run_diffusion_maps). Both steps used a random seed of 42. Pseudotime and fate probabilities were then estimated from a defined root cell and six terminal states: erythroblasts, megakaryocyte progenitors (MkPs), macrophages, mast cells, plasmacytoid dendritic cells (pDCs), and T cells (root and terminal cell barcodes are provided in Supplementary Table S6D) (palantir.core.run_palantir). For this step, the number of neighbors was set to k = 30, the number of waypoints to 1 200, and the random seed to 42. To visualize gene expression trends along pseudotime, expression values were imputed by MAGIC [60] as implemented in Palantir (palantir.utils.run_magic_imputation), and the resulting trends were smoothed with a smoothness factor of 4.0 and a length scale of 0.7 (palantir.presults.compute_gene_trends).

Lineage assignment. Cells were assigned to lineages using the Palantir function palantir.presults.select_branch_cells based on pseudotime and fate probabilities. Branch membership was defined by a fate-specific quantile threshold q on the fate probability, such that only cells with a strong commitment to the respective fate were retained. Because the distributions of fate probabilities differ between lineages, q was set individually for each fate: erythroblasts, 0.02; MkPs, 0.12; macrophages, 0.20; mast cells, 0.05; pDCs, 0.02; and T cells, 0.005. The parameter ε was set to 0, so that branch membership was determined exclusively by q. For visualization, trajectories were displayed as arrows on the embedding, with a smoothness of 0.25 for the erythroid, megakaryocyte, pDC, mast cell, and T cell lineages and 0.7 for the macrophage lineage (palantir.plot.plot_trajectories).

#### Supplementary Figures

#### Supplementary Figure S1

**A**

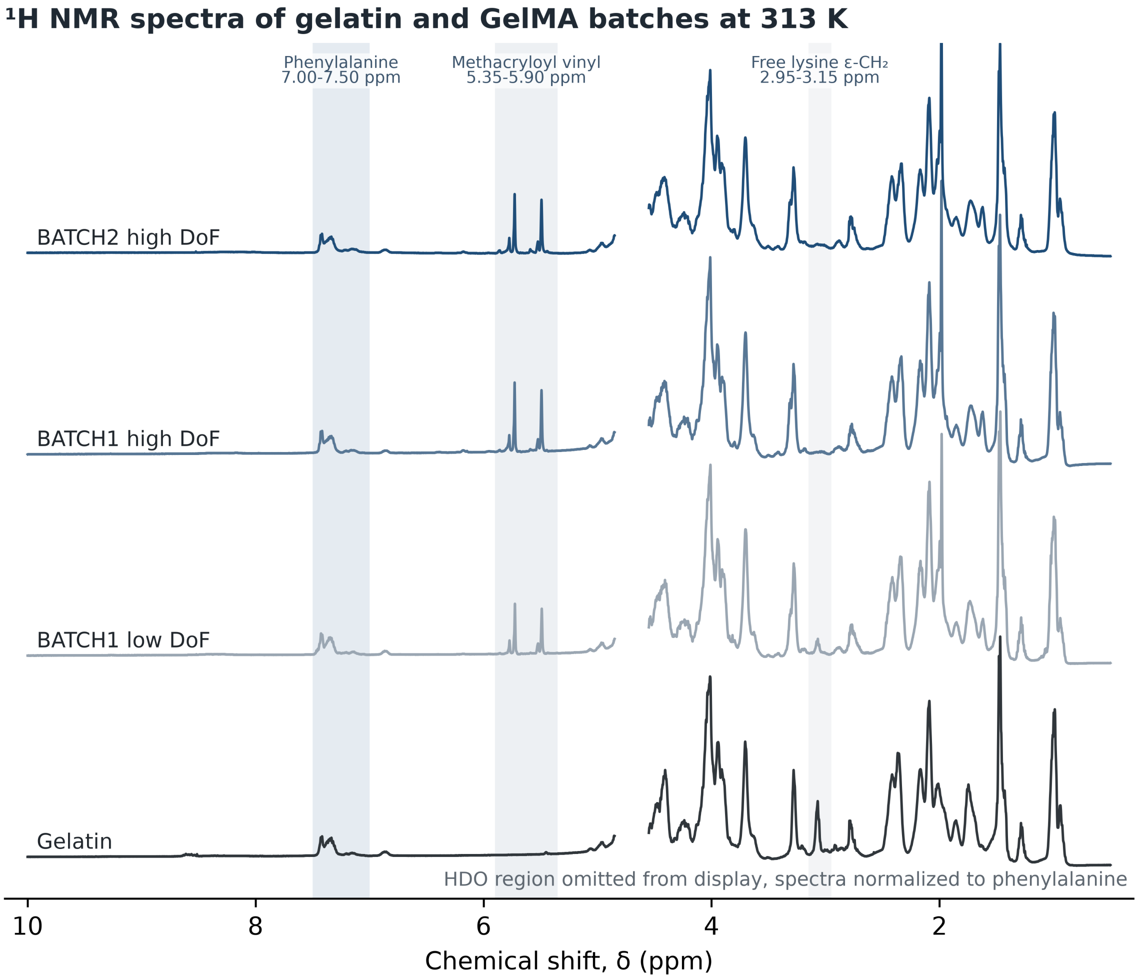

**B**

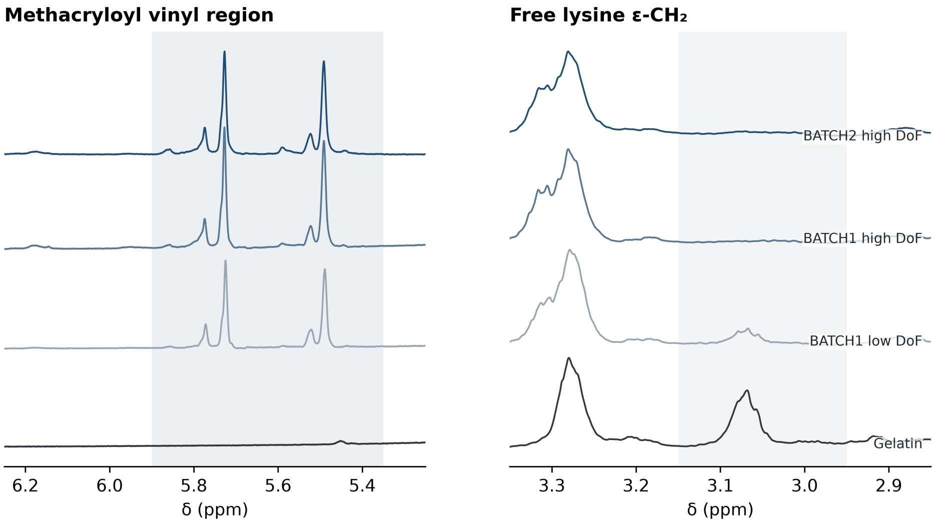


**C**

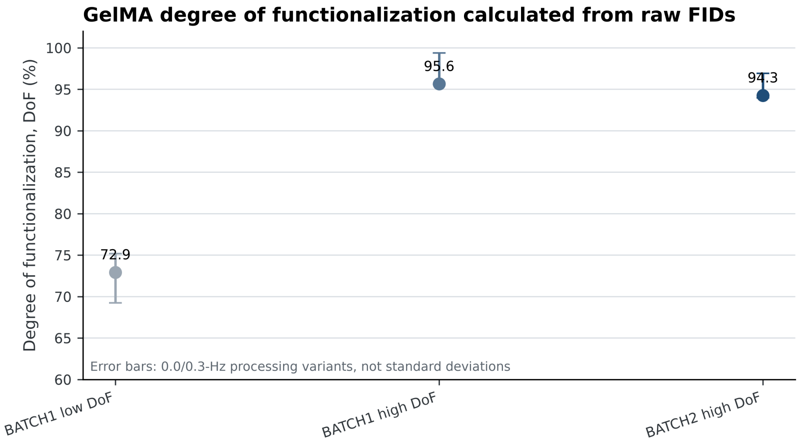


Supplementary Figure S1. Independent 313-K $^1$H-NMR characterization of the GelMA batches. (A) Full spectra of unmodified gelatin, BATCH1 low DoF, BATCH1 high DoF and BATCH2 high DoF. The HDO region is visually limited so that the polymer signals remain visible, but it was not deleted from the acquired data. (B) Expanded methacryloyl-vinyl and free lysine-ε-$CH_2$ regions used for qualitative confirmation of methacryloylation and primary DoF estimation. Normalization used the phenylalanine region in the full spectra shown in A. (C) Primary DoF estimates at 0.3-Hz line broadening with the standard 0/0.3-Hz processing range. BATCH1 low DoF was 72.9% (69.2–75.2%), BATCH1 high DoF 95.6% (95.6–99.4%) and BATCH2 high DoF 94.3% (94.0–96.9%). The range is a processing-sensitivity interval, not a biological SD or confidence interval.

#### Supplementary Figure S2

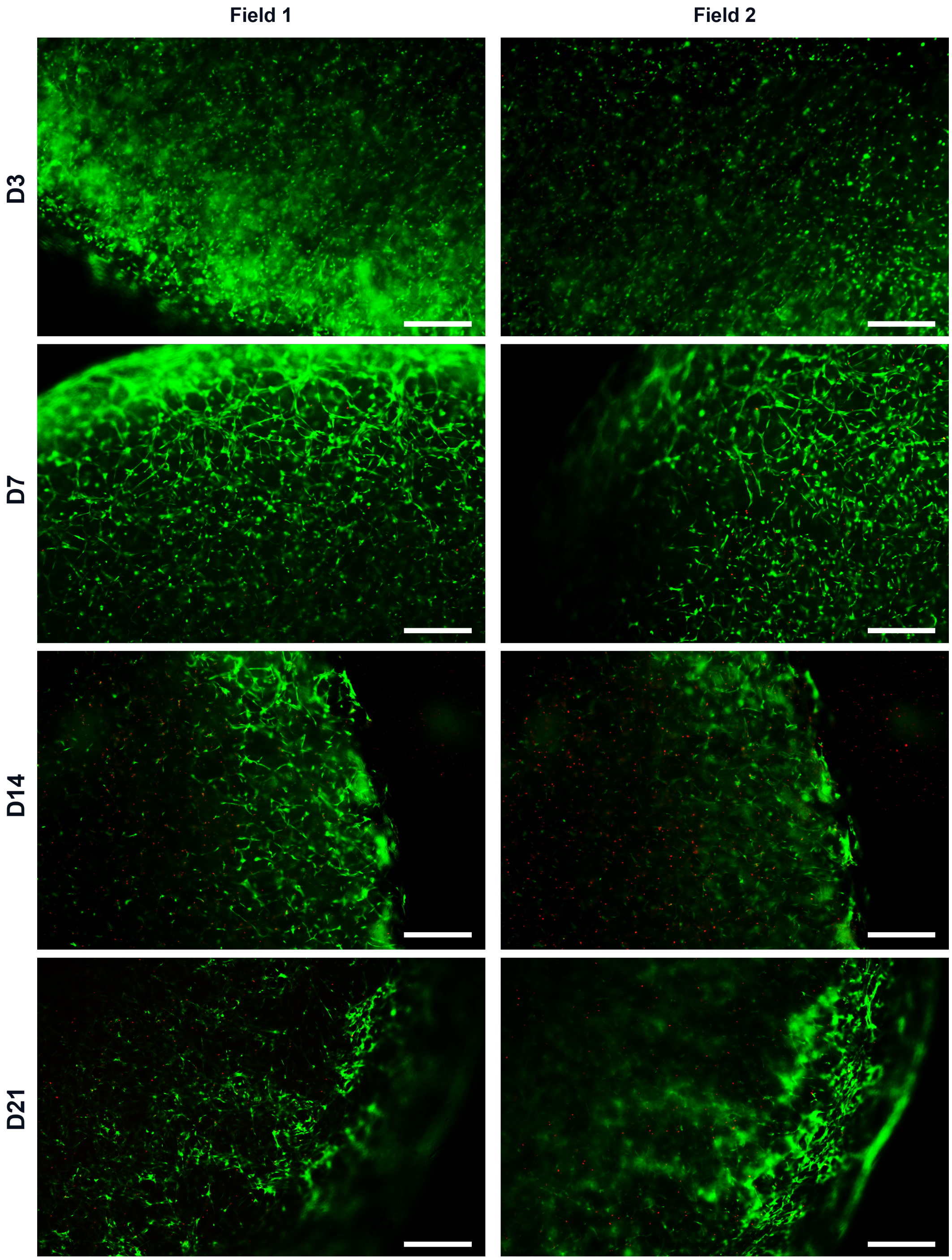


Supplementary Figure S2. CAM/PI microscopy of bulk-adMSC cultures at surface-proximal and deeper focal planes. Representative images are shown for days 3, 7, 14 and 21. Field 1 denotes a focal plane nearer the hydrogel surface, whereas Field 2 denotes a focal plane further into the hydrogel. Exact imaging depths were not recorded. Calcein AM (CAM) is shown in green and propidium iodide (PI) in red. Differences between fields may reflect both spatial heterogeneity and depth-dependent optical attenuation and should therefore not be interpreted as quantitative differences in viability. White scale bars represent 500 µm.

#### Supplementary Figure S3

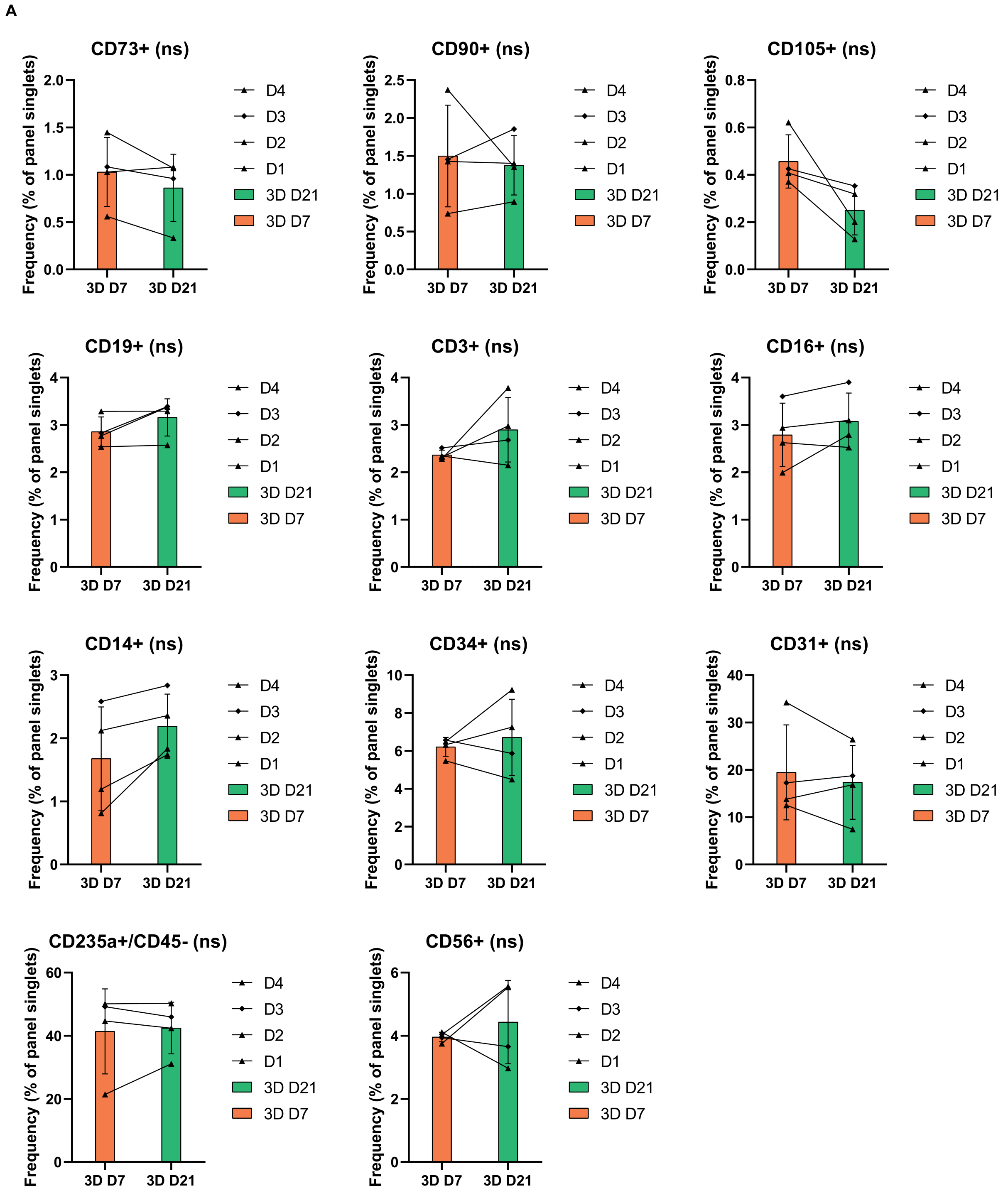

B

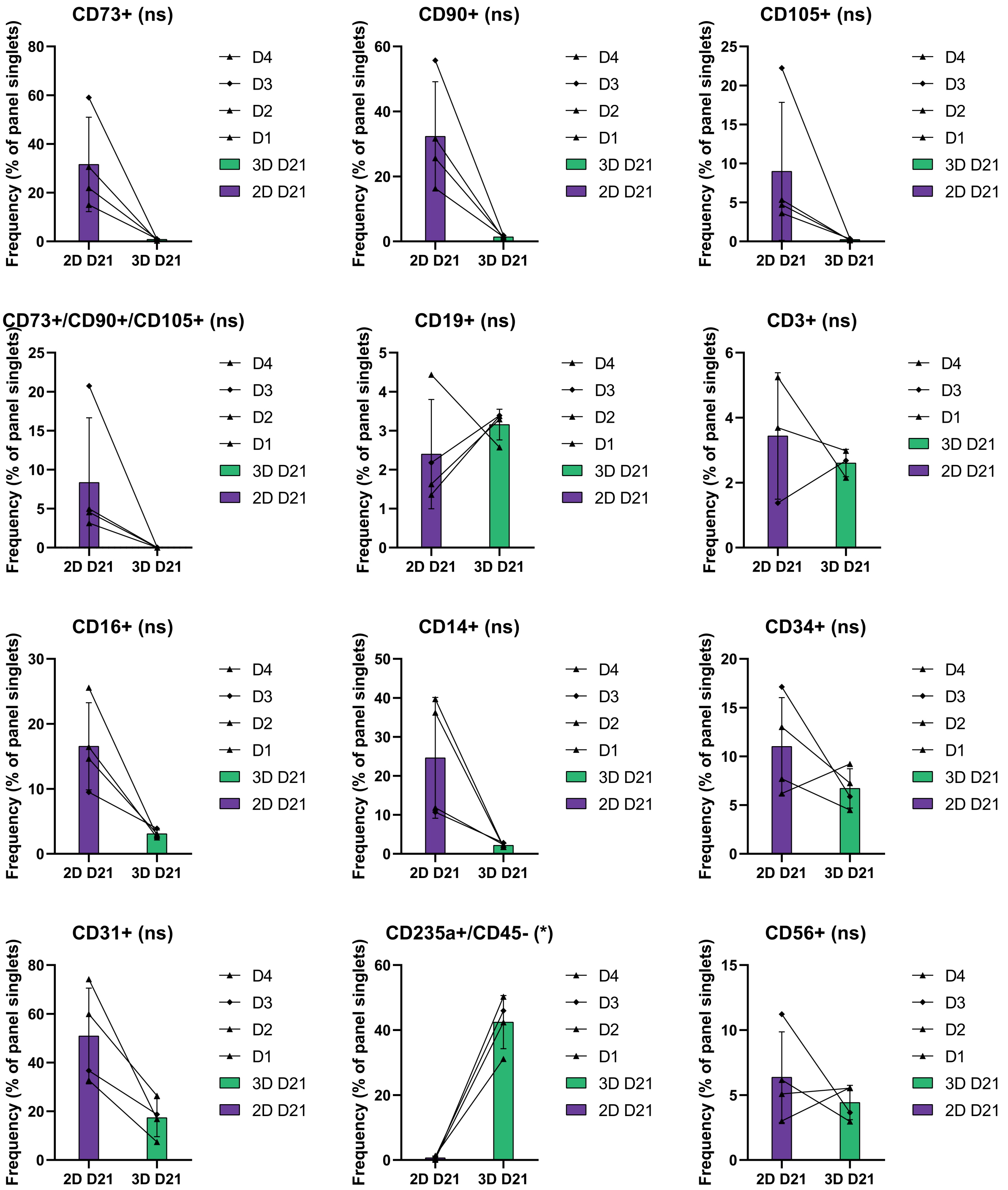
CD73+ (ns)
CD90+ (ns)
CD105+ (ns)
CD73+/CD90+/CD105+ (ns)
CD19+ (ns)
CD3+ (ns)
CD16+ (ns)
CD14+ (ns)
CD34+ (ns)
CD31+ (ns)
CD235a+/CD45- (*)
CD56+ (ns)
Frequency (% of panel singlets)
2D D21
3D D21
D4
D3
D2
D1

Supplementary Figure S3. Paired flow-cytometric comparisons of bulk GelMA and conventional 2D cultures. (A) Eleven paired comparisons between bulk 3D days 7 and 21. All comparisons were non-significant after Holm correction. Mean MSC-marker frequencies remained below 2%. (B) Twelve paired comparisons between 2D and bulk 3D at day 21. $CD235a^{+}/CD45^{-}$ frequency was higher in 3D after Holm correction ($p = 0.0286$). Other comparisons did not meet the adjusted significance threshold. Bars show mean ± SD, symbols show donors, and lines connect paired measurements. Black triangles denote D1, D2 and D4, and a black diamond denotes D3. Orange, green and purple identify 3D D7, 3D D21 and 2D D21, respectively. There are four complete pairs except for the CD3 endpoint comparison, which includes D1, D3 and D4 ($n = 3$). Two-sided paired t-tests were performed in GraphPad Prism 8.0.1. Holm correction was applied separately to the eleven temporal and twelve endpoint comparisons. ns denotes adjusted $p \geq 0.05$ and * denotes adjusted $p < 0.05$. Supplementary Table S4 reports exact summaries and p values. Readouts use panel-specific singlet denominators and are not additive cell-type proportions. Marker labels do not establish definitive cell identities.

#### Supplementary Figure S4

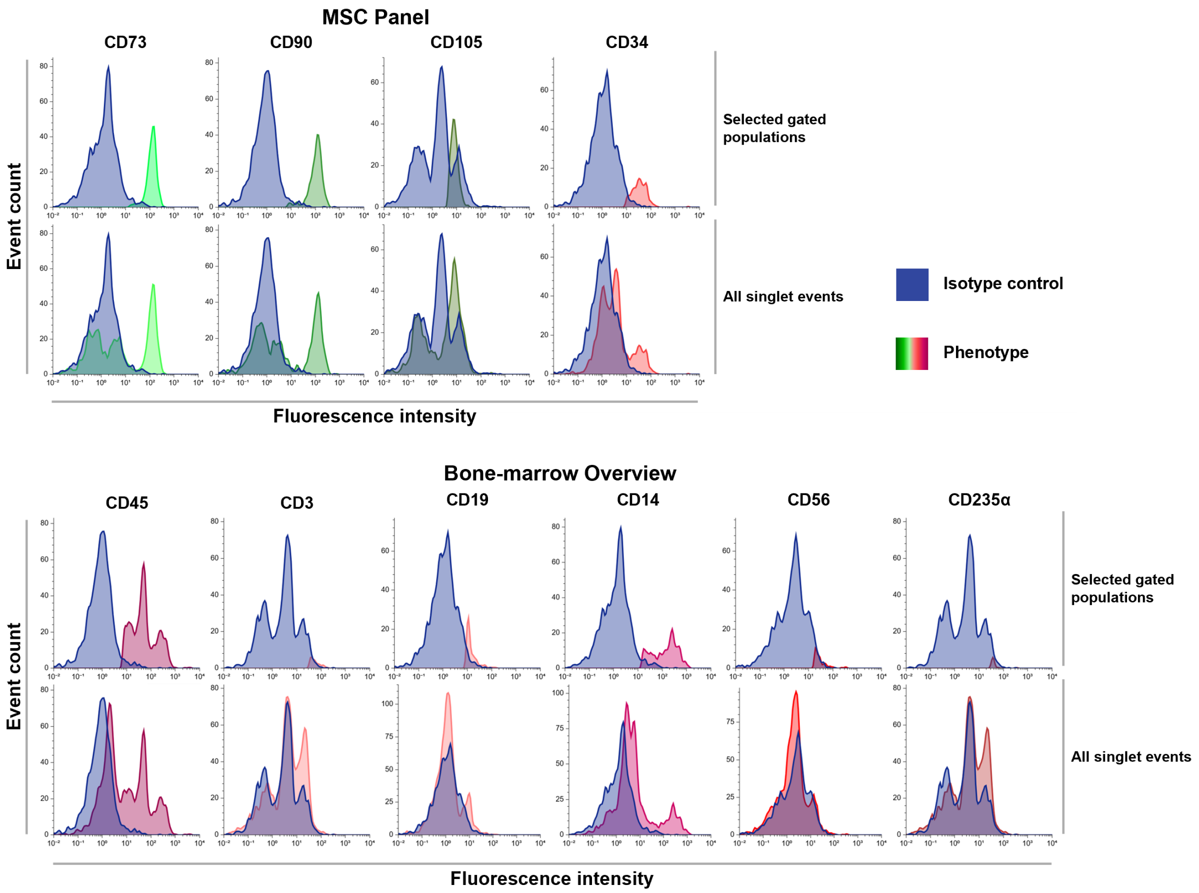


**Supplementary Figure S4. Fluorescence distributions and isotype controls for day-21 GMB flow cytometry.** The upper block shows the MSC panel and the lower block the bone-marrow overview panel from the exploratory single-donor experiment. Within each block, the upper row shows selected gated populations and the lower row the corresponding full singlet populations. Blue overlays indicate isotype controls, and coloured overlays indicate the respective antibody-stained populations. These descriptive distributions complement the marker-frequency readouts in Figure 2F. Antibody panels and gating hierarchies are detailed in Supplementary Table S5.

#### Supplementary Figure S5

**A**

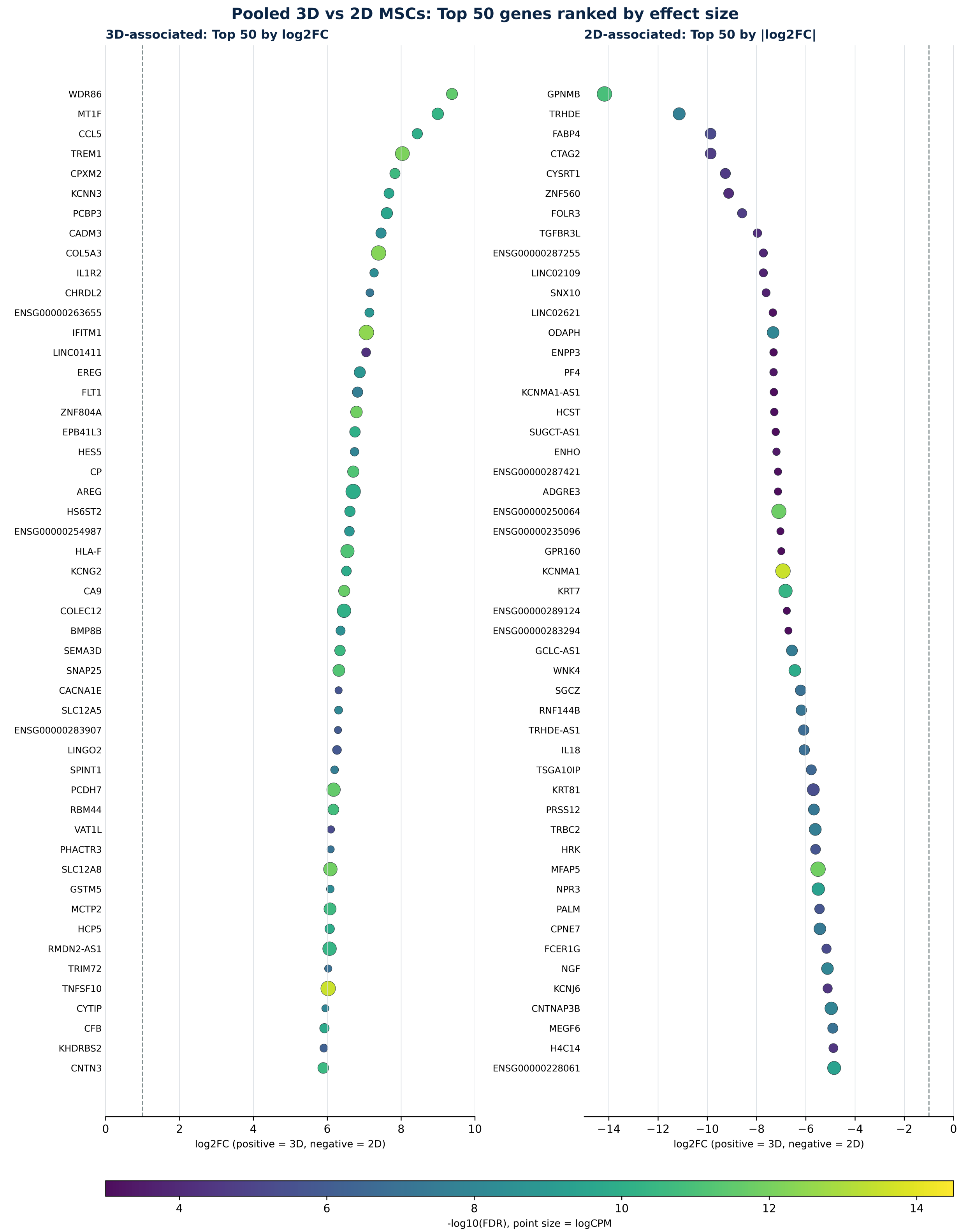

B

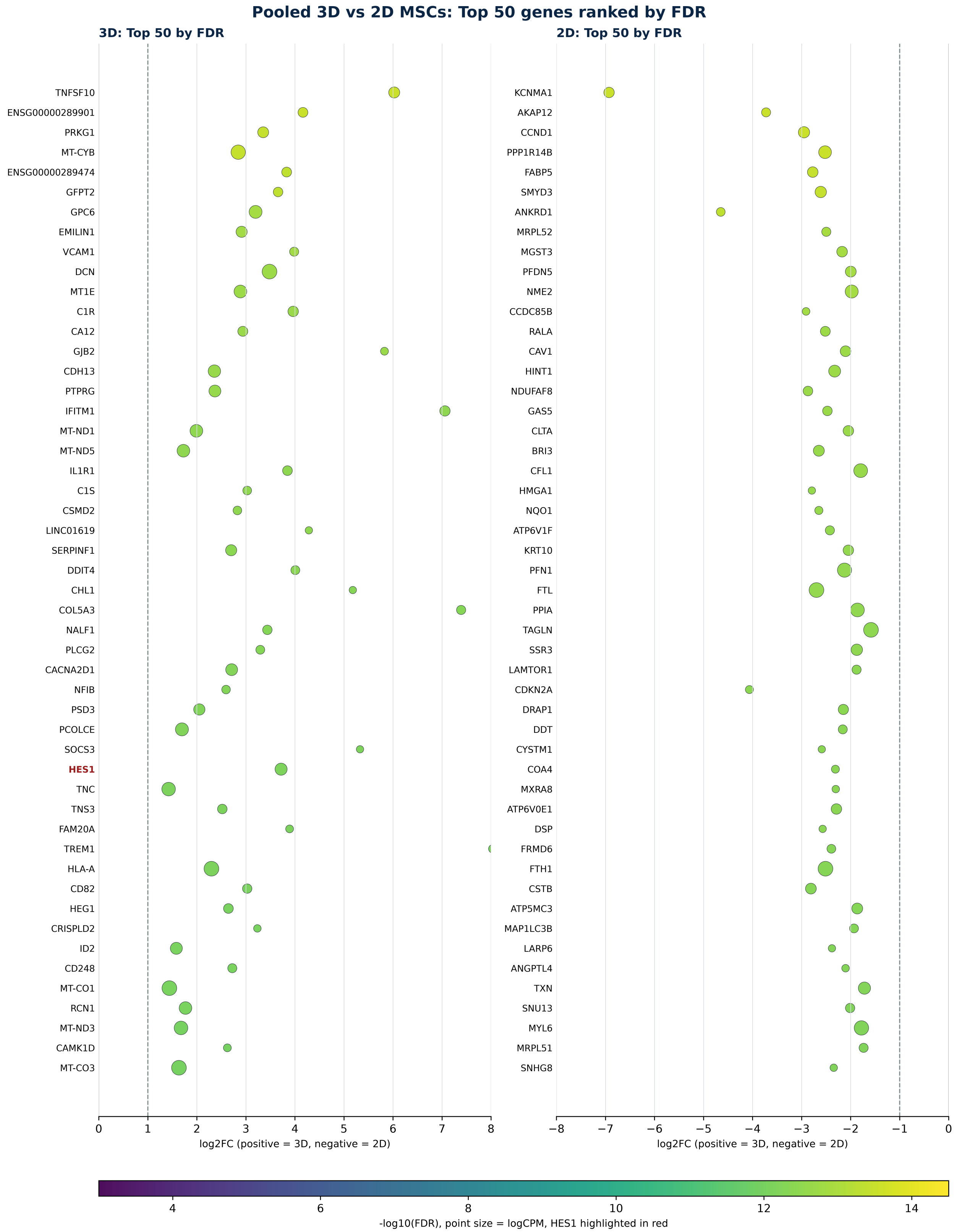

Pooled 3D vs 2D MSCs: Top 50 genes ranked by FDR
3D: Top 50 by FDR
2D: Top 50 by FDR
TNFSF10
ENSG00000289901
PRKG1
MT-CYB
ENSG00000289474
GFPT2
GPC6
EMILIN1
VCAM1
DCN
MT1E
C1R
CA12
GJB2
CDH13
PTPRG
IFITM1
MT-ND1
MT-ND5
IL1R1
C1S
CSMD2
LINC01619
SERPINF1
DDIT4
CHL1
COL5A3
NALF1
PLCG2
CACNA2D1
NFIB
PSD3
PCOLCE
SOCS3
HES1
TNC
TNS3
FAM20A
TREM1
HLA-A
CD82
HEG1
CRISPLD2
ID2
CD248
MT-CO1
RCN1
MT-ND3
CAMK1D
MT-CO3
KCNMA1
AKAP12
CCND1
PPP1R14B
FABP5
SMYD3
ANKRD1
MRPL52
MGST3
PFDN5
NME2
CCDC85B
RALA
CAV1
HINT1
NDUFAF8
GAS5
CLTA
BRI3
CFL1
HMGA1
NQO1
ATP6V1F
KRT10
PFN1
FTL
PPIA
TAGLN
SSR3
LAMTOR1
CDKN2A
DRAP1
DDT
CYSTM1
COA4
MXRA8
ATP6V0E1
DSP
FRMD6
FTH1
CSTB
ATP5MC3
MAP1LC3B
LARP6
ANGPTL4
TXN
SNU13
MYL6
MRPL51
SNHG8
0
1
2
3
4
5
6
7
8
log2FC (positive = 3D, negative = 2D)
−8
−7
−6
−5
−4
−3
−2
−1
0
log2FC (positive = 3D, negative = 2D)
4
6
8
10
12
14
-log10(FDR), point size = logCPM, HES1 highlighted in red

244 c

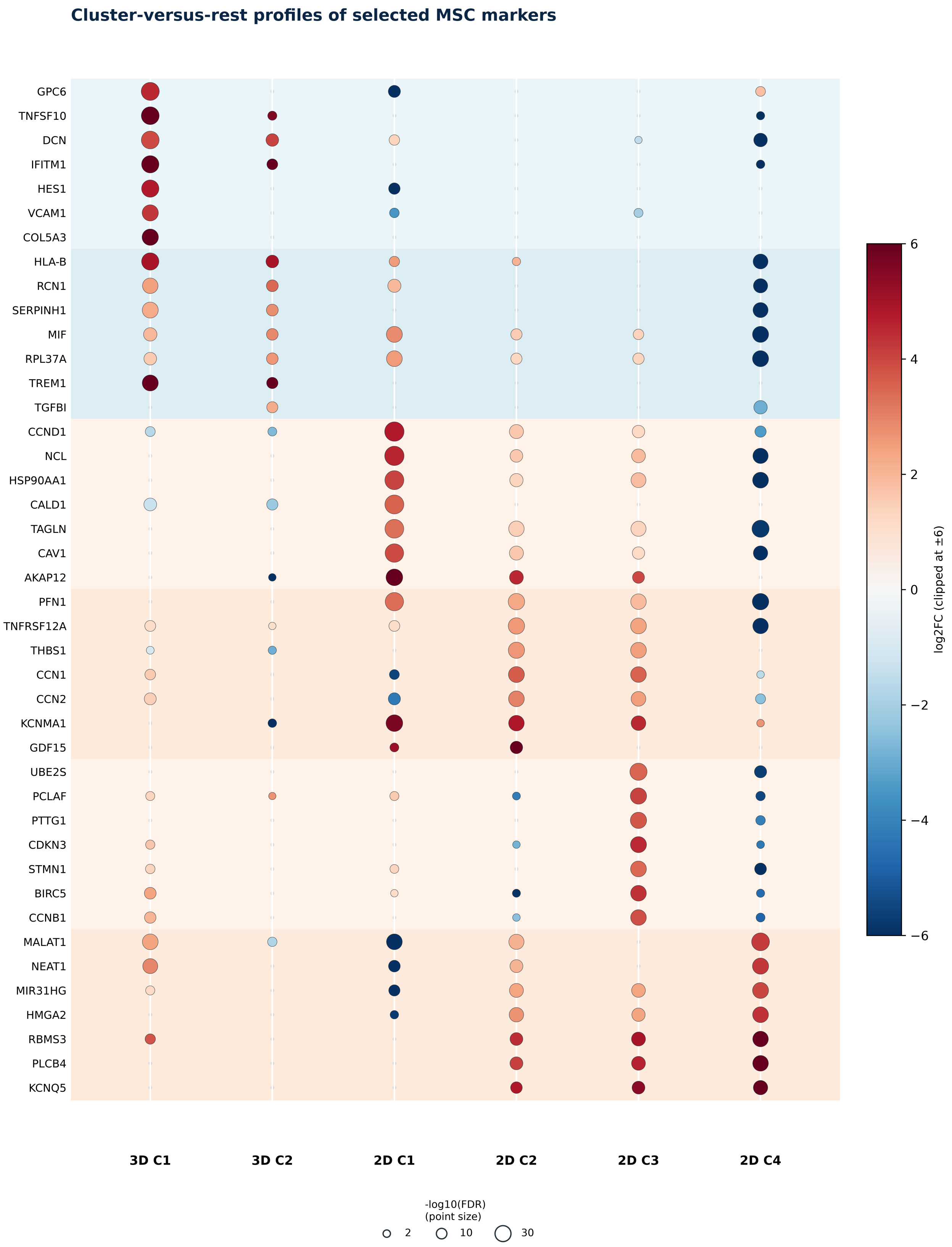

Cluster-versus-rest profiles of selected MSC markers
GPC6
TNFSF10
DCN
IFITM1
HES1
VCAM1
COL5A3
HLA-B
RCN1
SERPINH1
MIF
RPL37A
TREM1
TGFBI
CCND1
NCL
HSP90AA1
CALD1
TAGLN
CAV1
AKAP12
PFN1
TNFRSF12A
THBS1
CCN1
CCN2
KCNMA1
GDF15
UBE2S
PCLAF
PTTG1
CDKN3
STMN1
BIRC5
CCNB1
MALAT1
NEAT1
MIR31HG
HMGA2
RBMS3
PLCB4
KCNQ5
3D C1
3D C2
2D C1
2D C2
2D C3
2D C4
log2FC (clipped at ±6)
6
4
2
0
−2
−4
−6
-log10(FDR)
(point size)
2
10
30


245

Supplementary Figure S5. Extended differential-expression rankings and cluster-resolved marker profiles. (A) The top 50 pooled 3D-versus-2D genes per direction ranked by effect magnitude. (B) The top 50 pooled genes per direction ranked by FDR. In A and B, horizontal position indicates log2FC, colour indicates −log10(FDR), and point size indicates logCPM. HES1 is highlighted in red in B. (C) Cluster-versus-rest profiles for selected markers across 3D C1/C2 and 2D C1–C4. In C, colour denotes log2FC and point size denotes −log10(FDR), while small grey points do not meet the combined DEG threshold. The selected markers represent orthogonal biological axes—stromal/progenitor identity, ECM/adhesion, cytokine/immune response, hypoxia/glycolysis, mechanoadaptation and proliferation—rather than only the numerically strongest genes.

#### Supplementary Figure S6

##### A Progenitor/niche markers

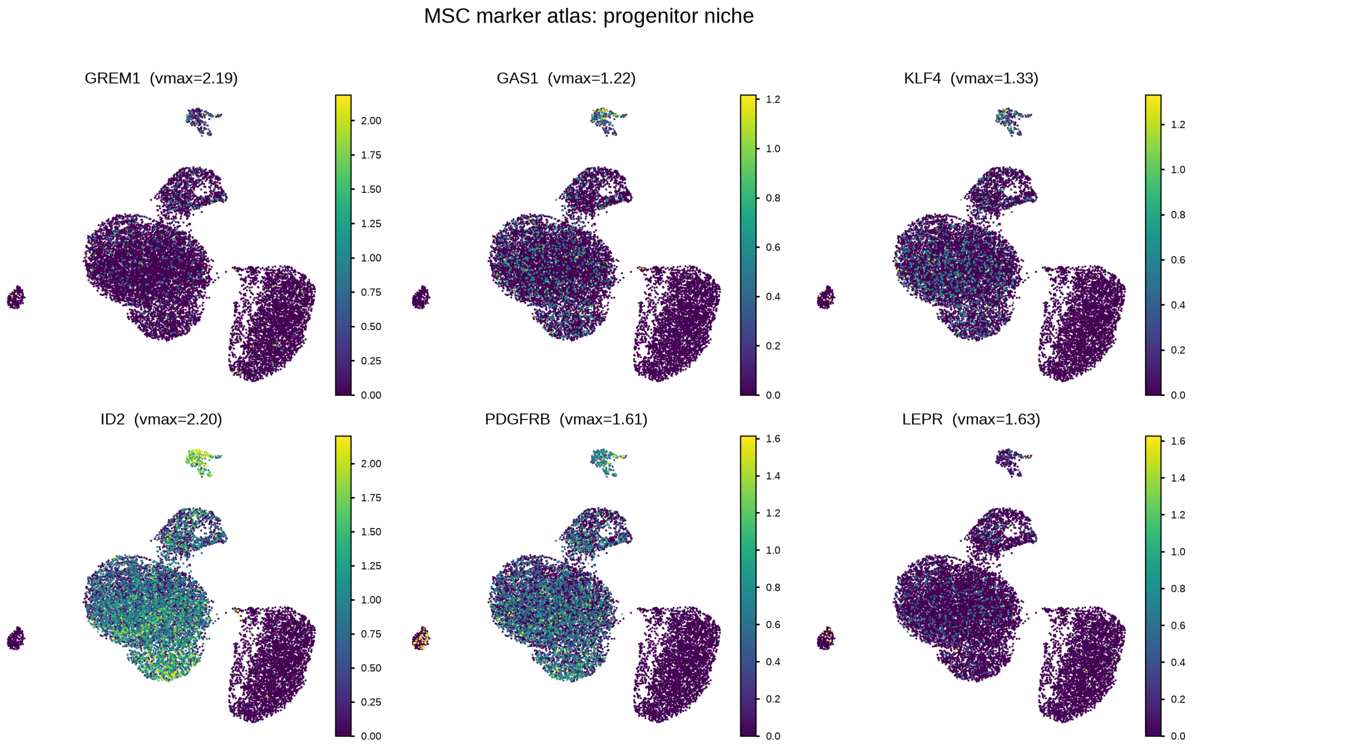


##### B Hypoxia/glycolysis markers

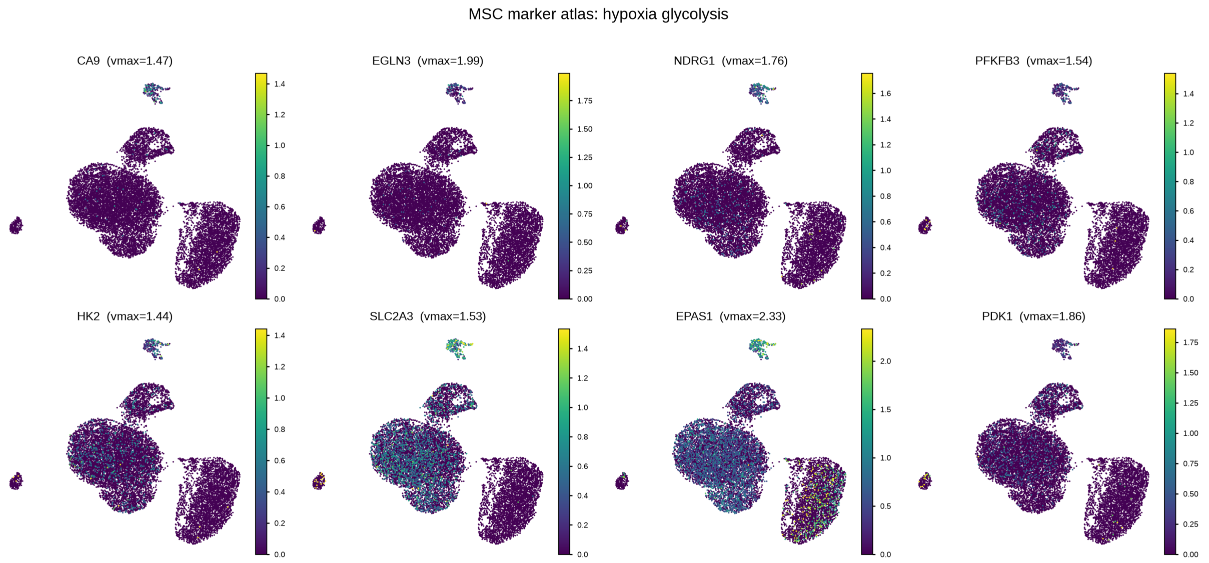

#### C Immune/ECM markers

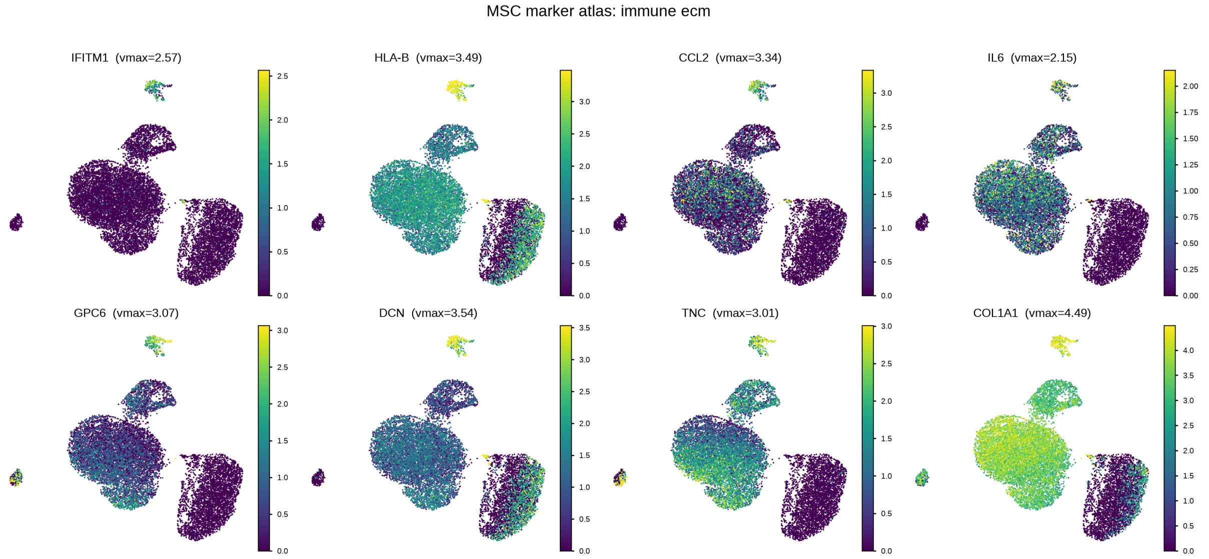


#### D 2D culture-adaptation markers

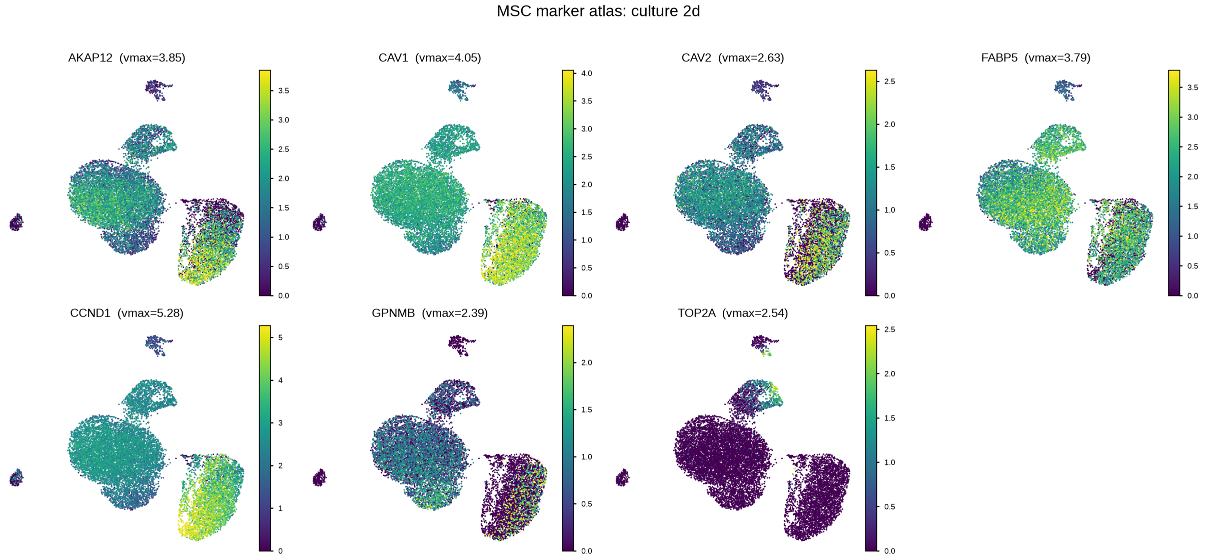


Supplementary Figure S6. Extended MSC marker atlas. (A) Progenitor/niche-associated genes. (B) Hypoxia/glycolysis-associated genes. (C) Immune-/ECM-associated genes. (D) 2D culture-adaptation genes. Feature-map colour limits were defined per gene and kept identical across the six clusters for that gene. Scales should not be compared between different genes or directly between this atlas and Figure 3E, which currently uses different displayed limits for some genes. The atlas localizes pooled changes to specific clusters and supports the coordinated interpretation of 3D C1 and the heterogeneous 2D compartment.

### Supplementary Figure S7

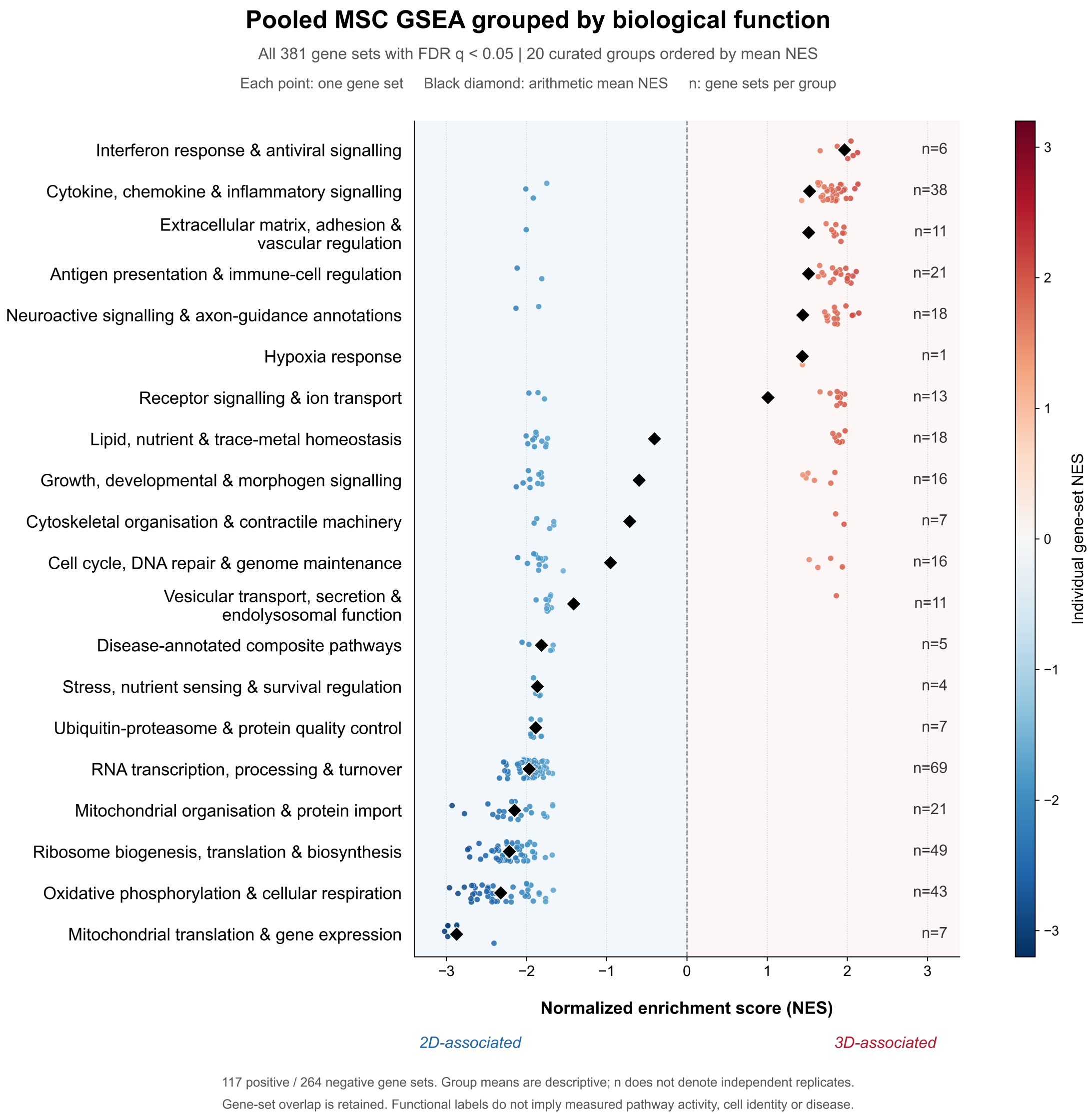


Supplementary Figure S7. Complete pooled GSEA landscape. All 381 gene sets with FDR q < 0.05 among 8,506 tested sets in the pooled 3D-versus-2D analysis. Sets from GO, Reactome, KEGG, MSigDB Hallmark and WikiPathways were assigned to 20 functional groups using annotations and, for ambiguous terms, leading-edge content. Each set was assigned to one primary group regardless of direction. Red and blue points show individual 3D- and 2D-associated NES values. Vertical jitter improves visibility and has no biological meaning. Black diamonds show unweighted arithmetic means of signed NES within groups, and n denotes gene-set counts, not biological replicates. Groups are ordered by their means. No additional NES or top-ranking filter was applied. Group means are descriptive and overlapping gene sets may share underlying genes.

Supplementary Figure S8

A

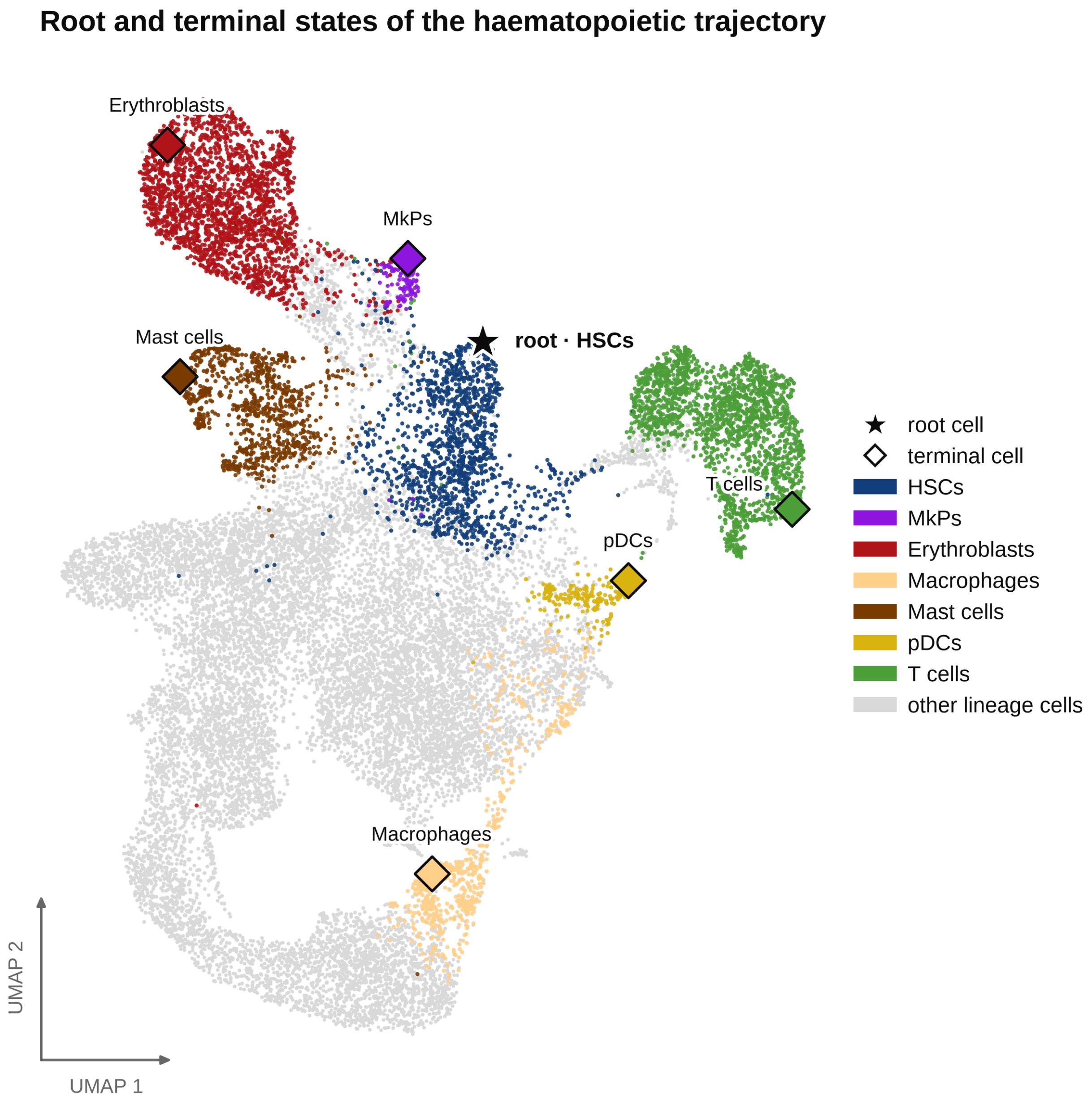

B

Palantir pseudotime and lineage trajectories · 21 901 cells

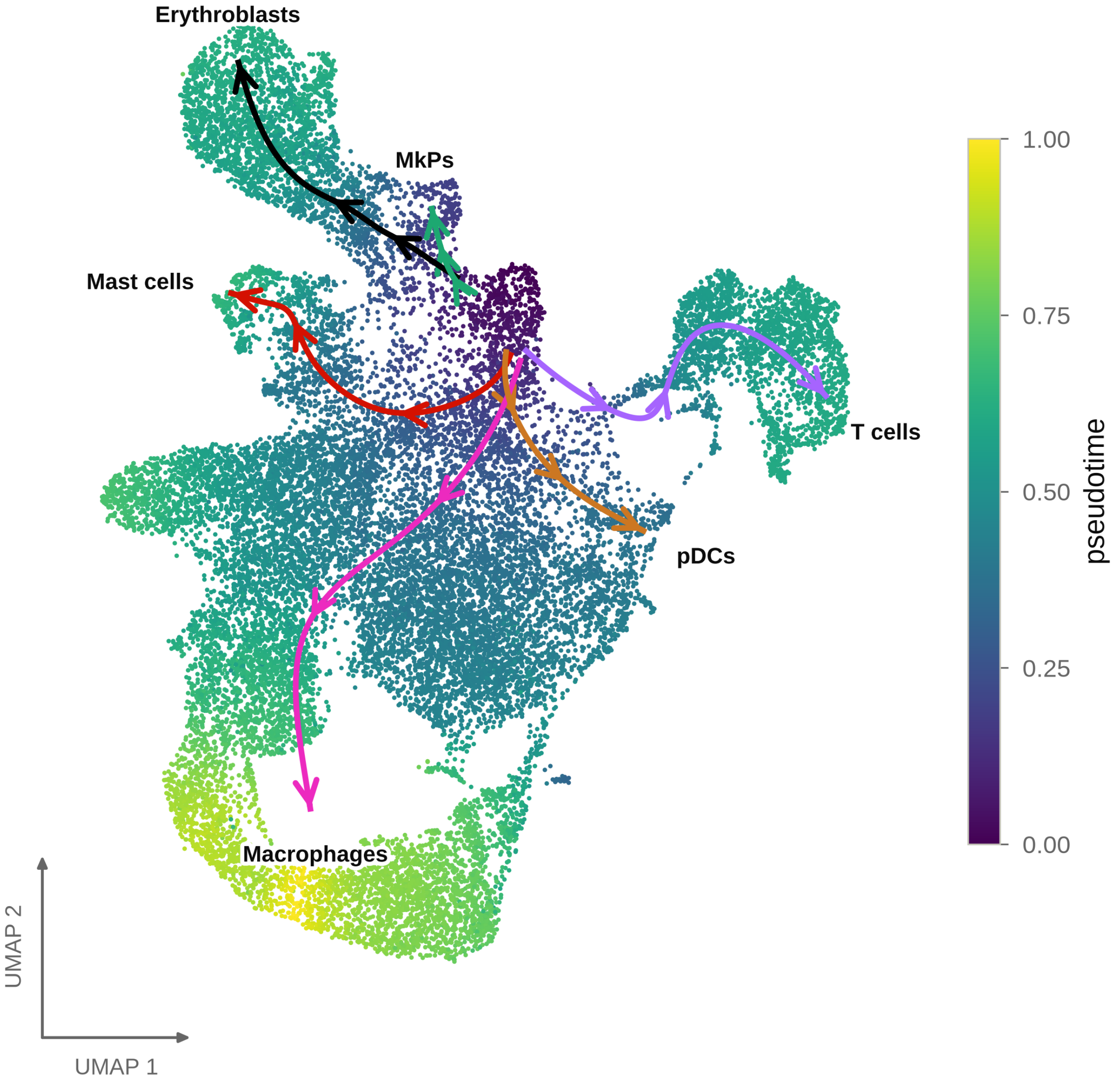

c

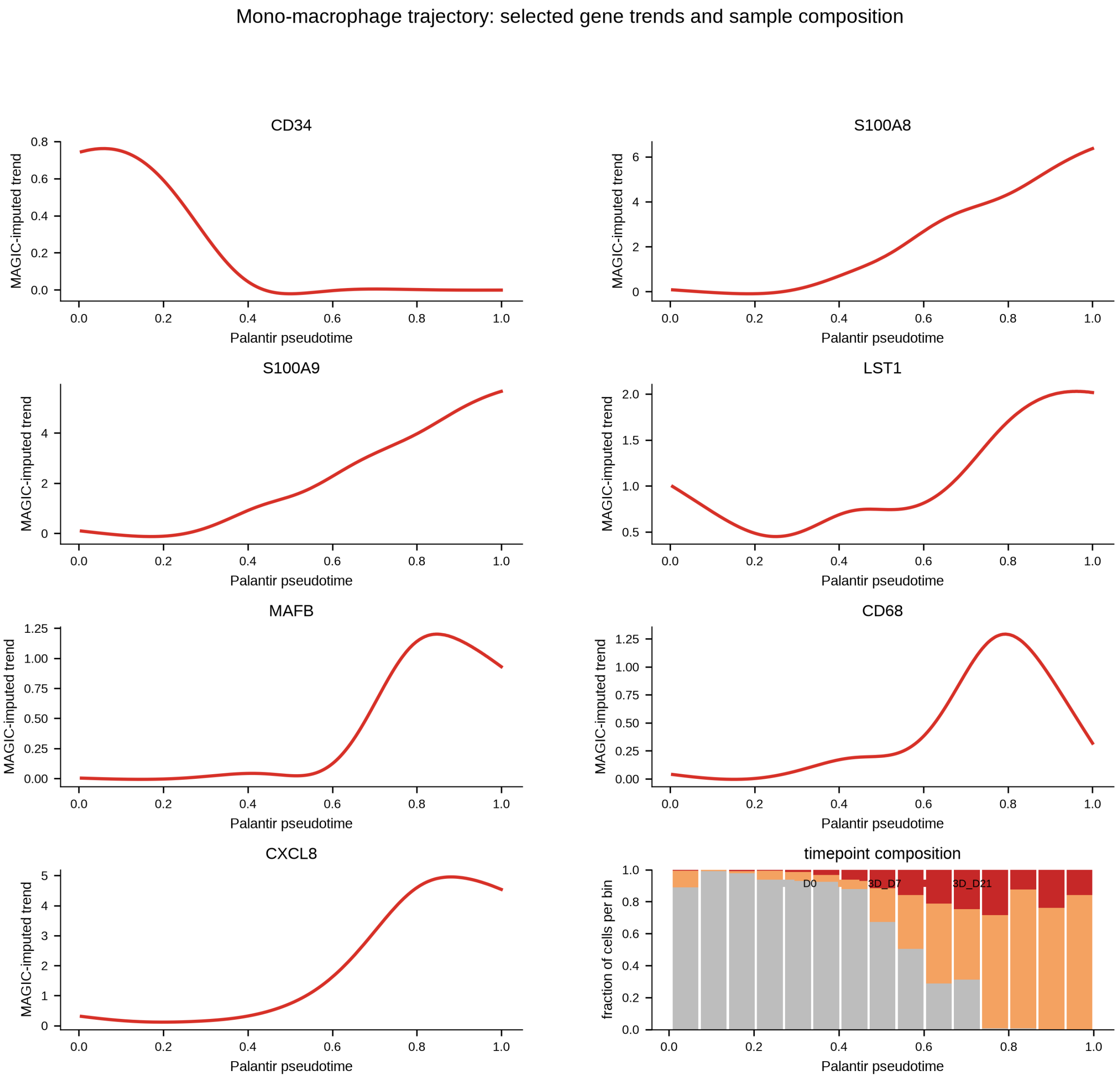


Supplementary Figure S8. Haematopoietic trajectory inference. (A) Computational HSC-proximal root and terminal states. (B) Palantir pseudotime and inferred lineage trajectories across 21,901 haematopoietic cells. (C) Monocyte/macrophage multi-gene trends and time-point composition. CD34 decreases along the inferred lineage, while S100A8, S100A9, LST1, MAFB, CD68 and CXCL8 occupy overlapping early-to-late myeloid regions. Pseudotime is a transcriptional ordering and must not be interpreted as direct culture time or experimental lineage tracing.

### Supplementary Figure S9

#### A Directed 3D-C1/macrophage candidates

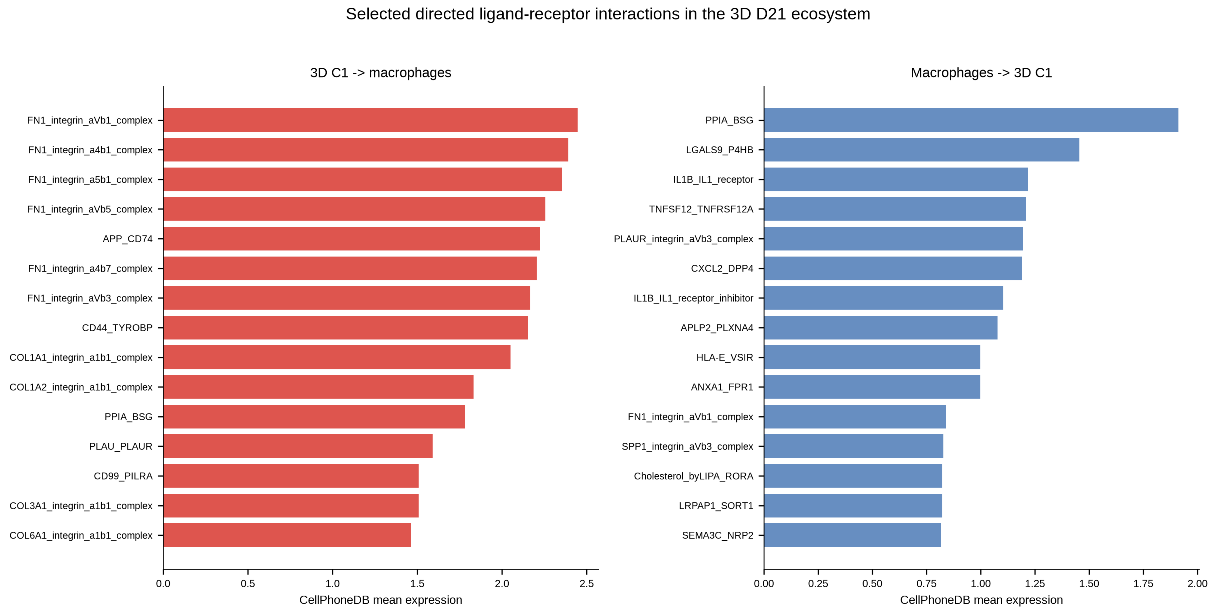

#### B Combined 23-cell-type context

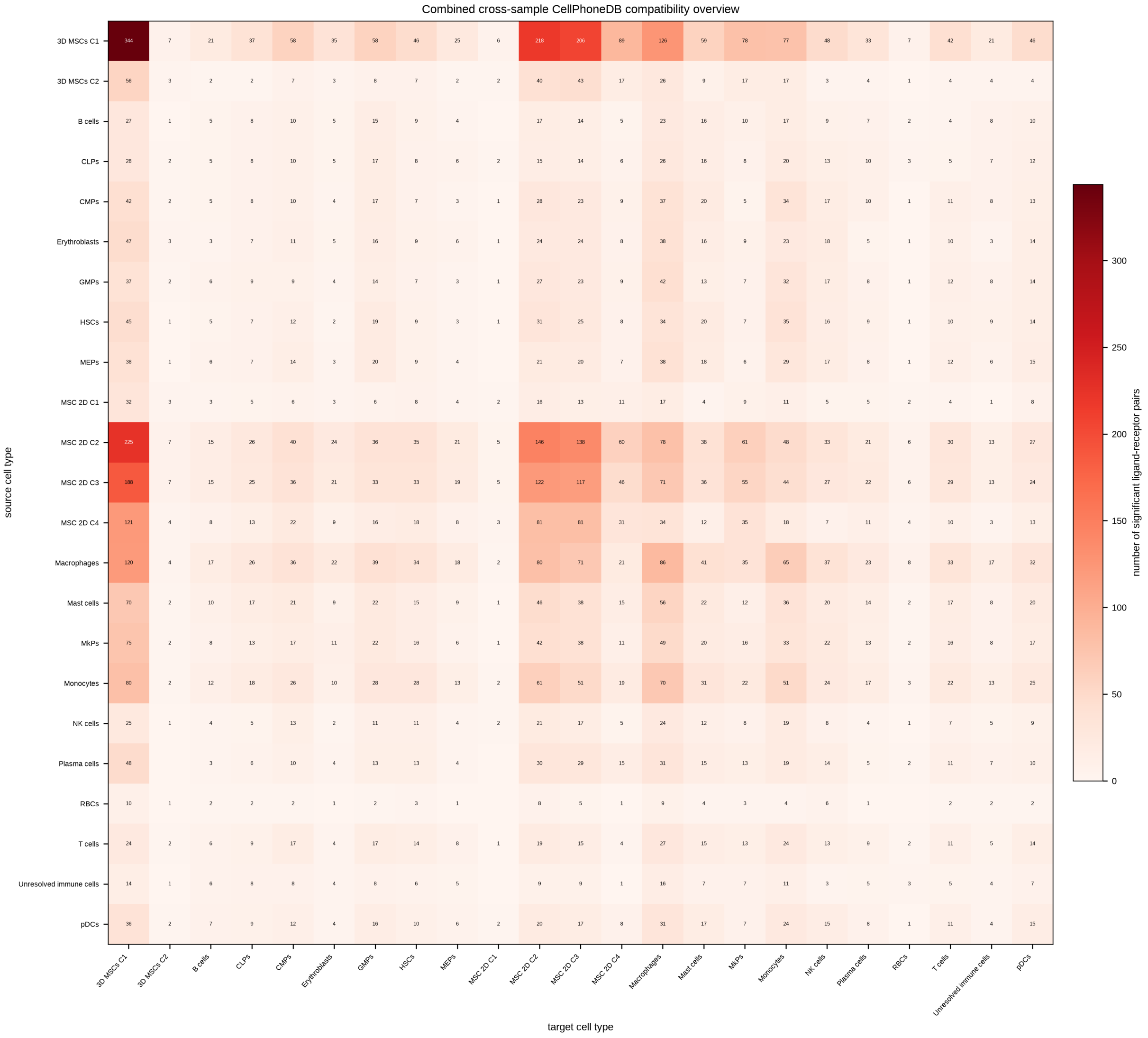

**C 2D day-21 context**

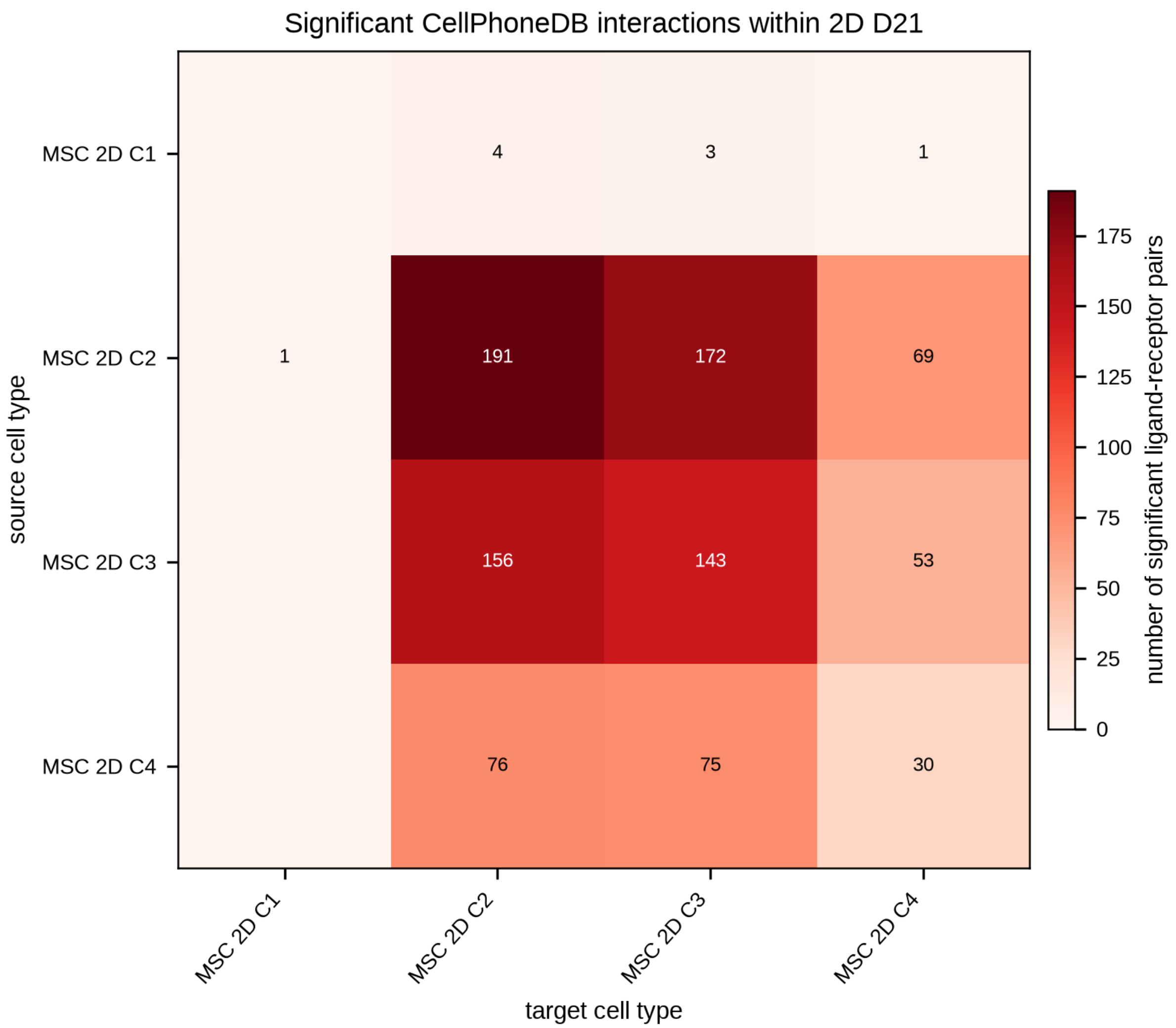


Supplementary Figure S9. Extended CellPhoneDB context. (A) Directed ligand–receptor candidates between 3D C1 and macrophages, including matrix/adhesion, proteolytic and inflammatory/reparative axes. (B) Combined 23-cell-type matrix shown as an expression-compatibility overview. Cell types that did not coexist in one sample must not be interpreted as direct communication. (C) 2D day-21 context matrix. Reported pair counts represent database-compatible expression and not communication strength.

### Supplementary Tables

#### Supplementary Table S1

Supplementary Table S1. GelMA batch definitions, independent 313-K DoF estimates and use in the project.

| Material | 313-K DoF | 0/0.3-Hz processing range | Project role |
|---|---|---|---|
| BATCH1 low DoF | 72.9% | 69.2–75.2% | Parent material and 1:1 blend component for medium-DoF bulk GelMA. |
| BATCH1 high DoF | 95.6% | 95.6–99.4% | Parent material and 1:1 blend component for medium-DoF bulk GelMA. High-DoF rheology formulation. |
| BATCH2 high DoF | 94.3% | 94.0–96.9% | High-DoF GelMA used for all GMB cultures. Characterized independently by NMR and cell-free rheology. |
| Medium-DoF GelMA | Not independently assigned | Not applicable | 1:1 BATCH1 low/high blend used for bulk cultures and medium-formulation rheology |

#### Supplementary Table S2

Supplementary Table S2. Final storage (G’) and loss (G”) moduli of cell-free 5% (w/v) GelMA formulations containing 0.2% (w/v) LAP. Values were obtained at the end of post-irradiation monitoring, as described in Supplementary Methods SM3, and represent one measurement run per formulation.

| Formulation | GelMA concentration | Material mapping | Endpoint G’ / G” (Pa) |
|---|---|---|---|
| Medium | 5% | 1:1 BATCH1 low/high | 174.3 / 1.533 |
| High | 5% | BATCH1 high DoF | 1924 / 1.569 |
| High | 5% | BATCH2 high DoF | 1376 / 3.318 |

#### Supplementary Table S3. Bulk flow cytometry

Antibody and control preparations for bulk flow cytometry. S3A–D document channels and dilutions in staining reactions. S3E describes gating sequences and denominators. Separately acquired panels provide marker-defined, partly overlapping frequencies.

##### S3A. isotype/control preparation

| Fluorochrome | Dilution |
|---|---|
| VioBlue | 1:50 |
| VioGreen | 1:50 |
| FITC | 1:50 |
| PE | 1:50 |
| PE-Vio615 | 1:50 |
| PE-Vio770 | 1:50 |
| APC | 1:1000 |
| APC-Vio770 | 1:50 |

##### S3B. Immune panel 1

| Marker/reagent | Fluorochrome/channel | Dilution |
|---|---|---|
| CD45 | VioBlue | 1:50 |
| CD4 | VioGreen | 1:50 |
| CD3 | FITC | 1:50 |
| CD56 | PE | 1:50 |
| 7-AAD | PerCP | 1:50 |
| CD19 | PE-Vio770 | 1:50 |
| CD73 | APC | 1:1000 |
| CD8 | APC-Vio770 | 1:50 |

##### S3C. Immune panel 2

| Marker | Fluorochrome | Dilution |
|---|---|---|
| CD45 | PE-Vio615 | 1:50 |
| CD235a | FITC | 1:50 |
| CD16 | PE | 1:50 |
| CD14 | APC | 1:50 |

| Marker | Fluorochrome | Dilution |
|---|---|---|
| CD31 | APC-Vio770 | 1:50 |

#### S3D. MSC panel

| Marker | Fluorochrome | Dilution |
|---|---|---|
| CD105 | VioBlue | 1:50 |
| HLA-DR | VioGreen | 1:50 |
| CD14 | FITC | 1:50 |
| CD19 | PE | 1:50 |
| CD45 | PE-Vio615 | 1:50 |
| CD34 | PE-Vio770 | 1:50 |
| CD73 | APC | 1:1000 |
| CD90 | APC-Vio770 | 1:50 |

**S3E. gating sequence, denominators and interpretation**

| Readout | Gating sequence | Panel | Denominator / interpretation |
|---|---|---|---|
| Common input | FSC/SSC cell gate → two sequential singlet gates | Respective panel | Singlets in the respective panel |
| CD73, CD90 or CD105 individually | Each marker⁺ → CD45⁻ → HLA-DR⁻ → CD34⁻ | MSC | MSC-panel singlets, MSC-associated |
| Combined CD73⁺/CD90⁺/CD105⁺ | CD105⁺ → CD90⁺ → CD73⁺ → CD45⁻ → HLA-DR⁻ → CD34⁻ | MSC | MSC-panel singlets, combined MSC readout |
| CD34⁺ | MSC-panel singlets → CD34⁺ (PE-Vio770), without a CD45 restriction | MSC | MSC-panel singlets, haematopoietic progenitor-associated |
| CD19⁺ | Immune-panel-1 singlets → CD19⁺ (PE-Vio770), without a CD45 restriction | Immune panel 1 | Panel singlets, B-cell-associated |
| CD3⁺/CD45⁺ | CD45⁺ → CD3⁺ | Immune panel 1 | Panel singlets, T-cell-associated |
| CD56⁺ | Immune-panel-1 singlets → CD56⁺ (PE), without a CD45 restriction | Immune panel 1 | Panel singlets, NK-cell-associated |
| CD14⁺/CD45⁺ or CD16⁺/CD45⁺ | CD45⁺ → CD14⁺ or CD16⁺ | Immune panel 2 | Panel singlets, myeloid-associated |
| Total CD31⁺ | Immune-panel-2 singlets → CD31⁺, without a CD45 restriction | Immune panel 2 | Immune-panel-2 singlets. CD31 association does not establish endothelial identity. |
| CD235a⁺/CD45⁻ | CD45⁻ → CD235a⁺ | Immune panel 2 | Panel singlets, erythroid-associated |

Frequencies refer to singlets in the respective panel, not the immediately preceding gate. Separate panels and partly overlapping readouts must not be summed into a cellular composition. Cell-type associations are based on the listed marker combinations and do not establish functional identity.

#### Supplementary Table S4

Supplementary Table S4. Paired bulk-flow-cytometry summaries calculated from donor-level event counts and panel-specific singlet denominators. Values are mean ± SD. S4A contains eleven temporal comparisons and S4B twelve day-21 comparisons. Raw two-sided paired t-test p values and Holm-adjusted p values are shown. Holm correction was applied separately within each comparison family. CD3 in S4B uses D1, D3 and D4 only because the 2D D21 observation for D2 was missing. Consequently, the CD3 3D-D21 mean in S4B differs from its four-donor mean in S4A and the condition overview. No missing values were replaced with zero. Statistical annotations refer to adjusted p values. Readouts are marker-defined and are not additive population proportions.

**S4A Paired 3D day 7 versus day 21 comparisons**

| Readout | Pairs | 3D D7 (%) | 3D D21 (%) | Raw p | Holm p |
|---|---|---|---|---|---|
| CD73$^+$ | 4 | 1.03 ± 0.36 | 0.86 ± 0.36 | 0.1585 | 1.0000 |
| CD90$^+$ | 4 | 1.50 ± 0.67 | 1.38 ± 0.39 | 0.7227 | 1.0000 |
| CD105$^+$ | 4 | 0.46 ± 0.11 | 0.25 ± 0.10 | 0.0841 | 0.8413 |
| CD19$^+$ | 4 | 2.86 ± 0.31 | 3.16 ± 0.39 | 0.1624 | 1.0000 |
| CD3$^+$/CD45$^+$ | 4 | 2.36 ± 0.10 | 2.90 ± 0.68 | 0.2418 | 1.0000 |
| CD16$^+$/CD45$^+$ | 4 | 2.79 ± 0.67 | 3.08 ± 0.60 | 0.2265 | 1.0000 |
| CD14$^+$/CD45$^+$ | 4 | 1.68 ± 0.82 | 2.19 ± 0.51 | 0.0684 | 0.7527 |
| CD34$^+$ | 4 | 6.21 ± 0.51 | 6.71 ± 2.02 | 0.6039 | 1.0000 |
| Total CD31$^+$ | 4 | 19.44 ± 10.05 | 17.36 ± 7.80 | 0.4811 | 1.0000 |
| CD235a$^+$/CD45$^-$ | 4 | 41.36 ± 13.50 | 42.43 ± 8.21 | 0.7407 | 1.0000 |
| CD56$^+$ | 4 | 3.96 ± 0.16 | 4.43 ± 1.32 | 0.5529 | 1.0000 |

**S4B Paired 2D versus 3D day 21 comparisons**

| Readout | Pairs | 2D D21 (%) | 3D D21 (%) | Raw p | Holm p |
|---|---|---|---|---|---|
| CD73$^+$ | 4 | 31.62 ± 19.36 | 0.86 ± 0.36 | 0.0505 | 0.4036 |
| CD90$^+$ | 4 | 32.34 ± 16.80 | 1.38 ± 0.39 | 0.0335 | 0.3260 |
| CD105$^+$ | 4 | 8.98 ± 8.86 | 0.25 ± 0.10 | 0.1417 | 0.8259 |
| CD73$^+$CD90$^+$CD105$^+$ | 4 | 8.35 ± 8.29 | 0.007 ± 0.010 | 0.1376 | 0.8259 |
| CD19$^+$ | 4 | 2.40 ± 1.40 | 3.16 ± 0.39 | 0.4564 | 1.0000 |
| CD3$^+$/CD45$^+$ | 3 | 3.44 ± 1.95 | 2.60 ± 0.42 | 0.5805 | 1.0000 |
| CD16$^+$/CD45$^+$ | 4 | 16.54 ± 6.71 | 3.08 ± 0.60 | 0.0326 | 0.3260 |
| CD14$^+$/CD45$^+$ | 4 | 24.61 ± 15.50 | 2.19 ± 0.51 | 0.0674 | 0.4718 |
| CD34$^+$ | 4 | 11.01 ± 5.02 | 6.71 ± 2.02 | 0.2429 | 0.9717 |
| Total CD31$^+$ | 4 | 50.89 ± 19.63 | 17.36 ± 7.80 | 0.0296 | 0.3251 |
| CD235a$^+$/CD45$^-$ | 4 | 0.68 ± 0.57 | 42.43 ± 8.21 | 0.0024 | 0.0286 |
| CD56$^+$ | 4 | 6.37 ± 3.49 | 4.43 ± 1.32 | 0.4467 | 1.0000 |

#### Supplementary Table S5. GMB flow cytometry

Antibody preparations and gating hierarchies for the single-donor GMB pilot on day 21. The MSC panel contained 1,263 singlet events and the bone-marrow overview panel 2,001. Frequencies in Figure 2F use these respective denominators. Corresponding fluorescence histograms and isotype controls are shown in Supplementary Figure S4. Components of a shared cocktail are measured in one channel and cannot be resolved individually from that signal. Dilutions refer to the staining reaction.

##### S5A. isotype/control preparation

| Fluorochrome | Dilution |
|---|---|
| VioBlue | 1:50 |
| VioGreen | 1:50 |
| FITC | 1:50 |
| PE | 1:50 |
| PE-Vio615 | 1:50 |
| PE-Vio770 | 1:50 |
| APC | 1:1000 |
| APC-Vio770 | 1:50 |

##### S5B. MSC panel

| Marker/reagent | Fluorochrome/channel | Dilution |
|---|---|---|
| CD105 | VioBlue | 1:50 |
| CD3 / CD14 / CD235a | FITC cocktail | 1:50 |
| CD56 / CD16 / CD11b | PE cocktail | 1:50 |
| CD19 / CD45 / HLA-DR | PE cocktail (same channel) | 1:50 |
| 7-AAD | PerCP | 1:50 |
| CD34 | PE-Vio770 | 1:50 |
| CD73 | APC | 1:1000 |
| CD90 | APC-Vio770 | 1:50 |

##### S5C. Bone-marrow overview

| Marker/reagent | Fluorochrome/channel | Dilution |
|---|---|---|
| HLA-DR | VioGreen | 1:50 |
| CD3 / CD235a | FITC cocktail | 1:50 |
| CD56 | PE | 1:50 |

| Marker/reagent | Fluorochrome/channel | Dilution |
|---|---|---|
| 7-AAD | PerCP | 1:50 |
| CD19 | PE-Vio770 | 1:50 |
| CD14 | APC | 1:50 |
| CD45 | APC-Vio770 | 1:50 |

#### S5D. Gating hierarchies and denominators

| Readout | Gating sequence | Denominator | Interpretation |
| --- | --- | --- | --- |
| SEF (stromal enrichment fraction) | FITC$^{-}$ / PE$^{-}$ / PE-Vio770$^{-}$ | MSC-panel singlets | Negative for CD3, CD14, CD235a, CD56, CD16, CD11b, CD19, CD45, HLA-DR and CD34 |
| CD73, CD90 or CD105 individually | SEF → APC$^{+}$ / APC-Vio770$^{+}$ / VioBlue$^{+}$ | MSC-panel singlets | Individual MSC-associated readouts |
| Combined CD73$^{+}$/CD90$^{+}$/CD105$^{+}$ | SEF → CD73$^{+}$CD90$^{+}$CD105$^{+}$ | MSC-panel singlets | Combined MSC readout |
| CD34$^{+}$/PE-cocktail$^{+}$ | PE-Vio770$^{+}$ → FITC$^{-}$ → PE$^{+}$ | MSC-panel singlets | CD34$^{+}$ with positive PE exclusion cocktail, CD45 alone cannot be resolved |
| CD45 | APC-Vio770$^{+}$ | Overview-panel singlets | Haematopoietic-associated |
| CD3 | CD45$^{+}$ → FITC$^{+}$ | Overview-panel singlets | T-cell-associated, FITC also contains CD235a |
| CD19 / CD14 / CD56 | CD45$^{+}$ → PE-Vio770$^{+}$ / APC$^{+}$ / PE$^{+}$ | Overview-panel singlets | B-cell-/myeloid-/NK-cell-associated |
| CD235a$^{+}$/CD45$^{-}$ | CD45$^{-}$ → FITC$^{+}$ | Overview-panel singlets | Erythroid-associated, FITC also contains CD3 |

SEF is an operational stromal enrichment fraction rather than evidence of purity. Individual and combined gates overlap. Frequencies are calculated as 100 × events/singlets in the respective panel. The pilot was analysed without inferential statistics. CD34$^{+}$/PE-cocktail$^{+}$ denotes CD34-positive, FITC-cocktail-negative and PE-cocktail-positive events, as detailed in S5D. Independent CD45 positivity cannot be inferred from the PE cocktail.

#### Supplementary Table S6. Single-cell samples, sequencing records and analysis parameters

Four samples from one donor. S6A describes sample preparation and final analysed cell counts. S6B reports cell loading, viability and mean sequencing depth. S6C provides total read counts and Cell Ranger cell estimates. S6D identifies the Palantir root and six terminal cells. S6E documents cluster-specific QC and whitelisting. S6F lists the annotation gene signatures. S6G reports the scVI hyperparameter search space. The sorting target of 50,000 7-AAD-negative cells, the number of cells loaded, the Cell Ranger cell estimate and the final analysed cell count are distinct quantities. Final analysed cell counts do not quantify absolute isolation yield.

##### S6A. Preparation and final cell counts

| Sample | Preparation/culture | Final cells | MSC-annotated | Interpretation |
|---|---|---|---|---|
| D0 | Density gradient, CD45/CD235a negative selection, uncultured | 14,387 | 0 | Selectively processed, not an unbiased baseline |
| 3D D7 | Erythrocyte lysis, GMB culture, 5% $O_2$ | 7,389 | 0 | Heterogeneous marrow population |
| 3D D21 | Erythrocyte lysis, GMB culture, 5% $O_2$ | 3,138 | 305 | 3D C1: 245, 3D C2: 60 |
| 2D D21 | Conventional adherent culture, 5% $O_2$ | 12,870 | 12,870 | 2D C1: 3,674, C2: 7,788, C3: 1,216, C4: 192 |
| Total | Four samples from one donor | 37,784 | 13,175 | Representation in the final object, not absolute yield |

##### S6B Cell loading, viability and sequencing depth

| Sample | Cells loaded | Viability | Mean reads per cell |
|---|---|---|---|
| D0 | 27,000 | "good", not numerical | 29,732 |
| 3D D7 | 20,000 | 91% | 45,101 |
| 2D D21 | 15,970 | 98% | 23,521 |
| 3D D21 | 20,000 | 95% | 75,316 |

D0 negative selection and cell loading differed between samples. Differences in recovery, capture, transcript quality and filtering may also affect final cell representation. Loaded cell numbers do not establish equal MSC input or absolute isolation yield. Sequencing depth is reported as mean reads per cell in the sample-level Cell Ranger summaries and is distinct from the final analysed cell counts in S6A.

##### S6C Total sequencing reads and estimated cells

| Sample | Total reads | Cell Ranger estimated cells |
|---|---|---|
| D0 | 490,520,902 | 16,498 |

| Sample | Total reads | Cell Ranger estimated cells |
|---|---|---|
| 3D D7 | 435,903,951 | 9,665 |
| 2D D21 | 262,073,658 | 11,142 |
| 3D D21 | 325,063,284 | 4,316 |

#### S6D Palantir root and terminal cells

| Role and annotation | Sample | Cell identifier |
|---|---|---|
| Root / HSCs | D0 | CGGGTGAGTTGGCGTC-1-MNC_nat_D0-D0 |
| Terminal / erythroblasts | D0 | CCAACTAAGTCGAACC-1-MNC_nat_D0-D0 |
| Terminal / MkPs | D0 | CCAACCCTCAGGTGAG-1-MNC_nat_D0-D0 |
| Terminal / macrophages | 3D D21 | GCTCAGTAGGATGGCA-1-MNC_3D_D21-3D_D21 |
| Terminal / mast cells | 3D D7 | ACAGCTATCTAACTCC-1-MNC_3D_D7-3D_D7 |
| Terminal / pDCs | D0 | CCACGGATCATGCTGA-1-MNC_nat_D0-D0 |
| Terminal / T cells | 3D D7 | CAGTTCAAGCTAGCAC-1-MNC_3D_D7-3D_D7 |

#### S6E Cluster-specific quality control and whitelisting

Whitelisted clusters flagged by the global QC filter and their handling after per-cluster MAD filtering. Flagged % is given for the global filter (Glob.) and after per-cluster MAD (MAD). mt med. is the median mitochondrial read fraction (%). Mode: A, automatic per-cluster MAD (flagged rate falls below 20% without whitelisting); S, soft whitelist (>20% still flagged locally, n ≥ 150, so per-cluster MAD is still applied and true outliers are removed); H, hard whitelist (n < 150, per-cluster MAD unreliable, all cells kept). Low mt in all clusters indicates that global flagging reflects cell-type biology (contractile/ECM genes, immunoglobulin secretion, granule genes, hemoglobin) rather than damage. Cluster 45 is borderline (mt P90 7.95%, hard limit 8%).

| Cl. | Cell type | n | Mode | Glob. | MAD | Kept, n (%) | mt med. |
|---|---|---|---|---|---|---|---|
| 1 | Fibro/myofibroblast | 2419 | A | 89.5 | 14.5 | 2069 (85.5) | 1.23 |
| 22 | Fibroblast | 788 | S | 98.9 | 34.3 | 518 (65.7) | 1.48 |
| 28 | Promyelocyte | 693 | S | 77.2 | 23.5 | 530 (76.5) | 2.34 |
| 34 | Myofibroblast | 596 | S | 91.4 | 32.6 | 402 (67.4) | 0.89 |
| 35 | Fibroblast | 583 | S | 55.6 | 23.3 | 447 (76.7) | 0.83 |
| 36 | IgA plasma cells | 560 | S | 95.5 | 24.3 | 424 (75.7) | 0.94 |
| 42 | IgG plasma cells | 395 | S | 95.2 | 22.0 | 308 (78.0) | 0.84 |
| 44 | Fibroblast, cycling | 317 | S | 90.9 | 29.0 | 225 (71.0) | 1.11 |

| Cl. | Cell type | n | Mode | Glob. | MAD | Kept, n (%) | mt med. |
|---|---|---|---|---|---|---|---|
| 45 | Fibroblast | 316 | A | 75.3 | 13.9 | 272 (86.1) | 3.35 |
| 57 | Erythroid | 94 | H | 100 | 74.5 | 94 (100) | 0.11 |
| 58 | Fibroblast | 86 | H | 100 | 73.3 | 86 (100) | 1.53 |

#### S6F Cell-type-specific marker gene signatures

Cell-type-specific marker gene signatures used for cell-type annotation. Gene symbols are grouped by their corresponding hematopoietic, immune, stromal, vascular, and bone-associated cell populations.

| Cell type | Marker genes |
|---|---|
| Hematopoietic stem/progenitor cells | CD34, PROM1, KIT, FLT3, MPL, THY1, AVP, HLF, CRHBP, MECOM, GATA2, MEIS1, SPINK2, CYTL1, MLLT3, HOXA9, RUNX1, MYB, SOX4, CD38 |
| B-cell lineage | CD19, MS4A1, CD79A, CD79B, CD74, MZB1, IGHM, IGHD, IGKC, IGLC2, CD22, PAX5, VPREB1, IGLL1, DNTT, RAG1, RAG2, TCL1A, EBF1, BANK1 |
| T-cell lineage | CD3D, CD3E, CD3G, CD247, TRAC, TRBC2, CD8A, CD8B, CD4, IL7R, CD2, CD5, CD28, CCR7, LCK, GZMK, FOXP3, IL2RA, CTLA4, TIGIT |
| NK cells | NKG7, GNLY, KLRD1, KLRF1, KLRB1, NCAM1, NCR1, FCGR3A, GZMB, PRF1, CX3CR1, TBX21, XCL1, IL18RAP |
| Monocytes | CD14, FCN1, LYZ, S100A8, S100A9, S100A12, VCAN, CST3, CTSS, FCGR3A, MS4A7, CSF1R, ITGAM, CD68, IRF8, CSF3R |
| Macrophages | CD68, CD163, MRC1, MSR1, MERTK, C1QA, C1QB, C1QC, APOE, CD14, CSF1R, MARCO, VSIG4, CD86 |
| Dendritic cells | CLEC9A, XCR1, CADM1, CD1C, FCER1A, CLEC10A, CLEC4C, LILRA4, IL3RA, IRF7, GZMB, TCF4, FLT3, ITGAX, CD83, LAMP3 |
| Erythroid progenitors/erythroblasts | GYPA, HBB, HBA1, HBA2, HBD, ALAS2, AHSP, CA1, KLF1, GATA1, TFRC, SLC4A1, GYPB, HBQ1, RHAG |
| Megakaryocyte lineage | ITGA2B, PF4, PPBP, GP1BA, PLEK, VWF, GP9, ITGB3, TUBB1, MPL |
| Mast cells | CPA3, MS4A2, TPSAB1, TPSB2, KIT, HPGDS, GATA2, CMA1, IL1RL1, SLC18A2 |
| Mesenchymal lineage | PDGFRA, PDGFRB, LEPR, CXCL12, THY1, NT5E, ENG, NGFR, MCAM, VIM, SUSD2, COL1A1, COL1A2, LUM, DCN, SPARC, ISLR, CSPG4, NES, FAP |
| Endothelial cells | PECAM1, CDH5, KDR, VWF, FLT1, PLVAP, EMCN, MCAM, TEK, CLDN5, S1PR1, SELE, ESAM |
| Osteoblasts | RUNX2, SP7, COL1A1, ALPL, BGLAP, SPP1, IBSP, COL1A2, POSTN, MMP13 |
| Osteocytes | SOST, DMP1, MEPE, PHEX, FGF23, BGLAP, ALPL, PDPN, DKK1 |

#### S6G Hyperparameter search space for scVI model optimization

Hyperparameter search space used for scVI model optimization. The search evaluated different latent-space dimensions, numbers of hidden layers, gene expression likelihood distributions, learning rates, and dropout rates.

| Hyperparameter | Tested values |
|---|---|
| Latent dimensions ($n_{latent}$) | 5, 10, 20, 30, 50 |
| Number of hidden layers ($n_{layers}$) | 1, 2, 3 |
| Gene likelihood (gene_likelihood) | Negative binomial (nb), zero-inflated negative binomial (zinb) |
| Learning rate (lr) | 0.0004, 0.001, 0.0001, 0.004 |
| Dropout rate (dropout_rate) | 0, 0.1, 0.2, 0.3 |

#### Supplementary Table S7

Supplementary Table S7. Pooled DEG rankings by effect magnitude and FDR in the pooled analysis. Positive log2FC denotes higher expression in 3D, whereas negative log2FC denotes higher expression in 2D. Four sub-tables show the top 20 genes per direction for each ranking criterion. These tabulated rankings do not apply the expression-prevalence filter displayed in Figure 4B and therefore need not contain exactly the same genes as those dotplots.

##### A. Higher in 3D, ranked by log2FC

| Gene | log2FC | logCPM | FDR |
|---|---|---|---|
| WDR86 | +9.379 | 2.147 | 3.03e-12 |
| MT1F | +8.993 | 2.518 | 4.89e-11 |
| CCL5 | +8.439 | 1.460 | 1.04e-10 |
| TREM1 | +8.035 | 4.996 | 8.35e-13 |
| CPXM2 | +7.833 | 1.300 | 2.39e-11 |
| KCNN3 | +7.673 | 1.141 | 2.28e-10 |
| PCBP3 | +7.615 | 2.355 | 2.60e-10 |
| CADM3 | +7.456 | 1.415 | 4.24e-09 |
| COL5A3 | +7.390 | 5.505 | 5.50e-13 |
| IL1R2 | +7.269 | 0.060 | 4.17e-09 |
| CHRDL2 | +7.156 | -0.361 | 5.51e-08 |
| ENSG00000263655 | +7.142 | 0.435 | 1.29e-09 |
| IFITM1 | +7.060 | 6.315 | 3.37e-13 |
| LINC01411 | +7.053 | 0.352 | 5.53e-05 |
| EREG | +6.882 | 2.170 | 1.45e-09 |

| Gene | log2FC | logCPM | FDR |
|---|---|---|---|
| FLT1 | +6.822 | 1.451 | 2.44e-08 |
| ZNF804A | +6.792 | 2.599 | 1.29e-12 |
| EPB41L3 | +6.752 | 1.630 | 8.04e-11 |
| HES5 | +6.740 | 0.071 | 1.38e-08 |
| CP | +6.705 | 2.318 | 7.14e-12 |

**B. Higher in 2D, ranked by |log2FC|**

| Gene | log2FC | logCPM | FDR |
|---|---|---|---|
| GPNMB | -14.175 | 6.263 | 1.23e-11 |
| TRHDE | -11.143 | 3.476 | 2.16e-08 |
| FABP4 | -9.864 | 2.249 | 5.91e-06 |
| CTAG2 | -9.862 | 2.246 | 2.05e-05 |
| CYSRT1 | -9.265 | 1.673 | 2.47e-05 |
| ZNF560 | -9.133 | 1.546 | 8.93e-05 |
| FOLR3 | -8.584 | 1.023 | 1.78e-05 |
| TGFBR3L | -7.963 | 0.440 | 9.13e-05 |
| ENSG00000287255 | -7.721 | 0.215 | 1.33e-04 |
| LINC02109 | -7.721 | 0.215 | 1.81e-04 |
| SNX10 | -7.611 | 0.115 | 2.33e-04 |
| LINC02621 | -7.336 | -0.134 | 6.30e-04 |
| ODAPH | -7.327 | 3.109 | 9.34e-09 |
| ENPP3 | -7.308 | -0.159 | 1.04e-03 |
| PF4 | -7.308 | -0.159 | 4.57e-04 |
| KCNMA1-AS1 | -7.294 | -0.172 | 1.14e-03 |
| HCST | -7.280 | -0.185 | 8.90e-04 |
| SUGCT-AS1 | -7.221 | -0.237 | 9.79e-04 |
| ENHO | -7.191 | -0.264 | 4.36e-04 |
| ENSG00000287421 | -7.129 | -0.319 | 7.53e-04 |

**C. Higher in 3D, ranked by FDR**

| Gene | log2FC | logCPM | FDR |
|---|---|---|---|
| TNFSF10 | +6.026 | 6.829 | 2.99e-14 |
| ENSG00000289901 | +4.164 | 5.935 | 2.99e-14 |
| PRKG1 | +3.353 | 6.737 | 3.42e-14 |
| MT-CYB | +2.845 | 9.794 | 3.42e-14 |
| ENSG00000289474 | +3.832 | 5.938 | 4.39e-14 |
| GFPT2 | +3.657 | 5.731 | 4.41e-14 |
| GPC6 | +3.198 | 8.380 | 1.36e-13 |
| EMILIN1 | +2.914 | 6.909 | 1.52e-13 |
| VCAM1 | +3.985 | 5.438 | 1.58e-13 |
| DCN | +3.482 | 10.163 | 1.86e-13 |
| MT1E | +2.889 | 8.279 | 1.86e-13 |
| C1R | +3.965 | 6.401 | 2.00e-13 |
| CA12 | +2.938 | 6.064 | 2.16e-13 |
| GJB2 | +5.827 | 4.722 | 2.18e-13 |
| CDH13 | +2.358 | 7.987 | 2.43e-13 |
| PTPRG | +2.369 | 7.530 | 2.47e-13 |
| IFITM1 | +7.060 | 6.315 | 3.37e-13 |
| MT-ND1 | +1.991 | 8.245 | 3.37e-13 |
| MT-ND5 | +1.727 | 8.196 | 3.50e-13 |
| IL1R1 | +3.848 | 5.863 | 3.63e-13 |

**D. Higher in 2D, ranked by FDR**

| Gene | log2FC | logCPM | FDR |
|---|---|---|---|
| KCNMA1 | -6.926 | 9.288 | 2.99e-14 |
| AKAP12 | -3.723 | 8.508 | 2.99e-14 |
| CCND1 | -2.950 | 9.818 | 2.99e-14 |
| PPP1R14B | -2.521 | 10.857 | 2.99e-14 |
| FABP5 | -2.772 | 9.388 | 3.42e-14 |
| SMYD3 | -2.608 | 9.898 | 3.42e-14 |
| ANKRD1 | -4.648 | 8.348 | 4.41e-14 |

| Gene | log2FC | logCPM | FDR |
|---|---|---|---|
| MRPL52 | -2.495 | 8.557 | 1.20e-13 |
| MGST3 | -2.173 | 9.415 | 1.36e-13 |
| PFDN5 | -1.996 | 9.561 | 1.36e-13 |
| NME2 | -1.977 | 11.067 | 1.36e-13 |
| CCDC85B | -2.908 | 7.827 | 1.58e-13 |
| RALA | -2.515 | 8.962 | 1.85e-13 |
| CAV1 | -2.102 | 9.478 | 1.86e-13 |
| HINT1 | -2.325 | 10.276 | 1.95e-13 |
| NDUFAF8 | -2.869 | 8.824 | 2.00e-13 |
| GAS5 | -2.473 | 8.782 | 2.00e-13 |
| CLTA | -2.045 | 9.332 | 2.00e-13 |
| BRI3 | -2.648 | 9.636 | 2.22e-13 |
| CFL1 | -1.794 | 11.676 | 2.43e-13 |

#### Supplementary Table S8

Supplementary Table S8. Representative cluster-versus-rest DEGs and biological descriptions for the six MSC clusters.

| Cluster | Representative positive DEGs | DEG-based biological description / limitation |
|---|---|---|
| 3D C1 (n=245) | GPC6, TNFSF10, DCN, IFITM1, HLA-B, HES1, VCAM1, COL5A3 | ECM/adhesion, immune-response and progenitor markers. Validate immune-signal origin at cell level. |
| 3D C2 (n=60) | HLA-B, DCN, RCN1, SERPINH1, MIF, RPL37A, TREM1, IFITM3, TGFBI | Small mixed stromal/biosynthetic cluster. Particularly sensitive to QC and sampling effects. |
| 2D C1 (n=3,674) | CCND1, NCL, HSP90AA1, CALD1, ANXA2, TAGLN, CAV1, AKAP12 | Broad contractile and biosynthetic culture cluster. Not exclusively proliferative. |
| 2D C2 (n=7,788) | PFN1, TNFRSF12A, THBS1, CCN1/CCN2, KCNMA1, GDF15 | Major matrix/mechanoadapted cluster with pronounced structural markers. |
| 2D C3 (n=1,216) | UBE2S, PCLAF, PTTG1, CDKN3, STMN1, BIRC5, CCNB1, TOP2A | Clearly proliferative 2D subcluster. |
| 2D C4 (n=192) | MALAT1, NEAT1, MIR31HG, HMGA2, RBMS3, PLCB4, KCNQ5 | Small lncRNA-rich cluster. These markers do not establish quiescence or senescence. |

### Supplementary Table S9

Supplementary Table S9. Representative pooled and cluster-resolved GSEA modules at q<0.05.

#### A. Pooled 3D-versus-2D analysis

| Direction | Representative gene set | NES | q value | Leading-edge examples |
|---|---|---|---|---|
| 3D | TNF signalling | +2.13 | 0† | CCL5, TNFRSF1B, CXCL6, SOCS3, ICAM1, CXCL2 |
| 3D | Interferon-gamma response | +2.13 | 0† | CCL5, TNFSF10, CFB, BST2, SOCS3, CD74 |
| 3D | Cytokine–receptor interaction | +2.04 | 0† | CCL5, IL1R2, BMP8B, TNFSF10, IL19, TNFRSF1B |
| 3D | Collagen biosynthesis/modification | +1.96 | 7.55e-03 | COL5A3, COL21A1, COL4A4, COL24A1, COL15A1, COL4A3 |
| 3D | Cell-adhesion molecules | +1.94 | 1.01e-03 | CADM3, HLA-F, ICAM1, HLA-DRA, NCAM1, VCAM1 |
| 3D | G2/M-Checkpoint | +1.63 | 8.20e-03 | SMAD3, RPS6KA5, LMNB1, EFNA5, TOP2A, PLK4 |
| 3D | Hypoxia | +1.44 | 4.82e-02 | CP, SLC2A5, ALDOC, NDRG1, DDIT4, AMPD3 |
| 2D | Actomyosin/contraction programme | -1.87 | 3.47e-02 | TPM2, TPM1, ACTN4, TPM3, DMD, ACTA2 |
| 2D | Translation | -2.73 | 0† | MRPL52, MRPL41, MRPL18, MRPL20, MRPL58, AURKAIP1 |
| 2D | Electron transport chain/OXPHOS system in mitochondria (WikiPathways WP111) | -2.85 | 0† | SLC25A4, NDUFA4, NDUFAB1, ATP5MC3, NDUFS8, COX7A1 |
| 2D | Oxidative phosphorylation | -2.96 | 0† | ATP6V1F, SLC25A4, ATP6V0E1, MGST3, NDUFA4, NDUFAB1 |
| 2D | Mitochondrial translation elongation (Reactome R-HSA-5389840) | -3.02 | 0† | MRPL52, MRPL41, MRPL18, MRPL20, MRPL58, AURKAIP1 |
| 2D | Respiratory electron transport and ATP synthesis (Reactome R-HSA-163200) | −2.78 | 0† | NDUFAF4, NDUFAB1, COX20, ATP5MC3, NDUFS8, COX6A1 |

#### B. Cluster-resolved representative modules

| Cluster | Direction | Module | NES | q value | Leading-edge examples |
|---|---|---|---|---|---|
| 3D C1 | positive | TNF/NF-κB | +1.90 | 0† | NR4A3, CXCL2, AREG, PTGS2, ICAM1, IL1B |
| 3D C1 | positive | Chemokine activity | +2.00 | 9.62e-04 | CXCL2, CCL5, CCL2, CCL7, CCL20, KLF5 |
| 3D C1 | negative | tRNA aminoacylation | -3.18 | 0† | FARSB, RARS1, LARS1, GARS1, EEF1E1, VARS1 |
| 3D C2 | positive | Ribosome | +3.64 | 0† | RPL31, RPS25, RPL30, RPS10, RPS5, RPS27 |
| 3D C2 | positive | Mitochondrial complex I | +3.30 | 0† | NDUFS3, NDUFA11, NDUFA5, NDUFB1, NDUFA13, NDUFB11 |
| 2D C1 | positive | Ribosome | +3.90 | 0† | UBA52, FAU, RPS10, RPL21, MRPL33, RPS20 |

| Cluster | Direction | Module | NES | q value | Leading-edge examples |
|---|---|---|---|---|---|
| 2D C1 | positive | Oxidative phosphorylation | +3.82 | 0† | PHYH, SLC25A4, ATP6V0E1, NDUFA4, ATP6V0C, MGST3 |
| 2D C1 | positive | Peptide-chain elongation | +3.82 | 0† | UBA52, FAU, RPS10, RPL21, RPS20, RPL23A |
| 2D C2 | positive | Oxidative phosphorylation | +1.79 | 0† | SLC25A4, ALAS1, ECI1, SURF1, PHYH, ATP6V0E1 |
| 2D C2 | negative | Chromatid segregation | -4.00 | 0† | TOP2A, NUSAP1, NDC80, KIFC1, CENPI, PLK1 |
| 2D C2 | negative | Mitotic spindle checkpoint | -3.52 | 0† | SPC25, CENPF, NDC80, NUF2, BUB1B, SPC24 |
| 2D C3 | positive | E2F targets | +1.60 | 8.68e-03 | CDCA3, CENPM, CDC20, KIF18B, MYC, GINS3 |
| 2D C3 | positive | G2/M-Checkpoint | +1.64 | 1.40e-02 | NEK2, CCNF, CDC20, MYC, POLQ, RBL1 |
| 2D C3 | positive | Wnt/β-Catenin | +1.62 | 1.00e-02 | DKK1, MYC, FZD8, NOTCH1, JAG1, MAML1 |
| 2D C4 | positive | PIP synthesis at the early endosome | +2.47 | 1.43e-03 | PIKFYVE, INPP4B, PIK3R4, INPP4A, MTM1, PIK3C3 |
| 2D C4 | negative | Respiratory chain/OXPHOS | -2.26 | 0† | NDUFA4, UQCRB, ATP5MC3, COX4I1, UQCRH, ATP5MG |
| 2D C4 | negative | Oxidative phosphorylation | -2.24 | 0† | NDUFA4, UQCRB, ATP5MC3, COX4I1, UQCRH, ATP5MG |

† A value of 0 is the numerical FDR q-value estimate reported by the enrichment software, not an exact zero probability. No validated positive reporting bound is available. It must not be converted to < 0.001 solely by assuming 1,000 permutations.

#### Supplementary Table S10

Supplementary Table S10. Detection of canonical and bone-marrow-associated markers across the six MSC clusters.

| Marker | 3D C1 | 3D C2 | 2D C1 | 2D C2 | 2D C3 | 2D C4 |
|---|---|---|---|---|---|---|
| NT5E/CD73 | 97.1 % | 28.3 % | 15.2 % | 82.3 % | 83.8 % | 22.9 % |
| THY1/CD90 | 97.1 % | 36.7 % | 56.9 % | 87.8 % | 97.8 % | 0.0 % |
| ENG/CD105 | 95.5 % | 16.7 % | 2.6 % | 53.7 % | 50.8 % | 4.7 % |
| NGFR/CD271 | 0.8 % | 0.0 % | 0.0 % | 0.1 % | 0.0 % | 0.0 % |
| LEPR | 25.7 % | 0.0 % | 0.2 % | 5.5 % | 5.4 % | 3.6 % |
| PDGFRB/CD140b | 94.3 % | 1.7 % | 0.2 % | 34.7 % | 28.5 % | 15.1 % |

| Marker | 3D C1 | 3D C2 | 2D C1 | 2D C2 | 2D C3 | 2D C4 |
|---|---|---|---|---|---|---|
| MCAM/CD146 | 44.9 % | 1.7 % | 0.0 % | 15.0 % | 20.9 % | 0.0 % |
| PRRX1 | 97.1 % | 8.3 % | 13.7 % | 83.2 % | 82.6 % | 36.5 % |
| COL1A1 | 100.0 % | 65.0 % | 16.3 % | 99.9 % | 100.0 % | 76.6 % |

#### Supplementary Table S11

Supplementary Table S11. CellPhoneDB pair-count summary and representative directed candidates in 3D day 21.

| Source | Target | Reported pairs | Representative candidate axes | Interpretation |
|---|---|---|---|---|
| 3D C1 | 3D C1 | 438 | ECM, adhesion and autocrine-compatible candidates | Largest within-cluster reported repertoire |
| 3D C1 | Macrophages | 200 | FN1–integrin, APP–CD74, PLAU–PLAUR | Predicted stromal-to-myeloid candidates |
| Macrophages | 3D C1 | 151 | IL1B–IL1 receptor, TNFSF12–TNFRSF12A, ANXA1–FPR | Predicted myeloid-to-stromal candidates |
| 3D C2 | 3D C2 | 6 | Limited reported set | Small cluster (60 cells) |
| 3D C2 | Macrophages | 49 | Expression-compatible candidates | Cluster-size sensitivity required |
| Macrophages | 3D C2 | 8 | Expression-compatible candidates | Cluster-size sensitivity required |

#### Supplementary Table S12

Supplementary Table S12. The six haematopoietic lineage groups shown in Supplementary Figure S8, with top trend-variance genes and interpretation/QC notes.

| Lineage | Top 10 genes by trend variance | Plausibility/QC note |
|---|---|---|
| T-cell lineage | BTG1, SPINK2, HLA-DRA, PDE3B, IL7R, CD74, GIMAP7, CAMK4, CCR7, STK17A | Plausible T-cell programme. Some early genes are shared with other branches. |
| Erythroid lineage | HBB, CA1, BLVRB, HBA1, PRDX2, VIM, S100A6, SERPINB1, HBA2, ATP5IF1 | Coherent erythroid programme including HBB, HBA1/HBA2, CA1 and BLVRB. |
| Mast-cell lineage | CLC, SRGN, PRG2, RPS2, RPS8, RPS18, SLC24A3, RPS3A, RPL13, RPS6 | CLC/PRG2 are plausible. The remaining ranking is strongly influenced by ribosomal genes. |
| Megakaryocyte lineage | PLEK, SPINK2, GP1BB, TMSB4X, CD74, HLA-DRA, H2AZ1, PDLIM1, ZFP36L2, PF4 | Plausible programme including GP1BB, PF4 and additional megakaryocytic genes. |
| Monocyte/macrophage lineage | S100A8, CXCL8, S100A9, ELANE, PRTN3, MPO, NAMPT, RPS18, RPLP0, AZU1 | Strong myeloid/inflammatory variation. Early granulocytic programmes require contextual review. |

| Lineage | Top 10 genes by trend variance | Plausibility/QC note |
|---|---|---|
| pDC lineage | LGALS1, CST3, SPINK2, JCHAIN, SAMHD1, CD74, IRF8, TMSB4X, GYPC, C12orf75 | Partly plausible with JCHAIN/IRF8. Several ubiquitous genes also rank highly. |

#### Supplementary Table S13

Supplementary Table S13. Literature-guided comparison of selected stromal and progenitor-associated genes in the MSC differential-expression results. Each numeric cell reports log2FC followed by FDR. Positive pooled log2FC indicates higher expression in 3D. The separate C1-versus-rest contrast compares 3D C1 with all remaining MSC clusters and must not be equated with the pooled contrast. The pooled threshold is FDR < 0.01 and |log2FC| ≥ 1. Literature context specifies the investigated species or culture system and supports interpretation rather than proving native identity in the present cells. The selection includes concordant, non-concordant and below-threshold genes. It is a targeted, literature-guided comparison rather than an unbiased native-similarity score or a new DEG analysis. FDR values derive from one donor with non-independent cell splits. They do not provide donor-level inference. CXCL12 was not retained in the pooled export and is considered descriptively elsewhere. Marker-detection fractions are reported in Supplementary Table S10.

| Gene | Pooled log2FC / FDR | C1 vs rest log2FC / FDR | Pooled criterion | Literature context |
|---|---|---|---|---|
| VCAM1 | +3.98 1.58e-13 | +4.24 5.47e-32 | Higher in 3D | Human uncultured stromal cells and adhesion [8, 21] |
| DCN | +3.48 1.86e-13 | +3.90 1.08e-39 | Higher in 3D | Human stromal matrix programmes [8, 21] |
| SERPING1 | +3.35 1.85e-12 | +4.14 3.18e-21 | Higher in 3D | Human primary-cell and sphere-culture comparison [2] |
| EBF3 | +1.53 1.14e-08 | +3.79 7.84e-11 | Higher in 3D | Murine CAR/LepR-positive niche stromal cells [22] |
| EGR1 | +1.50 1.1e-09 | +3.15 1.68e-23 | Higher in 3D | Human native programme and possible preparation response [21, 23] |
| FOSB | +2.44 4.61e-10 | +4.03 6.95e-20 | Higher in 3D | Human primary cells and possible preparation response [2, 23] |
| HES1 | +3.72 7.24e-13 | +4.75 3.01e-37 | Higher in 3D | Murine Notch-dependent progenitor maintenance [17] |

| Gene | Pooled log2FC / FDR | C1 vs rest log2FC / FDR | Pooled criterion | Literature context |
|---|---|---|---|---|
| GREM1 | +2.36<br>3.08e-08 | +2.14<br>4.66e-14 | Higher in 3D | Murine osteochondroreticular stem cells [24] |
| GAS1 | +2.06<br>4.72e-10 | +5.46<br>1.06e-16 | Higher in 3D | Murine amplifying descendants of endosteal stem cells [25] |
| LEPR | +0.45<br>0.053 | +2.15<br>0.115 | Combined threshold not met | Human stromal niche populations [8, 21] |
| PDGFRA | +0.85<br>1.03e-08 | +2.70<br>1.82e-18 | Combined threshold not met | Human native MSC populations [21] |
| PDGFRB | +0.89<br>8.42e-09 | +2.16<br>2.06e-15 | Combined threshold not met | Human stromal and mural populations [21] |
| PTGDS | -2.19<br>5.06e-08 | +0.44<br>0.837 | Higher in 2D | Human CXCL12/LEPR-rich stromal population [8] |
| KITLG | -0.17<br>0.194 | +2.13<br>0.0641 | Combined threshold not met | Murine functional HSC niche support [20] |
| OAS2 | +0.88<br>0.0333 | — | Combined threshold not met | Human primary-cell and sphere-culture comparison [2] |
| PPARG | -1.68<br>0.0221 | — | Combined threshold not met | Human primary-cell and sphere-culture comparison [2] |
| TNFAIP6 | +0.03<br>0.469 | +1.63<br>2.41e-14 | Combined threshold not met | Human MSC spheroids and immunomodulation [30] |
